# Deletion of astrocyte intermediate filaments GFAP and Vimentin enhances protein synthesis and prevents early synaptic and cognitive dysfunction in a mouse model of Alzheimer’s disease

**DOI:** 10.64898/2026.03.24.713865

**Authors:** Cristina Boers-Escuder, Mandy S. J. Kater, Michiel J. D. Van Der Zwan, Yvonne Gouwenberg, Remco V. Klaassen, Christiaan F. M. Huffels, Milos Pekny, Elly M. Hol, August B. Smit, Mark H. G. Verheijen

## Abstract

In Alzheimer’s disease (AD) astrocytes become reactive, displaying hypertrophic morphology, increased expression of intermediate filament proteins GFAP and Vimentin and impaired homeostatic support to neurons. However, the contribution of reactive astrocytes to AD progression, particularly the role of cytoskeletal hypertrophy, remains unclear. Here, we investigate whether astrocyte intermediate filaments actively contribute to early AD progression. We show that astrogliosis appears as early as at 3 months in APP/PS1 mice, preceding amyloid-β plaque deposition, and is characterized by a strong upregulation of GFAP and Vimentin. Genetic ablation of GFAP and Vimentin attenuated astrogliosis, as evidenced by the absence of hypertrophy of astrocyte processes and restored expression of glutamine synthetase and other proteins altered in reactive astrocytes in AD. Importantly, GFAP and Vimentin deletion prevented cognitive decline in 4-month old male and female mice, independently of amyloid plaque pathology or microglial reactivity. Mass-spectrometry based proteomics of the dorsal hippocampus revealed a downregulation of synaptic proteins and dysregulation of ribosomal and RNA-binding proteins in APP/PS1 mice, both of which were rescued by GFAP and Vimentin deletion. Using astrocyte-specific CRISPR-Cas9-mediated knockout of GFAP and Vimentin, we further demonstrate translation impairments in AD astrocytes, and that GFAP and Vimentin deletion restores this impaired astrocytic translation. Together, our findings identify intermediate filament proteins GFAP and Vimentin as active regulators of astrocyte protein synthesis, and reveal a previously unrecognized mechanism by which reactive astrocytes contribute to early cognitive dysfunction in AD. This highlights these astrocyte intermediate filaments as promising therapeutic targets to counteract reactive astrocyte-driven cognitive decline in the early stages of Alzheimer’s disease.

## Introduction

Astrocytes are glial cells with a complex morphology that tile the central nervous system (CNS) and perform essential functions for proper brain functioning. They maintain ion homeostasis, recycle neurotransmitters, and provide metabolic support for neurons and protection from oxidative stress^1^. Moreover, astrocytes modulate synaptic transmission by close interactions with synapses through their fine perisynaptic astrocyte processes (PAPs)^2–4^. Under pathological conditions, astrocytes strongly contribute to neuroinflammation, in particular during neurodegenerative disorders^5–7^, by undergoing a morphological, functional and molecular transition to a state known as astrogliosis^8^.

Increasing evidence supports a central role for astrocytes in the pathophysiology of Alzheimer’s Disease (AD). Astrogliosis is observed at early stages of AD, preceeding amyloid plaque deposition and occuring in parallel to synapse loss, which emerges before the start of cognitive symptoms^9,10^. In fact, synapse loss is the best correlate of cognitive decline in AD^11–13^. Moreover, cerebrospinal fluid (CSF) and plasma levels of GFAP, a common marker for reactive astrocytes, are elevated in AD patients and can even be used to predict future converstion to AD in patients with mild cognitive impairment^14,15^. At the structural level, reactive astrocytes display pronounced morphological adaptations, becoming hypertrophic and upregulating their intermediate filament (IF) proteins GFAP and Vimentin^8,16^. Additionally, astrocytes in AD mouse models show diminished morphological complexity and reduced expression of PAP-associated proteins, suggesting that astrocyte reactivity affects the structural and functional interactions between astrocytes and synapses^2,17^. Functionally, astrogliosis can lead to disruptions in neuron-astrocyte signaling, and consequently synaptic dysfuction by impairing glutamate recycling, astrocyte calcium signalling and disregulating gliotransmitter release. These alterations can lead to glutamate excitotoxicity and neurotoxicity through release of saturated fatty acids^1,18,19^. Together, these observations underscore the importance of astrocytes in the early stages of AD pathogenesis, however the molecular mechanisms by which reactive astrocytes contribute to early disease progression are poorly understood.

To investigate whether and how astrogliosis contributes to early AD pathology, before overt reactivity is observed, we interfered with astrogliosis in an APPswe/PSEN1dE9 (APP/PS1) mouse model of AD^20^. One approach to attenuate astrocyte reactivity is the genetic ablation of the IF proteins GFAP and Vimentin^21–24^ (hereafter refered to as GFAP/Vim deletion). Astrocytes in *Gfap^-/-^Vim^-/-^* mice are devoid of IFs, lack hypertrophy of their cellular processes after CNS injury, display attenuated astrogliosis, and support improved neuronal and synaptic regeneration after a brain lesion^22,25–28^. This is in line with the functions of GFAP and Vimentin in stress resistance^29–32^. In the context of AD, *Gfap^-/-^Vim^-/-^* astrocytes in 15-18 month APP/PS1 mice show reduced association with amyloid plaques, accompanied by downregulation of neuroinflammatory genes^23^, but whether this attenuation in astrocyte reactivity contributes to AD progression has not been studied. Remarkably, under physiological conditions, astrocytes of *Gfap^-/-^Vim^-/-^* mice exhibit a morphology and tissue coverage comparable to wildtype (WT) astrocytes^22^. This model therefore allows to study the molecular, morphological and functional consequences of astrogliosis, and how it contributes to early AD pathology. For this goal, we crossed *Gfap^-/-^Vim^-/-^* mice with APP/PS1 mice and confirmed a reduction in astrogliosis. Next, we performed proteomic profiling in young, 4-month old, APP/PS1x*Gfap^-/-^Vim^-/-^*to assess the molecular changes caused by the attenuated astrogliosis at this early disease stage. Mechanistically, we found that, next to the upregulation of GFAP and Vimentin, astrogliosis was associated with impairments in RNA processing, protein translation and a loss of synaptic proteins in the hippocampus. In agreement with this, GFAP/Vim deletion restored astrocytic protein synthesis and prevented early cognitive decline in APP/PS1 mice, independently of amyloid-β (Aβ) plaque pathology or microgliosis. Combined, our results demonstrate a direct role for astrogliosis in early cognitive dysfunction in AD by impairing astrocytic protein synthesis.

## Results

### 1. Astrogliosis appears at 3 months in the hippocampus of APP/PS1 mice

Previously, we described that in the hippocampus of APP/PS1 mice, increased levels of soluble Aβ_42_ coincided with the first appearance of microgliosis at 3 months of age, and precede the presence of Aβ plaques detectable at 4 months^33^. To also detemine the temporal progression of astrogliosis in APP/PS1 mice, we analyzed the hippocampal levels of GFAP and glutamine synthetase (GS) at multiple timepoints between 2 and 6 months of age (Fig. 1A). Immunohistochemical staining performed on hippocampal CA1 revealed a significant increase in GFAP immunoreactivity in APP/PS1 mice, starting at 3 months of age, with further increases at 4 and 6 months of age (Fig. 1B, C). Consistently, immunoblotting analysis of the hippocampus also revealed an elevation of GFAP levels at 3 months of age (Fig. 1D). In addition, analysis of GS by immunohistochemistry (Fig.1C) and immunoblotting (Fig 1D) showed a reduction in GS levels in APP/PS1 mice detectable at respectively 4 and 3 months (Fig. 1C), which is in line with published observations showing decreased GS levels in reactive astrocytes^34^ and in astrocytes with progression of AD^35–38^. Together, these data suggest that Aβ-induced astrogliosis in the hippocampus can be detected as early as at 3 months of age, coinciding with the onset of microgliosi and preceding Aβ plaque deposition.

**Figure 1.**
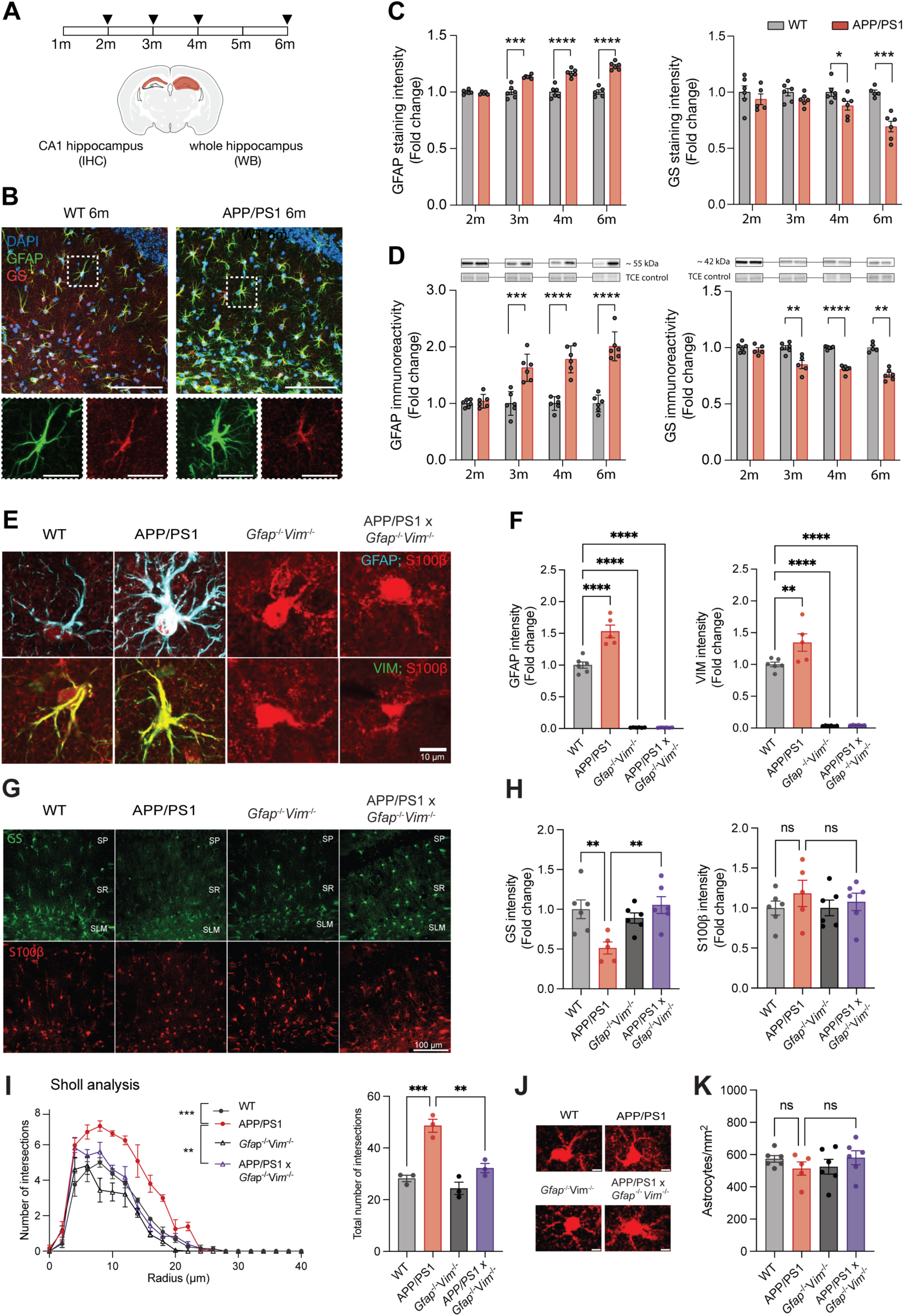
Astrogliosis appears in APP/PS1 mice at 4 months and can be reduced by deletion of GFAP and Vimentin. **A)** Schematic overview of the experiment. Tissue was collected from WT and APP/PS1 mice at 2, 3, 4 and 6 months of age. Immunohistochemical (IHC) analysis was performed on the CA1 hippocampus and immunoblotting (WB) on the whole hippocampus. **B)** Representative images showing astrocytes in the CA1 hippocampus stained for GFAP (green), GS (red), and nuclei stained with DAPI (blue). Top, overview showing multiple astrocytes for WT (left) and APP/PS1 (right) at age 6 months. Scale bar, 100 μm. Bottom, a zoom-in on a single astrocyte. Scale bar, 10 μm. **C)** Fluorescence intensity measured per individual astrocyte for GFAP (left) and GS (right). **D)** Immunoblotting for GFAP (left) and GS (right). Representative sections of corresponding proteins at immunoblot and 2,2,2-trichloroethanol (TCE) as loading control are shown above each condition. Data in C-D represented as fold-change relative to WT per timepoint; n = 5-6 mice per genotype. **E)** Representative images of single astrocytes in the hippocampal CA1 with all astrocytes stained for S100β (red) to appreciate the deletion of GFAP (cyan) and Vimentin (green). Scale bar, 10 μm. **F)** Quantification of fluorescence intensity measured per individual astrocyte for GFAP (left) and Vimentin (right). **G)** Representative images of GS and S100β protein levels in CA1. The SP (*stratum pyramidale),* SR (*stratum radiatum)* and SLM (*stratum lacunosum-moleculare)* are depicted. Scale bar, 100 μm. **H)** Quantification of fluorescence intensity measured per individual astrocyte for GS (left) and S100β (right). Data in F, H presented as fold-change relative to WT. n = 5-6 mice per genotype. **I)** Sholl analysis for CA1 *stratum radiatum* astrocytes based on S100β signal. Left, Sholl analysis profile displaying the mean number of intersections per radius. Right, quantification of the total number of intersections; n = 16-21 astrocytes from 3 mice per genotype. **J)** Representative images of individual astrocytes selected for the Sholl analysis. Scale bar, 5 μm. **K)** Total number of S100β+ astrocytes in CA1 per mm^2^. Statistical significance for C, D calculated by unpaired t-test; for F, H, I (right) and K calculated by one-way ANOVA; and for I (left) by two-way repeated-measures ANOVA. See table S3 for full statistical details. *p<0.05; **p<0.01; **p <0.001; ****p<0.0001. Data are presented as mean ± SEM.

### 2. GFAP/Vim deletion prevents astrogliosis in APP/PS1 mice

To study the mechanisms underlying astrogliosis in the context of AD, we generated APP/PS1 mice lacking both GFAP and Vimentin^20,27^, hereafter referred to as APP/PS1x*Gfap^-/-^Vim^-/-^* (Fig. S1A). Immunohistochemical stainings confirmed that hippocampal CA1 astrocytes from both *Gfap^-/-^Vim^-/-^*and APP/PS1x*Gfap^-/-^Vim^-/-^*mice lacked GFAP and Vimentin proteins (Fig. 1E, F, Fig. S2A), which was further validated by immunoblotting for GFAP (Fig. S2B) and Vim (Fig. S2C). From 2 months of age onward, an increase in mortality, which was accompanied by a decrease in body weight, was observed specifically for male, but not female, *Gfap^-/-^Vim^-/-^*and APP/PS1x*Gfap^-/-^Vim^-/-^* mice independent of AD genotype (Fig. S1B, C). Autopsy revealed an enlarged bladder and kidney abnormalities, suggesting failure of the renal system as cause of death (Fig. S1D), in line with reported essential functions of IFs in renal physiology^39^. Moreover, *Vim^-/-^* mice have been reported to display enhanced lethality due to reduced renal mass^40^. Since this increase in mortality was restricted to male mice, we continued our analysis in female mice to avoid animal suffering and confounding systemic effects.

To validate previously findings that GFAP/Vim deletion prevents astrocytes to become reactive in APP/PS1 mice, we used astrocyte markers GS and S100β. Immunohistochemical staining showed that the reduction in GS protein levels in APP/PS1 mice was restored to WT-level in APP/PS1x*Gfap^-/-^Vim^-/-^*using immunostainings (Fig. 1G, H), which was partially confirmed by immunoblotting (Fig. S2D). For the astrocyte protein S100β, we found a non-significant trend suggestive of upregulation of protein levels in 4-month-old APP/PS1 mice compared to WT (Fig. 1G, H). Because astrocyte reactivity in AD is also characterised by morphological hypertrophy, we assessed astrocyte morphology using Sholl analysis based on S100β immunostaining (Fig. 1I). Since GFAP is more commonly used for Sholl analysis in astrocytes, but absent in APP/PS1x*Gfap^-/-^Vim^-/-^* mice, we first confirmed the use of S100β for Sholl analyisis by comparing the results using GFAP or S100β on the same astrocytes in WT and APP/PS1 mice and observed no significant differences (Fig. S2E, F), indicating that S100β is indeed a suitable marker for astrocytic morphological assessment. Using this approach, we confirmed astrocyte hypertrophy in APP/PS1 mice with both GFAP- and S100β-based analysis (Fig. 1I, J, Fig. S2G), which was reduced by GFAP/Vim deletion (Fig. 2I). Of note, astrocytes stained positive for S100β in all conditions (Fig. 1J, Fig. S2E), and the density of S100β+ cells in the hippocampus of these mice was not affected by GFAP/Vim deletion, indicating that astrocyte viability was preserved (Fig. 1K). Based on our Sholl analysis, under homeostatic conditions *Gfap^-/-^Vim^-/-^* astrocytes displayed a morphology that is indistinguishable from that of WT astrocytes, in line with astrocyte domains occupying comparable tissue volumes^22^. Furthermore, electron microscopy analysis of PAP-synapse interaction in the hippocampus of 4-month animals confirmed a normal structure of *Gfap^-/-^Vim^-/-^* astrocytes by showing that they closely ensheath synapses similar to WT mice (Fig. S2H). These findings indicate that fine astrocytic processes are preserved despite disruption of the IF cytoskeleton by GFAP/Vim deletion. Overall, these results indicate that GFAP/Vim deletion effectively attenuates astrogliosis in early stage AD without compromising astrocyte viability.

**Figure 2.**
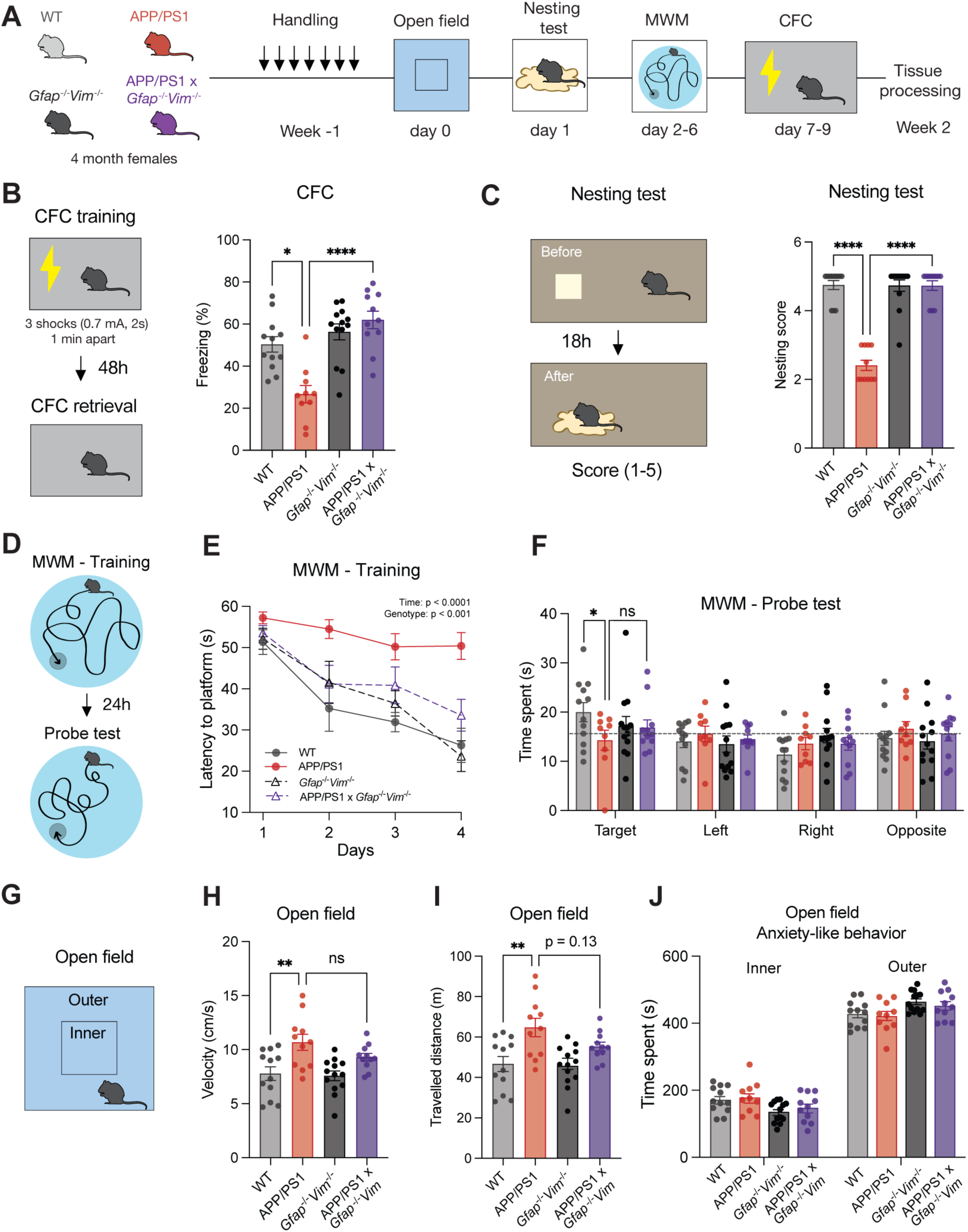
Reducing astrogliosis rescues cognitive decline in 4-month-old APP/PS1 female mice. **A)** Schematic overview of the behavioral experiments. **B)** Left, experimental scheme of the contextual fear conditioning (CFC) training and retrieval. Right, freezing levels (%) measured during the CFC retrieval session 48 hours after shock. **C)** Left, experimental diagram of the nesting test. Right, nest-building scores assessed with the nesting test. **D)** Experimental scheme of the Morris water maze (MWM) training and probe test. **E)** MWM training latencies (in seconds) to reach the platform measured over four training days. **F)** Time spent (in seconds) in the MWM quadrants during the probe test, where the target quadrant refers to the original location of the platform. **G)** Representative image of the open field area and the inner and outer areas. **H, I)** Locomotor activity determined by the velocity (H) and distance travelled (I) during the open field experiment, irrespective of the zone. **J)** Time spent (in seconds) in the inner zone and outer area of the open field experiment. Kruskal-Wallis (B, C), Repeated measures ANOVA (I), One-way ANOVA (E-G, J). n = 9-13 mice per genotype. See table S3 for full statistical details. *p<0.05, **p<0.01, ****p<0.0001, ns: not significant. Data are presented as mean ± SEM.

### 3. GFAP/Vim deletion rescues early cognitive impairment of APP/PS1 mice

To determine whether attenuation of reactive astrogliosis in APP/PS1 mice by GFAP/Vim deletion prevents early cognitive decline, we subjected 4-month-old WT, APP/PS1, *Gfap^-/-^Vim^-/-^*and APP/PS1x*Gfap^-/-^Vim^-/-^* mice to a battery of behavioral tests (Fig. 2A). This age precedes Aβ plaque deposition^33,^ but corresponds to a stage at which young APP/PS1 mice already show measurable cognitive deficits^33,41,42^. Behavioral tests included contextual fear conditioning (CFC), nesting test and Morris mater maze (MWM) test. In the CFC task APP/PS1 mice showed reduced freezing behavior compared to WT mice (Fig. 2B). Remarkably, freezing levels in APP/PS1x*Gfap^-/-^Vim^-/-^*mice were restored to WT-levels (Fig. 2B). Similarly, APP/PS1 mice showed impaired nest-building behavior, with lower nesting scores than WT mice (Fig. 2C), which returned to WT-levels in APP/PS1x*Gfap^-/-^Vim^-/-^* mice (Fig. 2C). Next, we assessed spatial learning and memory by performing the MWM test (Fig. 2D). During the training phase, we observed that APP/PS1x*Gfap^-/-^Vim^-/-^* mice demonstrated improved spatial learning than APP/PS1 mice (Fig. 2E). During the probe trial, APP/PS1x*Gfap^-/-^Vim^-/-^* mice showed a similar preference for the target quadrant as *Gfap^-/-^Vim^-/-^* mice (Fig. 2F). No differences in swimming velocity were observed between genotypes (Fig. S3A). To evaluate general locomotor activity and anxiety-like behavior mice were exposed to a novel environment in the open field test (Fig. 2G). APP/PS1 mice exhibited enhanced locomotor activity relative to WT (Fig. 2H, I) in line with previous observations^33,43^. In contrast, APP/PS1x*Gfap^-/-^Vim^-/-^*mice did not show significantly increased locomotor activity (Fig. 2H, I). Neither, APP/PS1 nor APP/PS1x*Gfap^-/-^Vim^-/-^* mice did show preference for inner versus outer areas of the open field area, indicating no changes in anxiety-like behavior (Fig. 2J). Together, these results indicate that preventing reactive astrogliosis by GFAP/Vim deletion rescues learning and memory deficits in 4-month APP/PS1 mice.

We next asked whether reduction of astrogliosis would be sufficient to rescue cognitive decline at a later AD stage, when plaque deposition is ongoing^33^. To address this, and to rule out sex-specific effects of GFAP/Vim deletion in APP/PS1 mice, we examined cognitive performance in the 6-month old male mice that survived despite increased mortality associated with GFAP/Vimentin deletion (Fig. S1B-D, S4A). Similar to female mice, male APP/PS1x*Gfap^-/-^Vim^-/-^* mice displayed improved CFC performance, nesting behavior and MWM training latencies (Fig. S4B-E). Moreover, while APP/PS1 mice spent less time in the MWM target quadrant than WT mice, APP/PS1x*Gfap^-/-^Vim^-/-^* mice spent a similar amount of time as WT mice (Fig. S4F). Swimming velocity of 6-month-old male mice did not differ between genotypes (Fig. S3B). In the open field test, we did not find any differences in locomotion activity for 6-month-old male APP/PS1 mice (Fig. S4G-I). However, unlike the phenotype observed in 4-month-old females, 6-month-old APP/PS1 male mice showed a significant increase of anxiety-like behavior that was normalized in APP/PS1x*Gfap^-/-^Vim^-/-^* mice (Fig. S4J). In summary, both male and female APP/PS1x*Gfap^-/-^Vim^-/-^* mice show a persistent rescue of AD-associated cognitive deficits, indicating that attenuation of reactive astrogliosis through GFAP/Vim deletion is sufficient to prevent early memory impairments, even at a disease stage when plaque deposition is present^33,42,44^.

### 4. Attenuation of astrogliosis does not affect early A**β** pathology

In AD, reactive astrocytes are often found surrounding Aβ plaques, and astrocytes have been shown to phagocytose Aβ^45^. Accordingly, the attenuation of astrocyte reactivity was reported to increase Aβ plaque load in certain AD mouse models^46^, although other studies found no change^23,47^ or even decreased Aβ plaques^48^. All of these studies investigated late stage AD, whereas the role of reactive astrocytes at the onset of Aβ plaque deposition has not been studied. In light of this, we quantified Aβ plaques (Fig. 3A) and soluble Aβ_42_ levels in 4-month-old female mice. As expected, soluble Aβ_42_ levels were strongly increased in APP/PS1 mice with similar levels in APP/PS1x*Gfap^-/-^Vim^-/-^* mice (Fig. 3B). When examining the accumulation of Aβ plaques at 4-months, we detected small-sized plaques both in the cortex and hippocampus (Fig. 3C-F), likely reflecting newly-formed deposits. Similarly to soluble Aβ levels, plaque density did not differ between APP/PS1x*Gfap^-/-^Vim^-/-^* and APP/PS1 mice (Fig. 3C, D), indicating that astrogliosis does not affect Aβ deposition at early AD stages. Similarly, for 6-month-old males, no significant differences in plaque density were found between APP/PS1x*Gfap^-/-^Vim^-/-^* mice and APP/PS1 mice (Fig. S5A-D). Together, these findings show that GFAP/Vim deletion does not affect soluble Aβ levels or plaque deposition during the early stages of AD.

**Figure 3.**
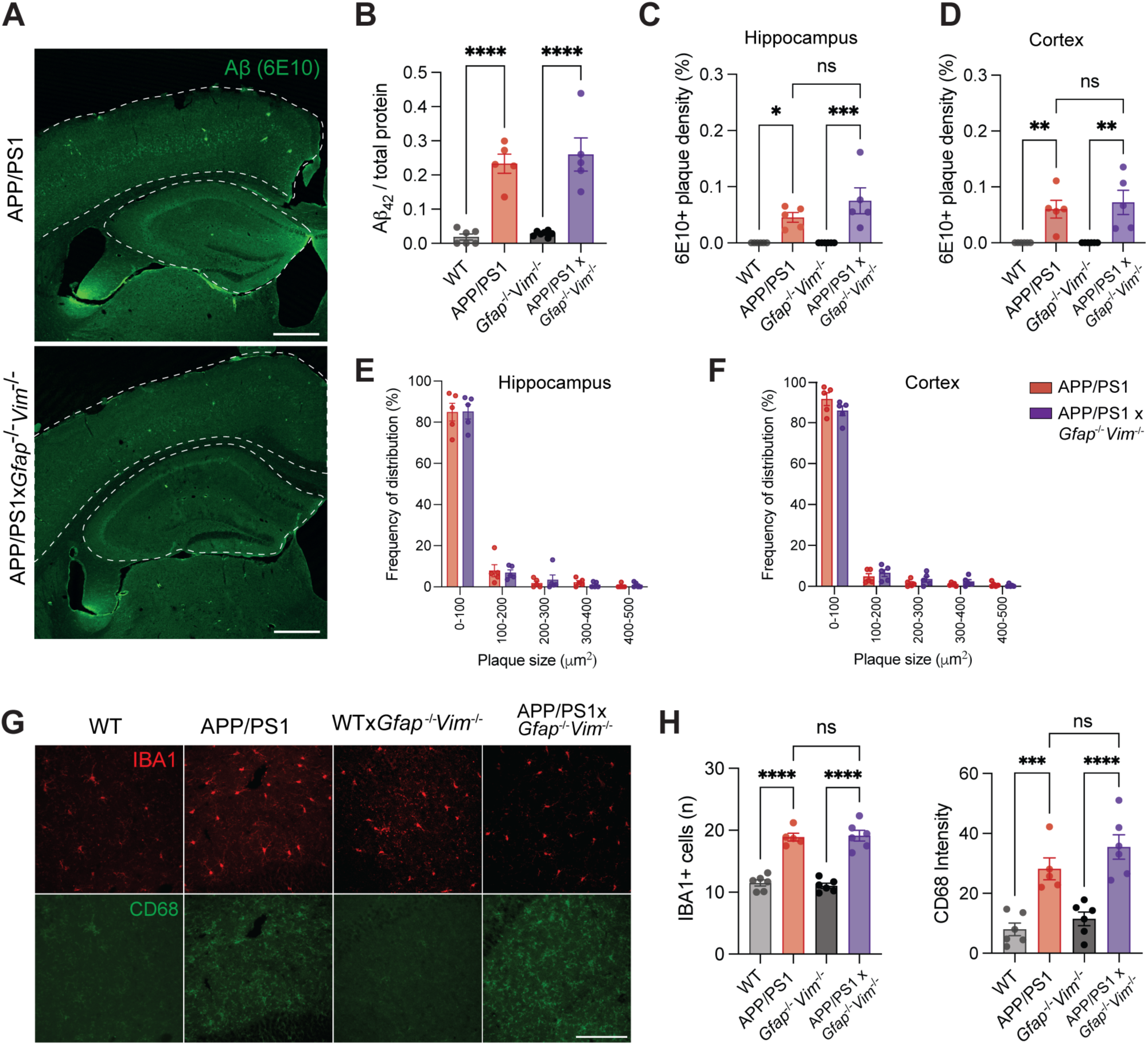
Deletion of GFAP and Vimentin does not affect Aβ pathology or microgliosis. **A)** Representative images of Aβ immunohistochemistry. The dashed lines indicate boundaries of either the cortex or hippocampus. Scale bar, 500 μm. **B)** Levels of soluble Aβ42 measured by ELISA in hippocampus homogenate. **C, D)** Quantification of plaque density in hippocampus (C) and cortex (D), measured as the total Aβ+ plaque area relative to total area. **E, F)** Frequency distribution analysis based on plaque size in hippocampus (E) and cortex (F). **G)** Representative images of CA1 microglia expressing IBA1 (red) and CD68 (green). Scale bar, 100 μm. **H)** Left, quantification of the number of microglia (IBA1+ cells) per field of view (0.5 mm^2^). Right, quantification of mean CD68 intensity per field of view. Statistical analysis performed with one-way ANOVA (B, C, D, H) or two-way ANOVA (E, F); n = 5-7 mice per genotype. See table S3 for full statistical details. *p<0.05, ***p<0.001, ****p<0.0001, ns: not significant. Data are presented as mean ± SEM.

Because the observed astrogliosis (Fig 1) coincides with previously reported microgliosis in APP/PS1 mice^33^ and pharmacological inhibition of microgliosis was shown to rescue early cognitive deficits in APP/PS1 mice^33^, we determined whether GFAP/Vim deletion affects microgliosis. The number of IBA1-positive microglia was unaltered in *Gfap^-/-^Vim^-/-^* mice compared to WT mice. In contrast, we found increased microglial numbers in APP/PS1 mice compared to WT, as previously reported^33^. Importantly, microglial numbers remained elevated in APP/PS1x*Gfap^-/-^Vim^-/-^,* indicating that GFAP/Vim deletion did not affect AD-dependent microgliosis (Fig. 3G, H). Similarly, the levels of CD68, a marker for microglial reactivity, were unaffected in *Gfap^-/-^Vim^-/-^* mice compared to WT, while increased in both APP/PS1 and APP/PS1x*Gfap^-/-^Vim^-/-^* mice (Fig. 3G, H). In summary, the absence of GFAP and Vimentin proteins in astrocytes selectively attenuates astrogliosis without affecting Aβ plaque deposition or microgliosis in early AD stages, enabling us to dissect the astrocyte-specific contributions to AD. Collectivelly, these results demonstrate a role for astrogliosis in early cognitive impairment in AD, independent of amyloid deposition or microglial activation.

### 5. Hippocampal proteomics reveals a rescue of pathways related to synaptic function and protein synthesis in APP/PS1x*Gfap^-/-^Vim^-/-^* mice

Transcriptomic studies have underscored the heterogeneity of astrocyte subtypes in different CNS disorders, including AD, supporting the notion that astrocyte reactivity is a complex dynamic and context-dependent response shaped by the type of disorder, brain region and disease stage^8,49–54^. However, transcriptomic profiles do not necessarily reflect protein abundance, especially given the importance of posttranslational modifications and subcellular localization for protein function^55^. Proteomic studies, combining analysis of whole tissue and subcellular fractions, provide a more direct link to astrocyte-associated functions. To investigate how astrogliosis affects the hippocampal proteome in AD, we performed data-independent acquisition (DIA) micro liquid chromatography coupled to tandem mass spectrometry (LC–MS/MS) on homogenates from the dorsal hippocampus from 4-month-old female WT, APP/PS1, *Gfap^-/-^Vim^-/-^* and APP/PS1x*Gfap^-/-^Vim^-/-^*mice (Fig. 4A). All the samples passed quality control criteria and showed low coefficient of variation in protein abundances (ranging between 0.11 and 0.122 for hippocampal homogenate samples, (Fig. S6A)) and consistent abundance distributions and retention times, indicating high reproducibility between samples (Methods S1). We detected more than 64.000 unique peptides per sample corresponding to more than 8.000 protein groups confidently identified in a minimum of 3 samples per group (Fig. S6A). Differential abundance analysis using MS-DAP^56^ (methods S1) revealed numerous differentially expressed proteins (DEPs; p<0.05) between the different groups, with partial overlap between contrasts (Fig. 4B). A list of all proteins detected in hippocampal homogenates is provided in Table S1. Comparison of WT and APP/PS1 hippocampi revealed extensive proteomic changes with 180 proteins significantly upregulated and 326 downregulated in APP/PS1 (p<0.05). Among the upregulated were well-established AD-associated proteins such as APP, PSEN1 and APOE, and downregulated proteins included many synaptic proteins (Fig. 4C; Fig. S7A-B). In line with prior studies reporting synaptic loss in early AD^33,57–59^, gene set enrichment analysis (GSEA) of DEPs in WT vs APP/PS1 showed a downregulation of synaptic terms, including the postsynaptic structure, synaptic signaling and the regulation of membrane potential (Fig. 4D). SynGO-based overrepresentation analysis of WT vs APP/PS1 DEPs further emphasized the depletion of synaptic proteins in AD hippocampi (Fig. 4E). In addition, we found a downregulation of pathways related to protein synthesis, including mRNA processing and splicing, and upregulation of pathways associated wth in protein degradation (Fig. 4D, Table S2).

**Figure 4.**
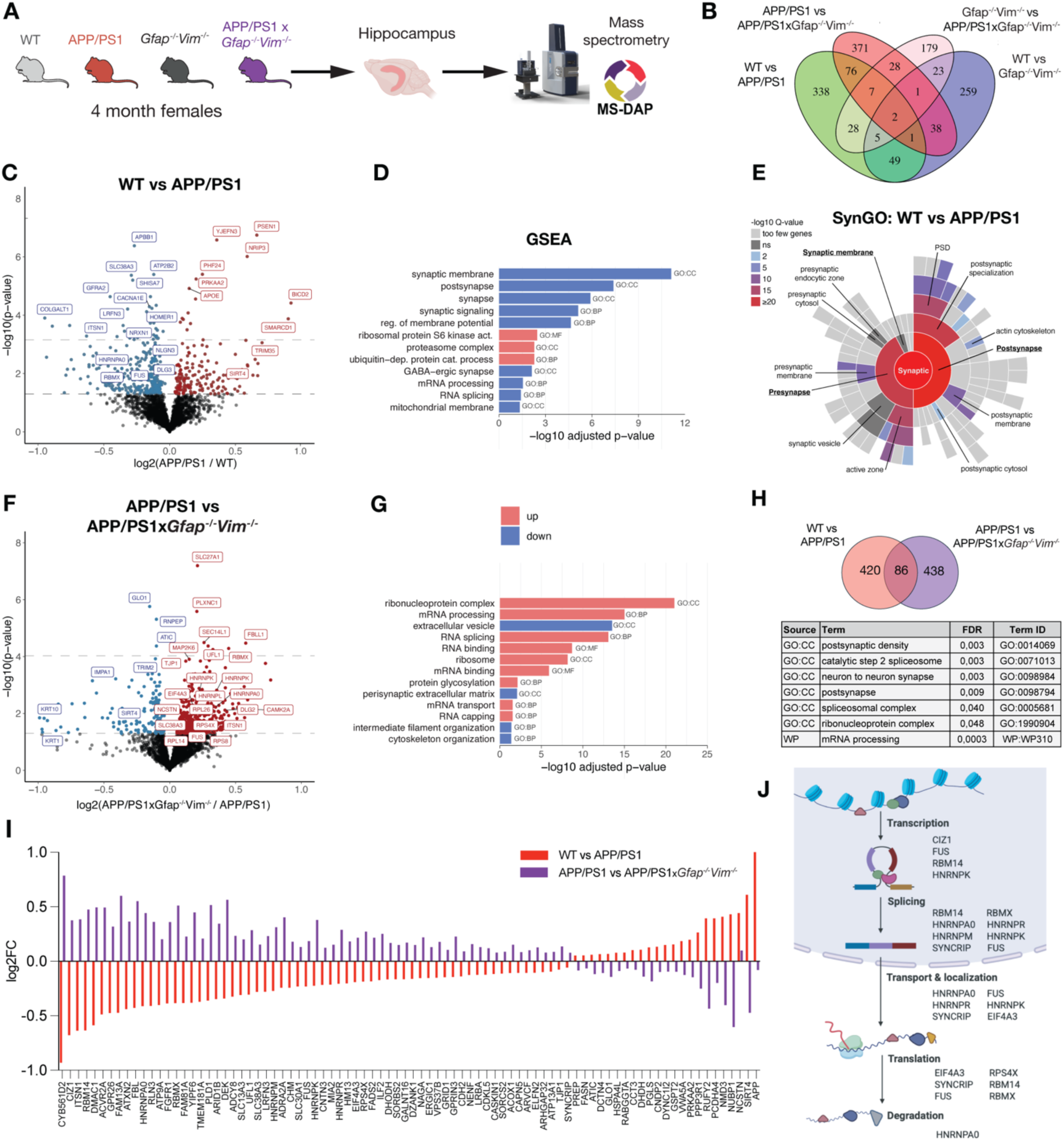
Reduction of astrogliosis rescues hippocampal protein synthesis. **A)** Workflow of hippocampal proteomics analysis. After hippocampal dissection, we performed DIA mass-spectrometry followed by differential expression analysis with MS-DAP. **B)** Venn diagram showing overlapping differentially expressed proteins (DEPs, p<0.05) between the different contrasts. **C)** Volcano plots showing differential expression of proteins in WT vs APP/PS1 (WT: n = 6 mice; APP/PS1: n = 5 mice). Lower dotted line at p<0.05; Upper dotted line at FDR<0.05. For visualization purposes, the y-axis shows values below 8; the x-axis values between -1 and +1. Complete volcano plots are shown in FigS4. Complete DEP list in Table S1. DEPs upregulated in APP/PS1 (red), downregulated DEPs (blue). **D)** Overview of enriched pathways in downregulated (blue) or upregulated (red) DEPs in WT vs APP/PS1 based on gene-set enrichment analysis (GSEA) for gene ontology (GO) terms as indicated (CC, cellular component; BP, biological process; MF, molecular function). Adjusted p-values are FDR-corrected. **E)** Overrepresentation analysis of DEPs in WT vs APP/PS1 using SynGO. **F)** Same as in (C), but for APP/PS1 vs APP/PS1xGfap^/-/^Vim^-/-^ (APP/PS1, n = 5 mice; APP/PS1xGfap^/-/^Vim^-/-^, n = 5 mice). **G)** Same as in (D), but for APP/PS1 vs APP/PS1xGfap^/-/^Vim^-/-^. **H)** Enriched terms based on GO analysis of the 86 shared DEPs between WT vs APP/PS1 and APP/PS1 vs APP/PS1xGfap^/-/^Vim^-/-^ contrasts. **I)** Shared DEPs between WT vs APP/PS1 and APP/PS1 vs APP/PS1xGfap^/-/^Vim^-/-^ ordered by fold-change in WT vs APP/PS1. Note the opposite directions of change for 85 out of the 86 shared DEPs. **J)** Scheme indicating the different stages of protein synthesis at which the shared DEPs are involved. See table S3 for full statistical details. Created in BioRender.

Notably, a part of these AD-driven changes were reversed in APP/PS1x*Gfap^-/-^Vim^-/-^,* in which 350 proteins were upregulated and 174 were downregulated compared to APP/PS1 (Fig. 4F), suggesting that reactive astrocytes substantially contribute to AD-related proteomic changes. Consistent with this interpretation, comparison of *Gfap^-/-^Vim^-/-^* vs APP/PS1x*Gfap^-/-^Vim^-/-^* revealed a total of 273 DEPs, almost half the number observed for WT vs APP/PS1 (Fig. 4B). As expected, GSEA revealed a downregulation of terms for IF, cytoskeleton organization and extracellular vesicles, the latter largely driven by IF keratins known to be in extracellular vesicles^60^. This is consistent with the fact that IFs in astrocytes play a role in vesicle motility^61–63^. Moreover, DEPs upregulated in APP/PS1 vs APP/PS1x*Gfap^-/-^Vim^-/-^* were significantly enriched for protein synthesis-related terms, such as translation, ribosomal components, mRNA transport and splicing (Fig. 4G, table S2). These findings point to a restoration of translational capacity upon reduction of astrogliosis. To substantiate this idea, we selected significant DEPs in both WT vs APP/PS1 and APP/PS1 vs APP/PS1x*Gfap^-/-^Vim^-/-^*, which resulted in 86 shared DEPs, approximately 20% of all DEPs in WT vs APP/PS1 (Fig. 4H). Strikingly, 85 out of these 86 proteins showed a change in the opposite direction in APP/PS1x*Gfap^-/-^Vim^-/-^* compared with APP/PS1 mice, indicating a robust rescue of AD-associated expression changes upon suppression of astrocyte reactivity (R=-0.91, p<2.2e-16, Fig. 4I, Fig. S7E). GO enrichment analysis of these shared DEPs identified biological processes related to mRNA processing (Fig. 4H). Many of these proteins were RNA binding proteins (RBPs) involved in various stages of protein synthesis, including transcriptional regulation and splicing (FUS, hnRNPK), mRNA transport (hnRNPR, hnRNPA0, SYNCRIP), protein translation and ribosome function (eIF4A3, FUS, RPS4X, RBMX, RBM14) (Fig, 4J). A focused analysis of proteins annotated under the GO term “translation” (GO:0006412)^64^ revealed many hnRNPs, ribosomal proteins and proteins involved in mitochondrial translation among the top downregulated proteins in APP/PS1 mice. Many of these were upregulated in APP/PS1x*Gfap^-/-^Vim^-/-^*, especially hnRNPs and cytosolic ribosomal proteins, whereas proteins involved in mitochondrial translation remained mostly downregulated in WT vs APP/PS1x*Gfap^-/-^Vim^-/-^*(Fig. S10). This suggests that GFAP/Vim deletion rescues the expression of proteins involved in cytosolic translation, but not or to a lesser extent mitochondrial translational machinery (Fig. S10).

In addition, the shared DEPs were enriched for synaptic proteins, specifically for postsynaptic components (Fig. 4H), which was confirmed using the SynGO expert-curated database for synaptic proteins (Fig. S7C, D). This suggests that GFAP/Vim deletion partially rescues early synapse loss in AD. In line with this interpretation, approximately 500 proteins were downregulated between WT vs APP/PS1 hippocampi, while only about 250 were downregulated between *Gfap^-/-^Vim^-/-^* and APP/PS1x*Gfap^-/-^Vim^-/-^* (Fig. 4B). Accordingly, DEPs identified between *Gfap^-/-^Vim^-/-^* and APP/PS1x*Gfap^-/-^Vim^-/-^* did not show synaptic enrichment, whereas mitochondrial pathways were upregulated and signal transduction and cytoskeleton pathways were downregulated (Fig. S7G), indicating that certain aspects of AD pathology, for instance the mitochondrial phenotype, are independent of astrogliosis. Together, these findings show that reactive astrocytes in early AD are associated with reduced protein synthesis and synaptic protein levels in the hippocampus, and that genetically reducing astrogliosis partly rescues the AD-associated proteomic profile, in particular protein translational homeostasis and synaptic protein expression.

### 6. Effect of preventing astrogliosis on the synaptic proteome

To accurately determine synapse-specific proteomic alterations in AD, we isolated synaptosomes from the dorsal hippocampus of WT, APP/PS1, *Gfap^-/-^Vim^-/-^* and APP/PS1x*Gfap^-/-^Vim^-/-^* mice, and subjected them to DIA LC-MS/MS proteomic analysis (Fig. 5A). All samples met quality control criteria with median coefficients of variation between 0.116 and 0.13 (Fig. S6A, Methods S1). As for hippocampal homogenates, we detected more than 64.000 unique peptides per sample corresponding to more than 8.000 proteins (Fig. S6A). We confirmed a strong synaptic protein enrichment in synaptosomes compared to hippocampal homogenates through GSEA and SynGO enrichment analysis (Fig. S6C-E, table S2). Differential expression analysis using MS-DAP identified 399 DEPs in WT vs APP/PS1 and 690 DEPs in APP/PS1 vs APP/PS1x*Gfap^-/-^Vim^-/-^*, with 91 proteins overlapping between both contrasts (Fig. 5B-C). Importantly, these expression changes were almost fully reversed by reducing astrogliosis (Fig. 5D), mirroring the reversal of proteomic changes observed in homogenates (Fig 4I). Several proteins rescued in synaptosomes were also restored at the tissue homogenate level, including hnRNPK, an RBP involved in mRNA processing^65^. A complete list of all proteins detected in hippocampal synaptosomes is provided in Table S1.

**Figure 5.**
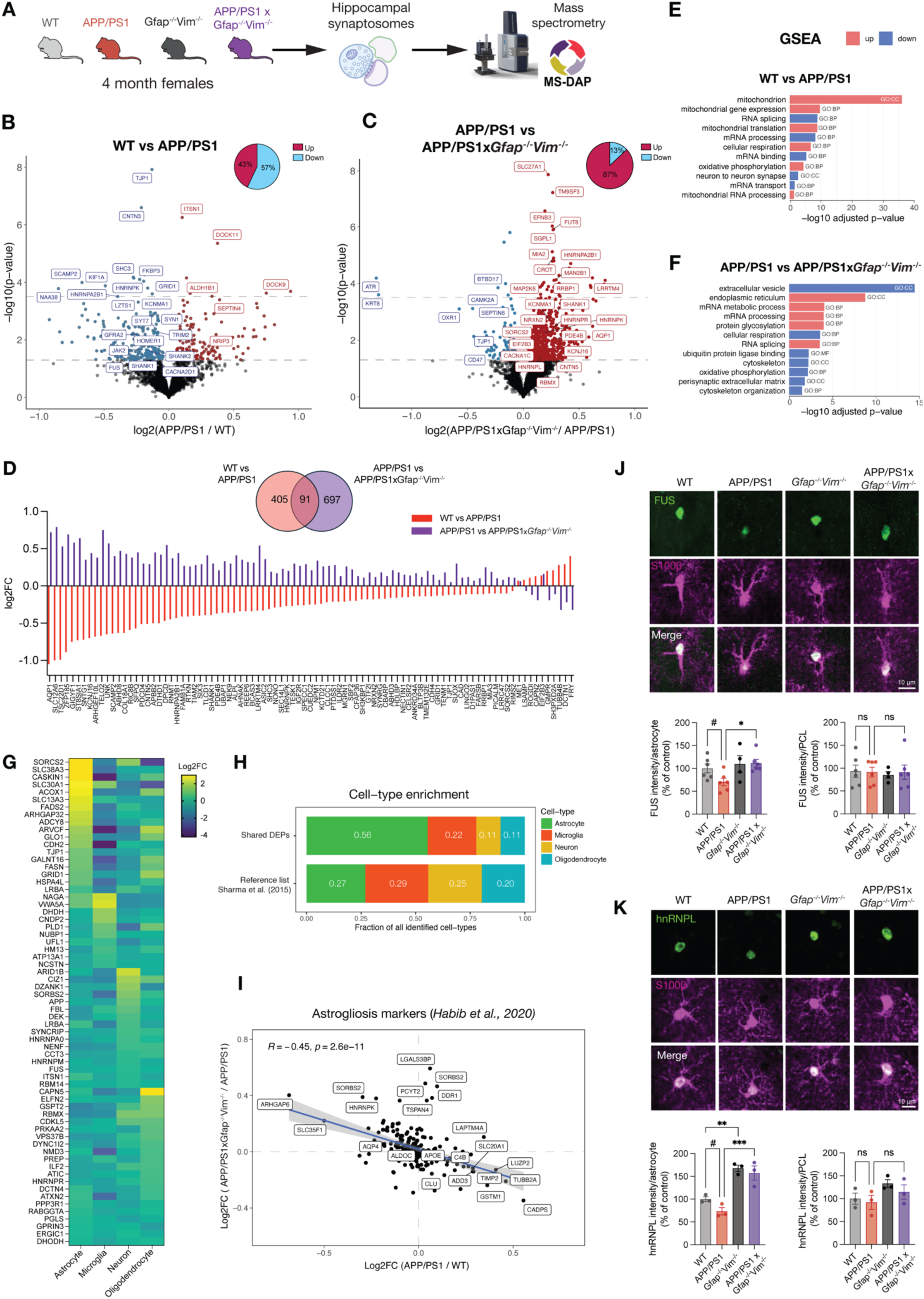
Reduction of astrogliosis rescues altered expression of synaptic and astrocytic proteins in APP/PS1 mice. **A)** Workflow of synaptosome proteomics analysis. After hippocampal dissection we isolated synaptosomes and performed DIA mass-spectrometry followed by differential expression analysis with MS-DAP. **B, C)** Volcano plots showing differential expression of proteins in synaptosomes of WT vs APP/PS1 (B, WT: n = 6 mice; APP/PS1: n = 5 mice) or APP/PS1 vs APP/PS1xGfap^/-/^Vim^-/-^ (C, APP/PS1: n = 5 mice; APP/PS1xGfap^/-/^Vim^-/-^: n = 5 mice). Lower dotted line at p<0.05; Upper dotted line at FDR<0.05. Complete DEP list in Table S1. DEPs upregulated in APP/PS1 (red) and downregulated DEPs (blue). Inset shows percentage of downregulated and upregulated proteins for each contrast. **D)** Shared DEPs between WT vs APP/PS1 and APP/PS1 vs APP/PS1xGfap/-/Vim-/- in hippocampal synaptosomes ordered by fold-change in WT vs APP/PS1. **E, F)** Overview of enriched pathways in downregulated (blue) or upregulated (red) DEPs in synaptosomes of WT vs APP/PS1 (E) or APP/PS1 vs APP/PS1xGfap^/-/^Vim^-/-^ (F) based on GSEA for GO terms (CC, cellular component; BP, biological process; MF, molecular function). Adjusted p-values are FDR-corrected. **G)** Heatmap showing expression of the hippocampal shared DEPs in isolated cell types based on cell-type specific proteomics data by Sharma et al. (2015). **H)** Cell-type enrichment analysis showing the fraction of annotated cell types for shared DEPs in hippocampal homogenates between WT vs APP/PS1 and APP/PS1 vs APP/PS1xGfap^/-/^Vim^-/-^ compared with fractions of cell types identified in the reference list by Sharma et al. (2015). **I)** Correlation plot showing an inverse correlation for markers of reactive astrocytes based on Habib et al. (2020) in hippocampal homogenates of WT vs APP/PS1 compared with APP/PS1 vs APP/PS1xGfap^/-/^Vim^-/-^. **J)** Top, representative images of FUS expression (green) in individual astrocytes (S100β, magenta). Scale bar, 10 μm. Bottom, normalized FUS intensity measured per individual astrocyte (left) or in the hippocampal CA1 pyramidal cell layer (right, see Fig. S8 for images). n = 4-6 mice per genotype. **K)** Same as in (J), but for hnRNPL. Statistical significance calculated by one-way ANOVA (J, K). See table S3 for full statistical details. *p<0.05, #p<0.1, ns: not significant. Data are presented as mean ± SEM.

Because synaptosomal protein amounts are normailzed during preparation, GSEA analysis of synaptosomal protein changes did not show a global loss in synaptic proteins in APP/PS1 mice (Fig. 5E, table S2), enabling a more precise identification of synapse-specific molecular alterations. GSEA analysis revealed an increase in mitochondrial proteins and oxidative phosphorylation in synapses from APP/PS1 mice, confirming our previously reported observations^66^. In parallel, we found a decrease in mRNA processing and other RNA-related processes, which is in line with earlier findings of protein synthesis reduction at AD synapses^67^. Remarkably, many of these pathways were upregulated in APP/PS1x*Gfap^-/-^Vim^-/-^*relative to AD (Fig. 5C, F), suggesting a functional rescue of oxidative phosphorylation and protein synthesis. This restoration was reflected by an upregulation of 87% of the DEPs in the APP/PS1 vs APP/PS1x*Gfap^-/-^Vim^-/-^* comparison. In addition, we observed a dysregulation of other pathways, such as those related to the cytoskeleton, most likely driven by the deletion of IF proteins, and those related to perisynaptic extracellular matrix, extracellular vesicle, and protein glycosylation and ubiquitination (Fig. 5F). Taken together, these results further support that reducing astrogliosis, through GFAP/Vim deletion, restores altered protein synthesis at synapses of APP/PS1 mice and implicates reactive astrocytes to have a central role in early synaptic dysfunction in AD.

### 7. Dysregulated astrocyte proteins in early AD are rescued by GFAP/Vim deletion

To determine the cellular origin of the rescued proteomic alterations (Fig 4I), we performed a cell type enrichment analysis using cell type-specific proteomic data from Sharma et al.^68^. This analysis revealed a significant overrepresentation of astrocytic proteins among the shared DEPs (p=2.78e^-06^, Fig. 5G, H), whereas no enrichment was observed for other cell types. Consisitent with this, we observed an inverse correlation in the expression of disease-associated astrocyte markers in AD^7^ between WT vs APP/PS1 and APP/PS1 vs APP/PS1x*Gfap^-/-^Vim^-/-^* (R=-0.45, p=2.6e^-11^, Fig. 5H), indicating that reactive astrocyte signatures are suppressed upon GFAP/Vim deletion.

Because the rescued protein synthesis in synaptosomes may reflect not only neuronal synaptic proteins but also contributions from astrocytic leaflets or PAPs that closely ensheathe synapses, we examined two published datasets of PAP-enriched transcripts, reflecting local translation in the PAPs^69,70^ (Fig. S8A), and observed a significant inverse correlation between APP/PS1 and APP/PS1x*Gfap^-/-^Vim^-/-^* for both datasets (Sakers et al., 2017^69^: R=-0.49, p=1e^-13^, Fig. S8B; Mazare et al^70^., 2020: R=-0.73, p=4.5e^-10^, Fig. S8C). Similarly, analysis of PAP-enriched proteins identified through proximity-labeling of the PAP protein ezrin^55^ revealed an inverse expression pattern in APP/PS1 mice vs APP/PS1x*Gfap^-/-^Vim^-/-^*mice (Soto et al., 2023^55^: R=-0.5, p=1.4e^-13^, Fig. S8D). These analyses indicate that the expression of locally-translated PAP proteins is restored upon GFAP/Vim deletion in APP/PS1 mice.

These findings suggest that a part of the observed proteomic rescue originates from astrocytes. To directly validate this, we examined the expression of one of the shared DEPs, the RBP FUS that is involved in RNA transport and local translation^71,72^. Immunohistochemical analysis revealed a trend toward decreased FUS expession in astrocytes, but not in the CA1 pyramidal cell layer in APP/PS1 mice. This reduction was significantly restored in APP/PS1x*Gfap^-/-^Vim^-/-^* animals (Fig. 5J, Fig. S9A). We similarly validated a different RBP, hnRNPL, which showed a comparable astrocyte-specific reduction in APP/PS1 mice and was rescued upon GFAP/Vim deletion (Fig. 5K). Similar to FUS, hnRNPL expression was unchaged in the CA1 pyramidal layer (Fig. 5K, Fig. S9B). Together, these results suggest that in early AD there is a selective downregulation of RBPs in astrocytes, which can be restored by preventing astrogliosis through GFAP/Vimentin deletion.

### 8. Astrocyte-specific GFAP/Vim deletion restores astrocytic, but not neuronal, protein translation

We next validated whether GFAP/Vim deletion rescues protein synthesis in APP/PS1 mice, and identified whether its origin is astrocytic and/or neuronal. For this purpose, we specifically deleted GFAP and Vimentin in hippocampal astrocytes through a CRISPR/Cas9 approach by expressing the HA-tagged saCas9 under the control of the astrocyte-specific GFAP promoter together with guide RNAs (gRNAs) complementary to GFAP or Vimentin (Fig. 6A, B). Both vectors were injected in the dorsal hippocampus and resulted in high transduction efficiency for CA1 astrocytes (74.9 ± 3.2%) with very high specificity after 6-7 weeks (Fig. 6C-E). This approach reduced GFAP and Vimentin expression to 38.2 ± 3.2% and 70.2 ± 4.1% of WT levels, respectively (Fig. 6F-H). Next, we used this CRISPR strategy for knockout (KO) of GFAP and Vimentin in APP/PS1, and prolonged the expression to 8-9 weeks (Fig. 6B), to account for the long half-life of these proteins^73^. We found that the expression of GFAP in transduced, HA+ hippocampal astrocytes, was decreased to 30.8 ± 5.0% and 24.2 ± 8.0% in 4-month WT and APP/PS1 mice, respectively (Fig. S11A-C). Similarly, Vimentin expression was reduced to 21.5 ± 5.6% in WT mice and 39.7 ± 13.2% in APP/PS1 mice (Fig. S11A-C). No differences were observed in KO efficiency between male and female mice (Fig. S11B).

**Figure 6.**
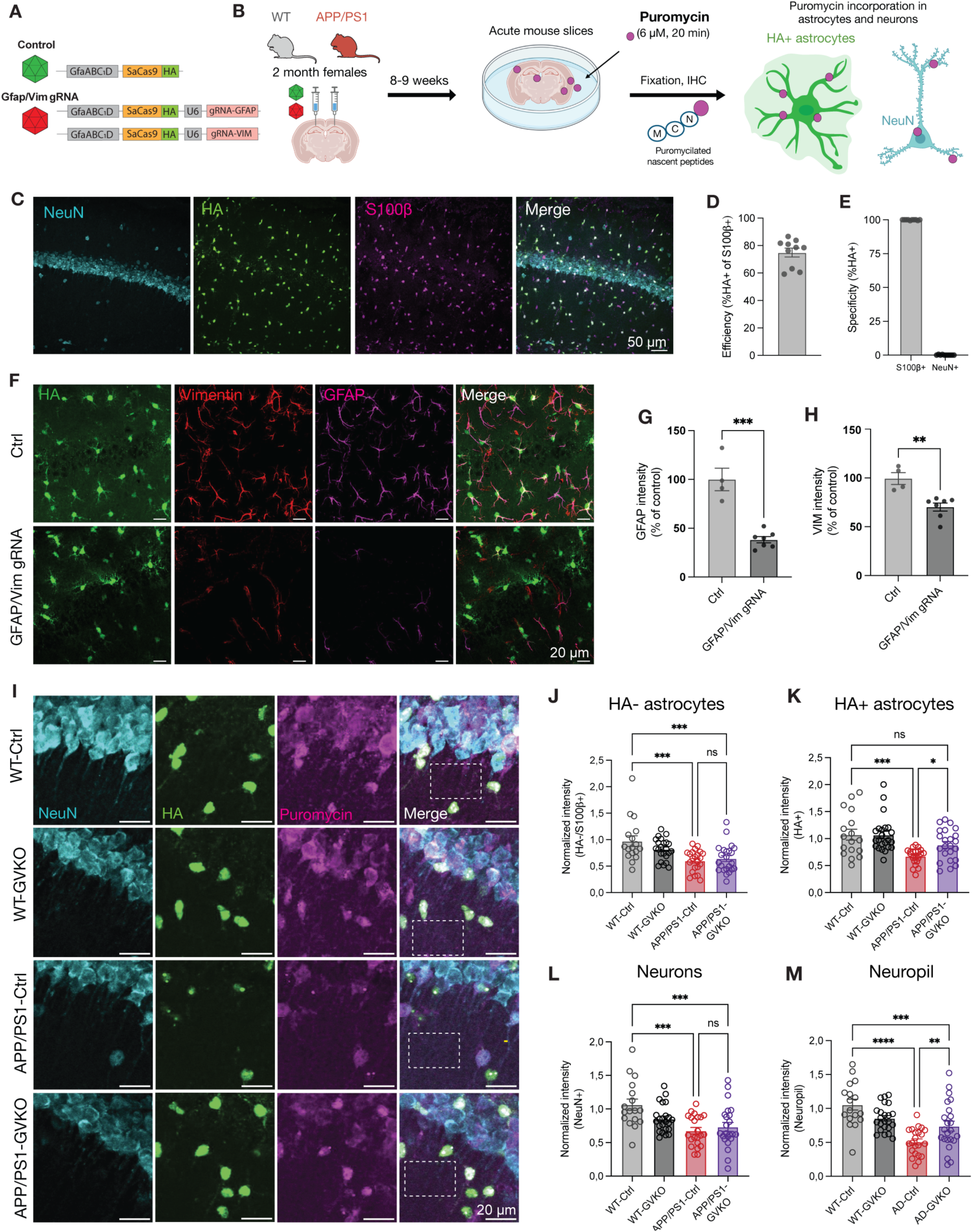
Astrocyte-specific deletion of GFAP and Vimentin in APP/PS1 female mice restores decreased protein synthesis in astrocytes, but not in neurons. **A)** AAVs used to specifically-delete GFAP and Vimentin in astrocytes through CRISPR-Cas9. **B)** Schematic of SUnSET assay for *in vivo* puromycin labelling of newly synthetized peptides in hippocampal neurons and astrocytes using astrocyte-specific CRISPR-Cas9 in 4-month female WT and APP/PS1 mice. **C)** Representative images of the transduction efficiency and specificity. Scale bar, 50 μm. **D)** Quantification of transduction efficiency **E)** and specificity (2472 cells from 10 mice). **F)** Representative images for Ctrl and GFAP/Vim gRNA mice stained for HA, GFAP and Vimentin. The dashed box indicates an example of an ROI selected in the *stratum radiatum* to measure puromycin intensity in the neuropil. Scale bar, 20 μm. **G, H)** Quantification of GFAP (F) or Vimentin (G) protein levels per astrocyte (HA+S100β+). Ctrl: 1256 astrocytes from 4 mice. GFAP/Vim gRNA: 915 astrocytes from 7 mice. **I)** Representative images showing puromycin incorporation (magenta) in neurons (NeuN, cyan) and transduced astrocytes (HA, green). Scale bar, 20 μm. **J-M)** Normalized puromycin incorporation in non-transduced (HA-) astrocytes (I), transduced, (HA+) astrocytes (J), neurons (K) and neuropil (L). n = 17-23 slices from 5 mice/group. Statistical significance calculated by unpaired t-test (G, H) or one-way ANOVA (J-M). See table S3 for full statistical details. *p<0.05, **p<0.01, ***p<0.001, ****p<0.0001, ns: not significant. Data are presented as mean ± SEM.

We then measured astrocyte and neuronal protein translation in acute hippocampal slices from young APP/PS1 mice using surface sensing of translation (SUnSET^74^). In short, labeling with puromycin, an aminoacyl-tRNA analog that incorporates into nascent polypeptide chains was used to tag newly-synthetized peptides, reflecting the global protein synthesis rate. Puromycin incorporation was subsequently detected by immunofluorescence to quantify global cell type-specific protein synthesis rates (Fig. 6B). Pre-treatment with the translational inhibitor anisomycin (200 μM for 10 min) prior to puromycin treatment strongly decreased the puromycin signal in all cell types assessed (Fig. S12A-D), confirming that puromycin labels newly-synthetized peptides.

Next, we investigated protein translation in hippocampal neurons (NeuN+) and astrocytes (S100β+) from 4-month-old APP/PS1 female mice, and found a significant decrease in puromycin incorporation for both cell types compared to WT controls (Fig. 6I-L). Next, we examined the impact of a double KO of GFAP and Vimentin (GVKO) on protein translation by measuring puromycin intensity in transduced (HA+) astrocytes. This revealed that GVKO causes a significant restoration of astrocytic protein synthesis in APP/PS1 mice (Fig. 6I, K). Importantly, analysis of the untransduced (HA-) astrocytes, which still expressed GFAP and Vimentin, did not reveal any differences in puromycin incorporation (Fig. 6J, Fig. 7E), indicating that GFAP and Vimentin upregulation in reactive astrocytes in APP/PS1 mice suppressed protein synthesis. Finally, we examined whether GFAP/Vim deletion in astrocytes affected translation in nearby CA1 neurons. In contrast to astrocytic translation, the reduction in somatic neuronal translation in APP/PS1 mice was not rescued by astrocytic GFAP/Vim deletion (Fig. 6L). As a proxy for synaptic and PAP translation, we measured puromycin intensity in the neuropil by selecting regions of interest (ROIs) in the *stratum radiatum* (Fig. 6I, M). Translation in the neuropil was strongly reduced in APP/PS1 mice, and partially restored in APP/PS1-GVKO mice (Fig. 6M), consistent with a rescue of astrocytic local translation in the PAPs in line with the proteomics data (Fig. S8). In contrast to females, 4-month male APP/PS1 mice did not exhibit significant reductions in astrocytic or neuronal somatic translation (Fig. S13A-E), whereas protein translation in the neuropil was found to be decreased (Fig. S13F). Similarly to females, GFAP/Vim deletion restored this reduction (Fig. S13F). These findings are in line with recent observations of more pronounced astrogliosis in women that may underlie the higher vulnerability of females to AD^75^. Together, these results indicate that astrocytic protein synthesis in APP/PS1 mice shows sex-specific alterations, but consistently depend on astrocytic GFAP and Vimentin in both sexes.

**Figure 7.**
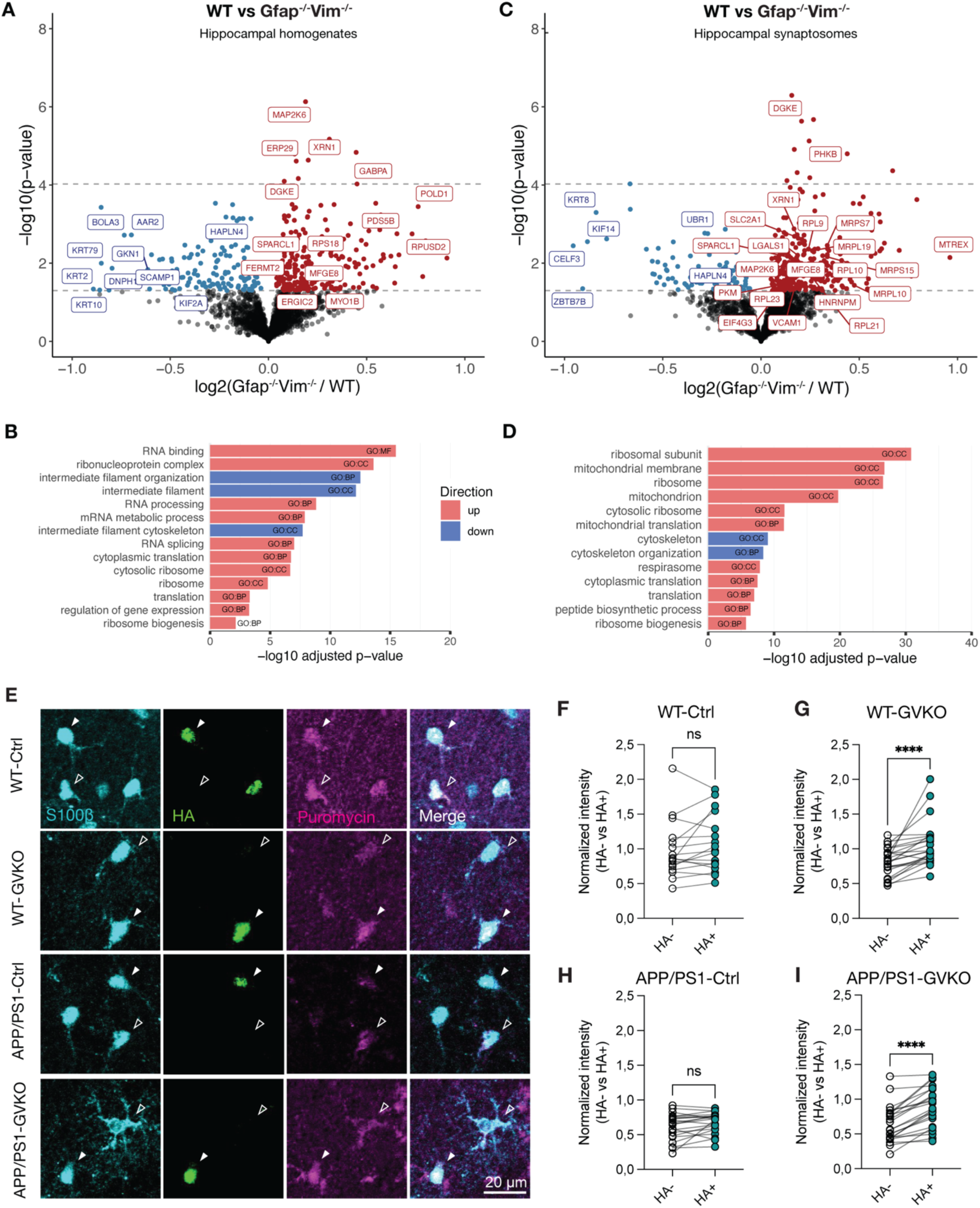
GFAP and Vimentin modulate astrocytic protein synthesis in both WT and APP/PS1 mice. **A)** Volcano plot showing differential expression of proteins in WT vs Gfap^-/-^Vim^-/-^ (WT: n = 6 mice; Gfap^-/-^Vim^-/-^. n = 7 mice) for hippocampal homogenates. Lower dotted line at p<0.05; Upper dotted line at FDR<0.05. Complete DEP list in Table S1. DEPs upregulated in APP/PS1 (red) and downregulated DEPs (blue). **B)** Overview of enriched pathways in downregulated (blue) or upregulated (red) DEPs in WT vs Gfap^-/-^Vim^-/-^ hippocampal homogenates based on GSEA for GO terms (CC, cellular component; BP, biological process; MF, molecular function). Adjusted p-values are FDR-corrected. **C)** Same as in (A), but for hippocampal synaptosomes. **D)** Same as in (B), but for synaptosomes. **E)** Representative images showing puromycin incorporation (magenta) in HA- or HA+ (green) astrocytes (S100β, cyan). Scale bar, 20 μm. **F-I)** Normalized puromycin incorporation in WT-Ctrl (F), WT-GVKO (G), APP/PS1-Ctrl (H) and APP/PS1-GVKO (I). n = 17-23 slices from 5 mice/group. Statistical significance calculated by paired t-test or Wilcoxon matched-pairs test (F-I). See table S3 for full statistical details. ****p<0.0001, ns: not significant. Data are presented as individual values.

### 9. GFAP and Vimentin modulate astrocytic protein synthesis in homeostatic conditions and in Alzheimer’s disease

Lastly, we examined how GFAP/Vim deletion affects astrocytes under homeostatic conditions. For this, we compared the proteomes of WT and *Gfap^-/-^Vim^-/-^*mice in hippocampal homogenates and synaptosomal fractions (Fig. 7A-D). As expected, we observed a downregulation of terms related to IF cytoskeleton and cytoskeletal organization in *Gfap^-/-^Vim^-/-^* mice (Fig. 7B, D). Importantly, the observed upregulation of processes related to protein translation, such as “cytoplasmic translation”, and ribosomal proteins observed upon GFAP and Vimentin deletion in APP/PS1 mice (Fig 4-6) were also found in homogenates and synaptosomes of *Gfap^-/-^Vim^-/-^*mice (Fig. 7B, D). This includes several ribosomal proteins, some of which are previously shown to be enriched in the local PAP translatome^69,70,76^ (Fig. 7A, C). These findings indicate that deletion of GFAP and Vimentin enhances astrocyte-associated translational machinery independently of AD pathology. To validate a cell-autonomous effect of GFAP/Vim deletion on astrocytic protein translation under homeostatic conditions in WT mice, we compared puromycin incorporation between astrocytes transduced with AAVs expressing gRNAs for GFAP and Vimentin (HA+, GVKO astrocytes) and untransduced astrocytes (HA-), within the same hippocampal slice (Fig. 7E-I, Fig. S11B). HA+ GVKO astrocytes of WT mice showed a significantly increased puromycin incorporation, reflecting an increased protein synthesis caused by GFAP/Vim deletion (Fig. 7E-G). A similar increase in protein synthesis was found in GVKO astrocytes of APP/PS1 mice (Fig 7E, H, I). Importantly, in WT and APP/PS1 control slices (Wt-Ctrl, APP/PS1-Ctrl), there was no difference in puromycin intensity between HA- and HA+ cells (Fig. 7E, F, H). Together, these data show that astrocyte IF proteins GFAP and Vimentin play a role in modulating protein synthesis under both physiological conditions and in an AD mouse model.

## Discussion

Reactive astrocytes are increasingly recognized as key players in Alzheimer’s disease pathophysiology, yet the molecular mechanisms governing astrocyte reactivity and its functional consequences remain incompletely understood. Here, we show that intermediate filament (IF) proteins GFAP and Vimentin are required for the widespread hypertrophic response of reactive astrocytes in early AD and play a key role in homeostasis of astrocytic protein synthesis. Genetic ablation of these cytoskeletal proteins in astrocytes of AD mice inhibited astrocyte reactivity while restoring astrocytic protein synthesis and synaptic proteome profiles, and resulted in a robust rescue of cognitive impairment in early AD in both male and female mice.

### Attenuation of astrogliosis prevents early cognitive decline in AD

The IF cytoskeleton, unlike actin and microtubule cytoskeletons, consists of a broad family of structurally related proteins with cell type, activation state and tissue-specific expression patterns^77^. GFAP and Vimentin form the core astrocytic IF network, and their upregulation is a principal hallmark of reactive astrogliosis in neurodegeneration^78^. Because GFAP and Vimentin can partially compensate for the absence of the other, a deletion of both is required to eliminate the IF network and study its function^25,27,79,80^. In line with previous reports, we found that *Gfap*^-/-^*Vim*^-/-^ astrocytes develop normally and are present in typical densities^22,46^, but fail to become hypertrophic under pathological conditions that normally induce astrogliosis^22,23,27^. This supports a critical role for these proteins in the reactive astrogliosis response to injury, while their role for homeostatic astrocyte function is still unclear, especially given the lower GFAP expression of cortical astrocytes compared to hippocampal astrocytes^3^. While the role of GFAP and Vimentin has been examined in late-stage neurodegeneration and in an infantile form of neurodegeneration^81^, their functional contribution to AD progression and associated cognitive impairment had, to our knowledge, not been assessed yet. Therefore, to causally examine the astrocyte-based mechanisms that drive the disease, we focussed on the early stages of AD. Here we report, for the first time, that deletion of GFAP and Vimentin prevents cognitive impairment in early AD, with APP/PS1x*Gfap*^-/-^*Vim*^-/-^ mice showing improved contextual fear memory, spatial memory and nest building behavior compared to APP/PS1 mice. These behavioral findings align with our previous work showing that pre-plaque stage 4-month-old APP/PS1 mice display hippocampal synapse loss and synaptic plasticity deficits, correlating with their behavioral deficits^33^. No differences were seen between WT and *Gfap*^-/-^*Vim*^-/-^ mice, consistent with previous research that reported normal cognition in *Gfap*^-/-^*Vim*^-/-^mice^26^. Furthermore, we found that while GFAP/Vim deletion prevented cognitive decline, plaque load was unaffected in 4-6 months APP/PS1x*Gfap*^-/-^*Vim*^-/-^ mice, although effects at later stages were not examined, so they cannot be excluded. Studies examining more advanced AD stages in these mice found opposite effects on Aβ pathology^23,46^, possibly driven by disease stage and the different time points examined. Also, whereas astrogliosis was prevented in APP/PS1x*Gfap^-/-^Vim^-/-^*mice, microgliosis was not (Fig. 3), consistent with previous studies that show that the modulation of astrocyte reactivity does not affect the number of plaque-contacting microglia or their transcriptional profiles^23,48^. This supports the notion that microglial reactivity, like in other pathological conditions^82–85^, occurs upstream of astrocyte reactivity in AD. Importantly, this selective suppression of astrocyte reactivity, without affecting microglia and Aβ pathology, allowed us to isolate astrocyte-specific contributions to early AD pathogenesis. Our results show that attenuating astrogliosis is sufficient to protect against early cognitive decline and to preserve hippocampal function, even in the presence of microgliosis and Aβ pathology, thereby identifying reactive astrocytes as key drivers of early functional deficits in AD.

### Astrocytic protein synthesis: mechanisms and implications to AD

Our proteomic analyses identified dysregulation of astrocytic RNA metabolism and protein synthesis as an early event in AD. These findings align with transcriptomics studies reporting early astrocytic changes in AD mouse models and patients^7,86,87^, which included the response to endoplasmic reticulum (ER) stress, impaired protein homeostasis, and altered expression of ribosomal proteins or RBPs^51,88–92^. Our data extend these reported findings by showing that proteostasis defects in AD astrocytes are not limited to regulation at the transcript level, but are also reflected at the protein level, and that these alterations are already present at the pre-plaque stage of AD. Moreover, we found that the alterations in protein synthesis were more pronounced in the neuropil than in the astrocytic somas and main processes, indicating that the reduced astrocytic protein translation in early AD is most prominent in PAPs and fine processes. In support of this idea, we show that the level of previously published locally translated PAP transcripts and PAP proteins was altered in APP/PS1 and restored by GFAP/Vim deletion, as evidenced by the significant inverse correlation (Fig. S9). This suggests that alterations in local protein synthesis of astrocytic PAPs are implicated in early AD pathology.

The restoration of astrocytic translation and the associated rescue of synaptic proteins that we observed in APP/PS1x*Gfap^-/-^Vim^-/-^* mice, provides a plausible mechanistic explanation on how prevention of reactive astrogliosis rescued learning and memory in APP/PS1 mice. Previously, we found that astrocytes undergo activity-dependent changes in their translatome, including PAP proteins involved in modulation of synaptic activity^76^. Indeed, several astrocytic mRNAs are locally translated within PAPs, including mRNAs encoding AD-related proteins APOE and CLU, as well as proteins essential for sustainining synaptic function^69,93–95^. Defects in neuronal proteostasis and translation have long been recognized in AD^96–98^, and have been linked to impairments in neuronal function, synaptic plasticity and memory^67,99–104^. Interestingly, the memory deficit in early AD parallels the retrieval deficit caused by inhibition of protein synthesis^105^. Astrocytes, however, are only recently recognized as active regulators of these processes^106–108^, and modulate them in part through local translation in their PAPs^70,76,109^. Accordingly, increasing protein translation specifically in hippocampal astrocytes, by downregulation of p-eIF2α, was shown to enhance long-term potentiation, excitatory synaptic transmission and memory formation^109^, which directly links astrocytic protein translation to cognitive function. In addition, impaired local translation in astrocytes has also been implicated in the pathogenesis of amyotrophic lateral sclerosis (ALS) and frontotemporal dementia (FTD)^94^, suggesting that astrocyte-specific translational dysregulation may represent a shared mechanism across neurodegenerative diseases. Together with longstanding evidence for translational perturbations in AD^67,98,99,102,104^, our findings support the emerging view that reduced protein synthesis in AD is not only restricted to neurons but also occurs in astrocytes. This astrocytic translational impairment has functionally relevant consequences for synaptic integrity and memory, positioning astrocytic protein synthesis as a critical and previously underappreciated component of early AD pathogenesis.

Our observations support a role for IF cytoskeleton proteins GFAP and Vimentin in regulating astrocytic protein translation, in both WT and APP/PS1 mice. In addition, our proteomic analyses provided insight into the underlying mechanisms. Several of the RBPs that were upregulated in *Gfap^-/-^Vim^-/-^* mice, including FUS and hnRNPs^110–112^, have previously been associated to AD pathology. In neurons, FUS has a role in local translation at axons and synapses^71,72^, and may serve a similar function in local translation in astrocytes. Consistent with this notion, the IF cytoskeleton is known to interact with translation factors, RBPs and ribosomal proteins^113–115^, indicating a direct role for IF proteins in local translation. Specifically, vimentin has a non-canonical function as an RBP, interacting with other RBPs to stabilize mRNAs, as was shown for collagen mRNA^116^. Moreover, IFs are critical for sensing and responding to cellular stress, in particular oxidative stress, a process characteristic of AD and influenced by astrocytes^101,117–119^. Notably, keratins, the main IF proteins in epithelial cells, were shown to modulate protein synthesis through mTOR signaling^120^. Similarly, vimentin has been reported to regulate mTOR signaling in fibroblasts^121^. This is likely conserved in astrocytes, since reactive astrocytes in the hippocampus of APP/PS1 mice present reduced mTOR/pS6 signalling, and reactivation of mTOR signalling recovers their normal morphology^88^. Alternatively, vimentin functions as a signalling scaffold^39,62,63,80^, and modulates proteostasis^118,122–124^ and the integrated stress response (ISR) pathway^125^. In neurodegeneration, the ISR is active in reactive astrocytes^126–128^. Recently, Chen et al., reported that astrocytic translation is decreased in AD models, due to chronic activation of astrocytic PERK and downstream eIF2α phosphorylation^126^. Similarly, Tapella et al. showed that hippocampal AD mouse astrocytes exhibit reduced protein synthesis driven by eIF2α phosphorylation^129^. In line with the described role of IF proteins in translational control in other cell types, our data identify the IF proteins GFAP and Vimentin as key modulators of protein translation in astrocytes, with direct relevance for astrogliosis and AD pathogenesis.

### Limitations of the study

Germline deletion of GFAP and Vimentin in *Gfap*^-/-^*Vim*^-/-^ mice caused male-specific renal dysfunction, consistent with previous observations linking vimentin loss to shorter life expectancy^130^ and renal failure, possibly through the impaired modulation of vascular tone^40,131^. Why GFAP/Vim-associated mortality is sex-specific remains unclear. To minimize animal suffering and avoid confounding systemic effects, we focused on female mice, which prevented in-depth examination of the effect of GFAP/Vim deletion in male mice. While data from a small number of *Gfap*^-/-^*Vim*^-/-^male mice confirmed that GFAP/Vim deletion prevents cognitive decline also in male mice, this data should be interpreted with caution because these mice could be experiencing discomfort. In addition, *Gfap*^-/-^*Vim*^-/-^ mice lack these proteins in all cell types, including endothelial cells that normally express Vimentin^132^ and radial glia that express GFAP^133^, and therefore the proteomic regulation observed cannot solely be attributed to astrocytes. Despite these limitations, CRISPR/Cas9-mediated astrocytic GFAP/Vim deletion resulted, like for females, in a restored protein translation in the neuropil for male mice, extending the relevance of our observations to both sexes.

### Outlook

Recent studies in human cohorts of AD and mild cognitive impairment have demonstrated that astrogliosis emerges early, during the asymptomatic stages of AD, and that this influences downstream neurodegenerative biomarkers^10,134^. CSF and plasma concentrations of GFAP are elevated in preclinical AD, and reliably predict future disease progression in patients^10,14,15,135^, underscording the translational relevance of targeting astrocytic responses in human AD. Although vimentin loss in the periphery is associated with renal dysfunction, selective astrocytic manipulation of these proteins or pathways involved in protein synthesis may hold therapeutic potential. Specifically, astrocytic IFs represent a promising therapeutic target, since we showed that interfering with them prevents astrogliosis and cognitive decline in AD mice, while *Gfap^-/-^Vim^-/-^*display normal behavior. Further, other studies showed that GFAP/Vim deletion improved regeneration after injury^24,26,28^. Moreover, vimentin has been proposed as a potential target for cancer treatment^136^, and several vimentin targeting compounds are currently in clinical trials for differenty cancer types^136–139^. Whether these compounds hold potential for the treatment of brain disorders, like AD and other neurodegenerative disorders, needs to be explored. In this study, we focused on the hippocampus, since it represents one of the earliest affected regions in AD and is crucial for learning and memory^140,141^. However, given the recently described astrocyte heterogeneity amongst brain regions^142^, it remains to be determined whether GFAP/Vim similarly modulate protein synthesis elsewhere in the brain. Interestingly, a constitutive reduction in GFAP expression during brain evolution, such as that observed for cortical astrocytes, has been proposed to confer functional advantages such as reduced unnecessary protein synthesis and increased astrocyte adaptability^143^, in accordance with our observations of increased translation after GFAP/Vim deletion. Finally, part of the RBP alterations that we find are also present in FTD, where proteomic analyses on human FTD brain tissue identified a dysregulation of RNA processing in astrocytes^144,145^, suggesting common mechanisms of astrocytes across neurodegenerative diseases. Taken together, our findings highlight the need for more research into the molecular mechanisms of astrocyte reactivity, including protein synthesis, as a critical component of AD pathogenesis and as a promising source of novel cell type specific therapeutic targets.

## Conclusion

In summary, we find that interfering with GFAP and Vimentin-dependent astrocyte reactivity prevents early cognitive impairments in the APP/PS1 mouse model, independently of microgliosis or Aβ accumulation. Through integrated proteomic and protein synthesis analyses we reveal IFs as key regulators of astrocytic translation, with downstream consequences for modulating synaptic proteostasis. Our findings support a causal model for early AD pathogenesis, in which reactive astrocytes upregulate GFAP and Vimentin, leading to suppression of protein synthesis in astrocytes, disruption of synaptic proteostasis, and memory impairment. Targeting astrocytic translational control therefore emerges as a promising strategy to counteract early cognitive dysfunction in Alzheimer’s disease.

## Materials and methods

### Animals

Generation of APP/PS1x*Gfap^-/-^Vim^-/-^*mice on a C57Bl/6J background: Mice heterozygous for null mutations in the *Gfap* and *Vim* genes, on a mixed background of C57Bl6/129Sv/129Ola^27^, were obtained from Prof. Milos Pekny via Janvier Labs and crossed to generate *Gfap^-/-^Vim^-/-^* mice (Fig. S1). Next, mice were backcrossed to C57Bl/6J mice, and MAX-BAX® speed congenics (Charles River Laboratories) was used to determine the coverage of background genes and to select the most suitable breeding pairs. After five generations, *Gfap^-/-^Vim^-/-^* mice were crossed with transgenic APPswe/PSEN1dE9 (APP/PS1) mice, a commonly used AD mouse model for increased amyloidosis, which co-expresses the Swedish double mutation K595N/M596L (APPswe) and the human PSEN1 gene harbouring an exon 9 deletion (PSEN1dE9), both under the control of the mouse prion protein promotor (MoPrP.Xho)^20^. The two knockout genes (Gfap and Vim) and transgenes (APPswe and PSEN1dE9) segregated independently and therefore multiple generations were required to produce the four genotypes that we aimed to study: WTx*Gfap^+/+^Vim^+/+^* (WT), APP/PS1x*Gfap^+/+^Vim^+/+^* (APP/PS1), WTx*Gfap^-/-^Vim^-/-^* (*Gfap^-/-^Vim^-/-^)* and APP/PS1x*Gfap^-/-^Vim^-^*

*^/-^* mice (Fig. S1A). For the temporal analysis of GFAP we used conventional male WT and APP/PS1 mice of either 2, 3, 4 or 6 months of age. For experiments including *Gfap^-/-^Vim^-/-^*and APP/PS1x*Gfap^-/-^Vim^-/-^* models, only female mice were included, unless specified otherwise. Females were used because we encountered an unexpected health issue affecting only male mice, which resulted in unforeseen mortality (Fig. S1C). From 2 months of age onwards, we found that male APP/PS1x*Gfap^-/-^Vim^-/-^* had a lower body weight compared to conventional APP/PS1 mice (Fig. S1B). No differences in body weight were observed between genotypes for female mice. Male *Gfap^-/-^Vim^-/-^*and APP/PS1x*Gfap^-/-^Vim^-/-^*experienced sudden death starting at the age of 2 months (Fig. S1C), potentially due to renal failure (Fig. S1D). This was not observed for female *Gfap^-/-^Vim^-/-^*and APP/PS1x*Gfap^-/-^Vim^-/-^.* These mice remained healthy until at least the observed age of 4 months. Whether health issues occur in female mice older than 4 months was not determined. Due to the inability to predict which mice were susceptible to this condition, we made the ethical decision to exclude male mice from future experiments to prevent unnecessary suffering. In addition, both males and females experienced AD-related mortality due to epileptic seizures, starting at the age of 7 weeks, as previously reported^146^. This was observed for both conventional APP/PS1 mice and APP/PS1x*Gfap^-/-^Vim^-/-^.* Mice were group housed (2 to 4 mice), unless specified otherwise, and the cages were enriched with sawdust bedding, nesting material, gnawing stick and a tube. Housing was controlled for temperature (∼21 °C), humidity (∼50-55%) and 12 h light-dark cycle. We minimized the number of required animals by performing post-mortem immunoblotting and immunohistochemical experiments on the tissue of mice that were used in behavioral analyses. When this was the case, animals were sacrificed one week after the last behavioral experiment. All experimental procedures were approved by the local ethical committees and comply with the European Council Directive 2010/63/EU.

### Immunohistochemistry

Mice were sedated with 250 mg/kg Avertin and transcardially perfused with ice-cold PBS (pH 7.4) followed by ice-cold 4% paraformaldehyde (PFA) in PBS (pH 7.4). Whole brains were extracted, post-fixed in 4% PFA for 24h and transferred to 30% sucrose solution for cryoprotection, and finally stored at -80 °C until further processing. Coronal cryosections of 15 - 35 μm were taken at the level of the hippocampus and kept in PBS with 0.02% NaN_3_. Antigen retrieval was performed prior to staining with antibodies against VIM and consisted of slice incubation with 0.05% Tween-20 (pH 6.0) for 20 min at 95 °C. To prevent non-specific antigen binding, the sections were incubated in blocking solution (0.2% Triton-X (Sigma Aldrich), 2.5% bovine serum albumin (Sigma Aldrich) and 5% normal goat serum (Sigma Aldrich) in PBS), for 1 hour at room temperature (RT). Antibodies were used against GFAP (1:1000, Dako, #Z0334 or 1:2000, SynapticSystems, #173004), VIM (1:4000, Sigma-Aldrich, #5733 or 1:1000, Abcam, #ab92547), S100β (1:250, Sigma-Aldrich, #S2532 or 1:600, Dako, #Z0311), IBA1 (1:500, SySy, #234308), CD68 (1:100, Novus Biologicals, #NB100-683), Aβ-6E10 (1:1000, ITK Diagnostic, #SIG-39320) FUS (1:1000, Life Technologies, #A300-293A), hnRNPL (1:1000, Santa Cruz, #sc-32317) and HA (1:750, Roche, #11867423001). After washing, slices were incubated for 2 h at RT with corresponding Alexa-conjugated secondary antibodies (Invitrogen). Slices were incubated with 4’,6-diamidino-2-phenylindole (DAPI, Thermo Fisher, #D3571). Finally, slices were mounted on glass slides. Four-Diazobicyclo-[2,2,2]-octane (DABCO™, Sigma Aldrich, #10981) was added as antifading agent before slices were cover-slipped.

### Confocal microscopy and image analysis

The experimenters were blinded for the experimental groups while imaging and during analysis. Imaging for GFAP, VIM, S100β and GS quantification was performed using a Zeiss confocal microscope (LSM510). For quantitative analysis, a total of 12 z-stacks (step size of 1 μm, 512 x 512 pixels) of the hippocampal CA1 region were acquired per animal at 400x magnification (∼0.05 mm^2^). For the analysis of Aβ, images were acquired with a Nikon A1 confocal microscope with NIS-Elements acquisition software. Here, one z-stack including the cortex and hippocampus were taken at 50x magnification (∼10 mm^2^). For quantification of FUS, hnRNPL and the CRISPR/Cas9-KO efficiency, the same Nikon A1 confocal microscope was used, and z-stacks (step size of 2 μm, range 10-30 μm, 1024 x 1024 pixels) were acquired from the hippocampal CA1 with a 40x objective (NA 1.35) with the experimenter blinded to the condition. Image rendering and analysis were performed in ImageJ (FIJI, 2.1.0., National Institutes of Health, USA). The levels of GFAP, VIM, S100β, GS, FUS and hnRNPL were determined by quantification of the levels per individual astrocyte, measured over all astrocytes in a single image resulting from the maximum-intensity projection of the acquired z-stack. Weused ImageJ to automatically outline intact complete cells that are not located at the border of the image. Each outlined cell was detected as a single object in which staining intensity was measured. To measure FUS and hnRNPL intensity in the PCL, the PCL was manually outlined in the DAPI channel, and subsequently fluorescence intensity was measured in the PCL-ROI. IBA+ microglia were manually counted using the “cell counter” plugin in ImageJ. For Aβ plaques analysis, the surface area of the full hippocampus and cortex were determined per individual image. Plaque density was subsequently determined by a brightness-dependent threshold analysis of the Aβ+ signal per brain region. To quantify GFAP and Vimentin for CRISPR/Cas9-KO efficiency, ROIs were manually selected around HA+ astrocytes, excluding those overlapping with artifacts or blood vessels. Background intensity was substracted and GFAP and Vimentin intensity were calculated relative to control.

### Electron microscopy

Naïve 4-month female mice were transcardially perfused with 4% PFA) in 0.1M PBS (pH 7.4). Brains were dissected and post-fixated in 4% PFA for 24 h followed by cryopreservation in 30% sucrose and stored at -80 °C until further processing. 50 μm thin coronal slices of the hippocampus were made on a cryostat and kept in PBS for a maximum of 24 h. Sample processing was performed as previously reported^17^. In short, free-floating sections were contrasted with 1% osmium tetroxide and 1% ruthenium. Consequently, the sections were dehydrated by increasing ethanol concentrations (30, 50, 70, 90, 96, 100%) and propylene oxide followed by embedding in epoxy resin and incubation for 72 h at 65 °C. An ultra-microtome (Reichert-Jung, Ultracut E) was used to generate 90 nm thin slices of the hippocampus CA1 region. Finally, the slices were post-contrasted with uranyl acetate and lead citrate in an ultra stainer (LEICA EM AC20). A Tecnai transmission electron microscope was used to acquire digital images of the hippocampus CA1 stratum radiatum at 60.000 x magnification. A total of 40-70 randomly selected images were captured from 2 WT and 3 Gfap^/-/^Vim^-/-^ and qualitatively explored to identify astrocyte contact with synapses.

### Immunoblotting

Mice were sedated with 250 mg/kg Avertin and subsequently transcardially perfused with ice cold 0.1M phosphate buffered saline (PBS; pH 7.4). The brains were removed, and bilateral hippocampi dissected and immediately frozen in liquid nitrogen followed and stored at -80 °C until used. The hippocampi were homogenized in ice cold homogenization buffer (0.32 M sucrose, 5 mM HEPES (pH 7.4) and protease inhibitor cocktail (Roche)) using a Teflon pestle, rotating at 900 rpm, attached to a dounce homogenizer. Supernatant was collected after centrifugation at 1000 g for 10 min at 4 °C. A Bradford assay (Bio-Rad) was performed to determine the protein concentration of each sample. Next, 2.5 μg (for gliosis markers on whole hippocampus samples) or 8 μg (for temporal analysis samples) of protein was incubated with Laemmli SDS sample buffer. Samples were denatured at 95 °C for 5 min and subsequently loaded and separated on a 4-15% Mini-PROTEAN® TGX™ Precast Protein Gel (Biorad), and run at 75V until samples reached the stacking gel followed by protein separation at 150V. Next, the samples were electroblotted onto a methanol-activated polyvinylidene difluoride membrane (PVDF, Bio- Rad) at 40V over-night (o/n). After blocking the membranes with 5% non-fat dry milk and 0.5% Tween-20 in Tris Buffered saline (TBS) for 1 h at RT, the membranes were incubated o/n at 4 °C with antibodies against GFAP (host rabbit, 1:1000, Dako, Z0334), GS (host rabbit, 1:1000, GenScript, A01305) and VIM (host mouse, 1:100, Santa Cruz, sc-373717), dissolved in 3% non-fat dry milk and 0.5% Tween-20 in TBS. This was followed by 1h RT incubation with HRP conjugated secondary antibodies matching the host of the primary antibody. Immunodetection was performed using chemiluminescence Femto detection (Super Signal West Femto Maximum Sensitivity Substrate, Thermo Scientific) and an Odyssey FC imaging system (LI-COR Biosciences). The protein band immunoreactivity was determined using Image Studio Lite (5.2, LI-COR Biosciences) and normalized to the total protein intensity detected by scanning the 2,2,2-trichloroethanol (TCE) staining of the gel using Image Lab (5.2.1, LI-COR Biosciences).

### ELISA

A human amyloid-beta 42 ELISA kit (Invitrogen, CA, USA) was used to quantify the levels of soluble human Aβ_1-42_ (Aβ_42_) in mouse hippocampus homogenate. Samples were prepared according to the manufacturer’s protocol. In brief, diluted hippocampus homogenate samples, standard curve samples and controls were loaded as duplicates onto a 94-wells plate and mixed with Aβ_42_ detection antibody solution taken from the ELISA kit and incubated for 3 hours. Next, the samples were exposed to anti-rabbit HRP, followed by stabilized chromogen solution and finally a stop solution, with washing steps with washing solution taken from the ELISA kit in between. Light absorbance was measured for each well at 450 nm using a microplate reader (Thermo Fisher). The values were normalized to total protein input as determined by a Bradford assay (BioRad). Finally, the values were averaged per genotype and mean WT values were subtracted as background to obtain the final human Aβ_42_ peptide levels.

### Behavior

In short, 4-month-old group housed female and 6-month-old individually housed male mice were exposed to a behavioral battery including the nesting test, Morris water maze (MWM), contextual fear conditioning and open field. Mice were handled for three consecutive days prior to the behavioral tests to limit experimenter-induced stress responses during the experiments. All of the behavioral experiments used were described previously in detail in Kater et al., (2023)^33^.

#### Nesting test

In brief, mice were housed over-night in a novel cage solely containing sawdust bedding and a pressed cotton square (5 cm x 5 cm). The morning after, the nest completeness was scored by two independent blinded observers and scored according to a five-point scale as described by Deacon (2006)^147^ Scores were averaged per mouse and subsequently averaged per genotype.

#### MWM

The mice were trained for four consecutive days with four training sessions per day and at day five the probe test was performed. During training, mice had to find the submerged platform within 60 s. The time required to find the platform was measured and averaged over the four trials per day per mouse. During the probe test, the platform was removed, and the time spent in each quadrant of the pool was measured by video tracking software (Viewer, Biobserve, Fort Lee NJ).

#### Contextual fear conditioning

Mice were placed in a chamber with a stainless-steel grid floor (MedAssociates Inc, USA) and allowed to explore the arena for 120 s after which they received three foot-shocks (0.7 mA, 2 s) with a 60 s interval. During the retrieval session, 48 h later, mice returned to the same context. Freezing levels (defined as a lack of movement for a minimum of 1.5 s) were determined by video tracking for four minutes (Video Freeze, MedAssociates, USA).

#### Open field

The mice were placed in an arena in which their behavior was tracked for 10 min (Viewer, Biobserve). Enhanced anxiety-like behavior was determined when mice spent less time in the inner zone. In addition, velocity was measured as parameter of locomotor activity.

Mice were sacrificed a week after the last behavioral test in order to use the brain tissue for immunohistochemistry and western-blotting.

### Tissue collection for MS proteomics and synaptosome isolation

After the behavioral assays, 4-month female mice were sacrificed via cervical dislocation. The bilateral hippocampi were dissected from the brain, frozen on dry ice and then stored at -80°C until use. The dorsal hippocampus was homogenized in homogenisation buffer (0.32M sucrose, 5mM HEPES, in PBS. pH 7.4 with protease inhibitor cocktail (Roche) using a Dounce homogenizer for 12 strokes at 900 rpm. After centrifugation at 1000g for 10 min, the supernatant was collected. Part of the hippocampal homogenates was stored at -80°C until sample preparation of MS. The rest of the homogenates were used to isolate synaptosomes on a discontinuous sucrose gradient as described previously^148^. In brief, the homogenates were centrifuged at 100,000g for 2 h in a 0.85-1.2 M sucrose gradient. Synaptosomes were collected from the interface, diluted, and centrifuged at 18,000g for 30 min to collect a synaptosome rich pellet that was resuspended in 75 μl of 5mM HEPES/NaOH pH = 7.4. All steps were performed on ice or at 4°C settings. Lastly, a Bradford assay was performed to determine the protein concentration.

### MS sample preparation using suspension-trapping (S-TRAP)

MS sample preparation was performed following a DNA micro spin column suspension trapping protocol^149^, with some minor adaptations. For each sample, 10 μg of protein was mixed with6.3% SDS (Sigma Aldrich), 50 mM Tris-HCl (pH 8.0) and 5 mM tris(2-carboxyethyl)phosphine (TCEP) and incubated at 55 °C, 1500 rpm for 15 min. Free sulfhydryl groups were alkylated by incubation with 20 mM methyl methanethiosulfonate (MMTS; 200 mM stock solution) for 15 minutes at RT. Next, samples were acidified by addition of 1.1% phosphoric acid (12% stock solution), mixed with six volumes of binding/washing buffer (90% methanol and 100mM Tris-HCl, pH 8) and loaded onto plasmid DNA micro-spin columns (HiPure from Magen Biotechnology). The protein suspension was retained in the column following centrifugation at 1400x g for 1 minute and the columns were washed four times with binding/washing buffer. Columns were transferred to new collection tubes (Eppendorf), supplemented with 0.4 μg Trypsin/Lys-C (Promega) in 50 mM NH_4_HCO_3_ and incubated overnight at 37°C in a humidified incubator. Tryptic peptides were eluted and pooled by subsequent addition of 50 mM NH_4_HCO_3_, 0.1% formic acid and 0.1% formic acid in acetonitrile. Collected peptides were dried by SpeedVac and stored at -80°C.

### LC-MS analysis

Each sample of tryptic digest was redissolved in 0.1% formic acid and the peptide concentration was determined by tryptophan-fluorescence assay^150^. 75 ng of peptide was loaded onto an Evotip Pure (Evosep). Peptide samples were separated by standardized 30 samples per day method on the Evosep One liquid chromatography system, using a 15 cm × 150 μm reversephase column packed with 1.5 μm C18-beads (EV1137 from Evosep) connected to a 20 μm ID ZDV emitter (Bruker Daltonics). Peptides were electro-sprayed into the timsTOF Pro 2 mass spectrometer (Bruker Daltonics) equipped with CaptiveSpray source and measured with the following settings: Scan range 100-1700 m/z, ion mobility 0.65 to 1.5 Vs/cm^2^, ramp time 100 ms, accumulation time 100 ms, and collision energy decreasing linearly with inverse ion mobility from 59 eV at 1.6 Vs/cm2 to 20 eV at 0.6 Vs/cm^2.^ Operating in dia-PASEF mode, each cycle took 1.38 s and consisted of 1 MS1 full scan and 12 dia-PASEF scans. Each dia-PASEF scan contained two isolation windows, in total covering 300-1200 m/z and ion mobility 0.65 to 1.50 Vs/cm^2^. Dia-PASEF window placement was optimized using the py-diAID tool^151^. Ion mobility was auto calibrated at the start of each sample (calibrant m/z, 1/K0: 622.029, 0.992 Vs/cm^2^; 922.010, 1.199 Vs/cm^2^; 1221.991, 1.393 Vs/cm^2^).

### MS data analysis

DIA-PASEF raw data were processed with DIA-NN 1.8.1^152,153^. An in-silico spectral library was generated from the Uniprot human reference proteome (SwissProt and TrEMBL, canonical and additional isoforms, release 2023-02) using Trypsin/P digestion and at most 1 missed cleavage. Fixed modification was set to beta-methylthiolation(C) and variable modifications were oxidation (M) and N-term M excision (at most 1 per peptide). Peptide length was set to 7-30, precursor charge range was set to 2-4, and precursor m/z range was limited to 280–1220. Both MS1 and MS2 mass accuracy were set to 15 ppm, scan window was fixed at 9, and the precursor False Discovery Rate (FDR) was set to 1%. Heuristic protein inference was disabled, protein identifiers (isoforms) were used for protein inference, double-pass mode and match-between-runs (MBR) were enabled. All other settings were left as default. Data quality control and statistical analysis were performed by using MS-DAP 1.0.6^56^ for downstream analyses of the DIA-NN results. Filtering and normalization was applied to respective samples per statistical contrast. Peptide-level filtering was configured to retain only peptides that were confidently identified in at least 50% of samples per sample group. Peptide abundance values were normalized using the VSN algorithm, followed by protein-level mode-between normalization. No outlier samples were identified in the quality control analyses presented in the MS-DAP report (methods S1), Differential expression analysis was performed by the MSqRob algorithm. Proteins were considered differentially regulated when the FDR-corrected p-value was less than 0.05, unless specified otherwise.

### Pathway and cell type enrichment analysis of proteomics data

#### Gene set analyses

GOAT online (version 1.0.2, http://ftwkoopmans.github.io/goat^154^) was used to perform gene set enrichment analyses using gene sets from the Gene Ontology database (version gene2go_2025-01-01^64^) and SynGO knowledgebase (SynGO v1.2^155^). The input gene list contained 7083 genes and its effectsize-derived gene scores were used to test for enriched gene sets that contained at least 10 and at most 1500 genes (or 50% of the gene list, whichever was smaller) that overlapped with the input gene list. Multiple testing correction was independently applied per gene set "source" (i.e. GO_CC, GO_BP, GO_MF) using the Benjamini-Hochberg procedure (FDR) and subsequently, all p-values were adjusted (again) using Bonferroni adjustment to account for 3 separate tests across "sources". The significance threshold for adjusted p-values was set to 0.05.

#### G:Profiler

Gene ontology (GO) enrichment analysis was performed using the g:Profiler web server (version e109_eg56_p17^156^). The analysis was performed on default settings, with as background input the total of DIA detected proteins for all samples. The P-values were corrected for multiple testing using the Benjamini-Hochberg method. The output of the analysis was a list of functional categories enriched in the set of differentially expressed genes. These categories were divided into biological processes (BP), cellular components (CC) and molecular functions (MF).

#### SynGO

To identify proteins of synaptic origin, we applied a Synaptic gene ontology analysis (SynGO v1.2^155^). Multiple testing correction was false discovery rate-based and we used the total of DIA quantified proteins as background for the analysis.

**Cell type enrichment analysis** was conducted using published MS proteomics data by Sharma et al^68^. Proteins were annotated to a specific cell type (neuron, astrocyte, microglia or oligodendrocyte) wehen the expression was at least 2-fold higher for a specific cell type to the cell type with the second-highest protein level. All significantly-enriched pathways are reported in table S2.

### Constructs and stereotactic in vivo microinjections of AAVs

The pAAV-GfaABC1D::SaCas9-HA-GFAP and pAAV-GfaABC1D::SaCas9-HA-VIM was generated by first replacing the cytomegalovirus promoter from pX601-AAV-CMV::NLS-SaCas9-NLS-3xHA-bGHpA;Bsal-sgRNA (Addgene # 61591)^157^ with the GFAP promoter (GfaABC1D). Next we inserted the designed single guide RNA (sgRNA) to target GFAP (CCTGCCAGGGTGGACTC) or Vimentin (CAGGGCATCGTTGTTCCGGT). Finally, we cloned the modified plasmid to an AAV2/5 vector. The control virus, pX601-AAV-GfaABC1D::SaCas9-HA, was generated following the same procedure, but it lacks the sequence to target GFAP or Vimentin. This resulted in the following AAVs: AAV-GfaABC1D::SaCas9-HA-GFAP (titer (GC/ml): 1.39e^11^, 0.5 μl/hemisphere), AAV-GfaABC1D::SaCas9-HA-VIM (titer (GC/ml): 5.6e^11^, 0.5 μl/hemisphere). AAV-GfaABC1D::SaCas9-HA-Ctrl (titer (GC/ml): 7.6e^9^, 1 μl/hemisphere).

For stereotaxic microinjection surgeries, mice received 0.067 mg/ml of carprofen (Rimadyl cattle, Zoetis B.V., the Netherlands) in the drinking water one day before surgery. On the day of the surgery, mice were anesthetized with 1.5 – 2.5% isoflurane, mounted onto a stereotactic frame and maintained under isoflurane anesthesia during the surgery. Lidocaine (2%, Sigma-Aldrich Chemie N.V, The Netherlands) was topically applied to the skull to provide local analgesia. The AAV vectors were infused bilaterally in the dorsal hippocampus (in mm relative to Bregma: AP: -2, DV: - 1.5, -1.7, -2, ML: ± 1.5). The desired volume was infused using a microinjection glass needle at a rate of 0.15 μl/min, with a total volume of 1 μl/hemisphere and 0.33 μl for each DV spot. AAV-GfaABC1D::SaCas9-HA-GFAP and AAV-GfaABC1D::SaCas9-HA-VIM were mixed in a 1:1 ratio, so a total of 0.5 μl of each virus was injected per hemisphere. Following infusion, the needle was left in place for an additional 5 min to allow virus diffusion, and was then slowly retracted over 2 min. After surgery, mice remained in their home-cage for 7-9 weeks until the start of experiments.

### Tissue collection for protein synthesis measurements

Protein synthesis measurements were performed 7-9 weeks after AAV injection. 4-month old WT and APP/PS1 mice were anaesthetized by avertin injection prior to intracardial perfusion with ice-cold partial sucrose solution containing: 70 mM sucrose, 25 mM D(+)-glucose, 70 mM NaCl, 25 mM NaHCO_3_, 2.5 mM KCl, 1.25 mM NaH_2_PO_4_, 1 mM CaCl_2_ and 5 mM MgSO_4_ (carboxygenated with 5% CO_2_/95% O_2_) and followed by rapid decapitation. Brains were quickly removed and submerged in ice-cold partial sucrose solution. Coronal hippocampal slices of 200 μm thickness were cut with a vibratome (VT1200S, Leica Microsystems, Germany). The slices were incubated at 35°C for 30 min and then stored for 40 min at room temperaure for recovery.

### Measurement of protein translation

Protein translation was measured with the SUnSET assay^74^ as previously reported^76^. Acute mouse slices were incubated in artificial cerebrospinal fluid (ACSF) containing: 125 mM NaCl, 25 mM NaHCO_3_, 2.5 mM KCl, 1.25 mM NaH_2_PO_4_, 25 mM glucose, 2 mM CaCl_2_, and 1 mM MgCl_2_ (equilibrated with 95% O_2_ and 5% CO_2_). 6 μM puromycin (Merck, #P8833) was added to the slices in ACSF and allowed to incubate for 20 min at 37°C. To measure activity-dependent translation, 6 μM puromycin was added to ACSF containing 10 mM KCl^+^, which has been reported to increase neuronal activity and protein translation^76^, and allowed to incubate for 20 min. Incubating slices with 10 mM KCl did not increase puromycin incorporation (Fig. S12A-C). To block translation, 200 μM anisomycin (Sigma, #A9789) in ACSF was added to the slices 10 min before puromycin, at 37°C. Slices were then washed in ACSF, fixed with 4% PFA and 1% sucrose in PBS for 20 min and then washed with PBS. Slices were then blocked with high normal goat serum and triton-containing solution (10% normal goat serum, 0.5% Triton X-100 in 1x PBS) and stained with anti-HA antibody (1:750, Roche, #11867423001), anti-puromycin antibody (1:1000, Sigma, #MABE343), anti-NeuN antibody (1:750, SySy, #266006) and anti-S100β antibody (1:600, Dako, #Z0311) in blocking solution for 24 h at 4°C. Slices were then washed and incubated for 2 h at RT with corresponding Alexa-conjugated secondary antibodies (Invitrogen). After washing, slices were mounted and coverslipped with antifade medium (DABCO, Sigma Aldrich, 10981). Imaging was performed using a Nikon AXR confocal microscope with NIS-Elements acquisition software. A 20x objective (NA 0.80) was used to acquire z-stacks (step size of 0.80 μm, range between 11-15 μm, 2048 x 2048 pixels) with the experimenter blinded to the condition. ImageJ was used for image analysis and rendering. To measure puromycin intensity in neurons, astrocytes and neuropil, custom-written macros were used to automatically detect S100β+ and HA- or HA+ astrocytes and NeuN+ neurons, and to manually select ROIs in the neuropil, where puromycin intensity was measured. Puromycin intensity in the negative control was substracted and data were normalized to the average intensity in WT-Ctrl in the presence of puromycin (without anisomycin). To quantify CRISPR-Cas9 transduction efficiency and specificity, the same images were used and HA+ cells were manually counted and annotated as overlapping with S100β+ or NeuN+ cells.

### Quantification and statistical analysis

Data visualization and statistics were performed using R version 4.3.2 (R Core Team, 2023), R-Studio (R-Studio version ‘2023.12.0.369’) and GraphPad Prism 10.0 software. Details on the statistical tests used and n numbers for each (statistical) comparison can be found in the figure legends. Full details and precise p-values are reported in Table S3. Parametric tests were used when the data were normally distributed according to the Shapiro-Wilk test, and non-parametric tests were used when the data were not normally distributed. Data were tested for outliers using the ROUT method (Q=0.5%). When testing one condition between two groups, we used paired and unpaired Student’s t-test or the nonparametric alternatives, as indicated. Data were analyzed by a (nested) one-way analysis of variance (ANOVA) or non-parametric Kruskal-Wallis test in case one condition was tested among more than two groups, or a two-way ANOVA when more than one condition was tested. Repeated measures ANOVA was used when data were measured and statistically tested over time. A Mantel-Cox analysis was applied for the analysis of survival plots. Reported numerical values are given as mean ± SEM, and presented along with individual data points. p≤0.05 was set for significance, except for GO enrichment analysis, which used FDR<0.05 for significance. Mice were assigned to experimental groups randomly.

## Author contributions

CB-E performed stereotaxic surgeries, histology, confocal imaging, electron microscopy, immunoblotting, analysis of protein translation and mass spectrometry. MSJK performed behavioral experiments, ELISA, histology and confocal imaging. MJDVDZ performed behavioral experiments. YG performed immunoblotting. RVK performed mass spectrometry. CB-E and MSJK analysed the data and designed the figures. CFMH, MP and EH critically advised on the project and revised the manuscript. MHGV, CBE-E, MSJK and ABS conceived the project. MHGV and ABS supervised all aspects of the project and secured funding. CB-E and MHGV wrote the manuscript with input from all the other authors.

## Acknowledgements

The authors would like to thank Rolinka van der Loo and Aline Mak (VU University, Amsterdam, The Netherlands) for assistance with animal experiments, Berna Ozer (VU University Amsterdam, The Netherlands) for assistance with mass-spectrometry sample preparation, Naz Simsek (VU University Amsterdam, The Netherlands) for analysis of CRISPR/Cas9 efficiency and Rogier Min (Amsterdam UMC, Amsterdam, The Netherlands) for providing equipment and materials for preparation of acute slices and Prof. Marcela Pekna for her comments on data interpretation..

## References

1. Brandebura, A.N., Paumier, A., Onur, T.S., and Allen, N.J. (2023). Astrocyte contribution to dysfunction, risk and progression in neurodegenerative disorders. Nat. Rev. Neurosci. 24, 23–39. 10.1038/s41583-022-00641-1.

2. Badia-Soteras, A., Mak, A., Blok, T.M., Boers-Escuder, C., van den Oever, M.C., Min, R., Smit, A.B., and Verheijen, M.H.G. (2025). Astrocyte-Synapse Structural Plasticity in Neurodegenerative and Neuropsychiatric Diseases. Biol. Psychiatry. 10.1016/j.biopsych.2025.04.011.

3. Khakh, B.S., and Sofroniew, M. V (2015). Diversity of astrocyte functions and phenotypes in neural circuits. Nat. Neurosci. 18, 942–952. 10.1038/nn.4043.

4. Baldwin, K.T., Murai, K.K., and Khakh, B.S. (2024). Astrocyte morphology. Trends Cell Biol. 34, 547–565. 10.1016/j.tcb.2023.09.006.

5. Mathys, H., Boix, C.A., Akay, L.A., Xia, Z., Davila-Velderrain, J., Ng, A.P., Jiang, X., Abdelhady, G., Galani, K., Mantero, J., et al. (2024). Single-cell multiregion dissection of Alzheimer’s disease. Nature 632, 858–868. 10.1038/s41586-024-07606-7.

6. Mathys, H., Peng, Z., Boix, C.A., Victor, M.B., Leary, N., Babu, S., Abdelhady, G., Jiang, X., Ng, A.P., Ghafari, K., et al. (2023). Single-cell atlas reveals correlates of high cognitive function, dementia, and resilience to Alzheimer’s disease pathology. Cell 186, 4365–4385.e27. 10.1016/j.cell.2023.08.039.

7. Habib, N., McCabe, C., Medina, S., Varshavsky, M., Kitsberg, D., Dvir-Szternfeld, R., Green, G., Dionne, D., Nguyen, L., Marshall, J.L., et al. (2020). Disease-associated astrocytes in Alzheimer’s disease and aging. Nat. Neurosci. 23, 701–706. 10.1038/s41593-020-0624-8.

8. Escartin, C., Galea, E., Lakatos, A., O’Callaghan, J.P., Petzold, G.C., Serrano-Pozo, A., Steinhäuser, C., Volterra, A., Carmignoto, G., Agarwal, A., et al. (2021). Reactive astrocyte nomenclature, definitions, and future directions. Preprint at Nature Research, 10.1038/s41593-020-00783-4 https://doi.org/10.1038/s41593-020-00783-4.

9. Milà-Alomà, M., Brinkmalm, A., Ashton, N.J., Kvartsberg, H., Shekari, M., Operto, G., Salvadó, G., Falcon, C., Gispert, J.D., Vilor-Tejedor, N., et al. (2021). CSF Synaptic Biomarkers in the Preclinical Stage of Alzheimer Disease and Their Association With MRI and PET. Neurology 97, e2065–e2078. 10.1212/WNL.0000000000012853.

10. Fernández-Matarrubia, M., Valera-Barrero, A., Renuncio-García, M., Aguilella, M., Lage, C., López-García, S., Ocejo-Vinyals, J.G., Martínez-Dubarbie, F., Molfetta, G. Di, Pozueta-Cantudo, A., et al. (2025). Early microglial and astrocyte reactivity in preclinical Alzheimer’s disease. Alzheimer’s & Dementia 21, e70502. 10.1002/ALZ.70502.

11. Mecca, A.P., O’Dell, R.S., Sharp, E.S., Banks, E.R., Bartlett, H.H., Zhao, W., Hoffman, J.M., McDonald, J.W., Holtzman, D.M., Gordon, B.A., et al. (2022). Synaptic loss in Alzheimer’s disease measured by ^11C-UCB-J PET imaging. Alzheimer’s & Dementia 18, 1074–1085. 10.1002/alz.12438.

12. Terry, R.D., Masliah, E., Salmon, D.P., Butters, N., DeTeresa, R., Hill, R., Hansen, L.A., and Katzman, R. (1991). Physical basis of cognitive alterations in Alzheimer’s disease: synapse loss is the major correlate of cognitive impairment. Ann. Neurol. 30, 572–580. 10.1002/ana.410300410.

13. DeKosky, S.T., and Scheff, S.W. (1990). Synapse loss in frontal cortex biopsies in Alzheimer’s disease: correlation with cognitive severity. Ann. Neurol. 27, 457–464. 10.1002/ana.410270502.

14. Sperling, R.A., Aisen, P.S., Beckett, L.A., Bennett, D.A., Craft, S., Fagan, A.M., Iwatsubo, T., Jack Jr., C.R., Kaye, J., Montine, T.J., et al. (2011). Toward defining the preclinical stages of Alzheimer’s disease: Recommendations from the National Institute on Aging-Alzheimer’s Association workgroups on diagnostic guidelines for Alzheimer’s disease. Alzheimer’s & Dementia 7, 280–292. 10.1016/j.jalz.2011.03.003.

15. Carter, S.F., Herholz, K., Rosa-Neto, P., Pellerin, L., Nordberg, A., and Zimmer, E.R. (2019). Astrocyte Biomarkers in Alzheimer’s Disease. Trends Mol. Med. 25, 77–95. 10.1016/j.molmed.2018.11.006.

16. Pekny, M., and Pekna, M. (2014). Astrocyte Reactivity and Reactive Astrogliosis: Costs and Benefits. Physiol. Rev. 94, 1077–1098. 10.1152/physrev.00041.2013.

17. Kater, M.S.J., Badia-Soteras, A., van Weering, J.R.T., Smit, A.B., and Verheijen, M.H.G. (2023). Electron microscopy analysis of astrocyte-synapse interactions shows altered dynamics in an Alzheimer’s disease mouse model. Front. Cell. Neurosci. 17. 10.3389/fncel.2023.1085690.

18. Guttenplan, K.A., Weigel, M.K., Prakash, P., Wijewardhane, P.R., Hasel, P., Rufen-Blanchette, U., Münch, A.E., Blum, J.A., Fine, J., Neal, M.C., et al. (2021). Neurotoxic reactive astrocytes induce cell death via saturated lipids. Nature 599, 102–107. 10.1038/s41586-021-03960-y.

19. Smit, T., Deshayes, N.A.C., Borchelt, D.R., Kamphuis, W., Middeldorp, J., and Hol, E.M. (2021). Reactive astrocytes as treatment targets in Alzheimer’s disease — Systematic review of studies using the APPswePS1dE9 mouse model. Glia 69, 1852–1881. 10.1002/glia.23981.

20. Jankowsky, J.L., Slunt, H.H., Gonzales, V., Jenkins, N.A., Copeland, N.G., and Borchelt, D.R. (2004). APP processing and amyloid deposition in mice haplo-insufficient for presenilin 1. Neurobiol. Aging 25, 885–892. 10.1016/j.neurobiolaging.2003.09.008.

21. Desclaux, M., Perrin, F.E., Do-Thi, A., Prieto-Cappellini, M., Gimenez y Ribotta, M., Mallet, J., and Privat, A. (2015). Lentiviral-mediated silencing of glial fibrillary acidic protein and vimentin promotes anatomical plasticity and functional recovery after spinal cord injury. J. Neurosci. Res. 93, 43–55. 10.1002/jnr.23468.

22. Wilhelmsson, U., Li, L., Pekna, M., Berthold, C.-H., Blom, S., Eliasson, C., Renner, O., Bushong, E., Ellisman, M., Morgan, T.E., et al. (2004). Absence of glial fibrillary acidic protein and vimentin prevents hypertrophy of astrocytic processes and improves post-traumatic regeneration. The Journal of Neuroscience 24, 5016–5021. 10.1523/jneurosci.0820-04.2004.

23. Kamphuis, W., Kooijman, L., Orre, M., Stassen, O., Pekny, M., and Hol, E. (2015). GFAP and vimentin deficiency alters gene expression in astrocytes and microglia in wild-type mice and changes the transcriptional response of reactive glia in Alzheimer’s disease model mice. Glia 63, 1036–1056. 10.1002/glia.22704.

24. Liu, Z., Li, Y., Cui, Y., Roberts, C., Lu, M., Wilhelmsson, U., Pekny, M., and Chopp, M. (2014). Beneficial effects of GFAP/Vimentin reactive astrocytes for axonal remodeling and motor behavioral recovery in mice after stroke. Glia 62, 2022–2033. 10.1002/glia.22723.

25. Pekny, M., Levéen, P., Pekna, M., Eliasson, C., Berthold, C.H., Westermark, B., and Betsholtz, C. (1995). Mice lacking glial fibrillary acidic protein display astrocytes devoid of intermediate filaments but develop and reproduce normally. EMBO J. 14, 1590–1598. 10.1002/j.1460-2075.1995.tb07147.x.

26. Wilhelmsson, U., Pozo-Rodrigalvarez, A., Kalm, M., de Pablo, Y., Widestrand, Å., Pekna, M., and Pekny, M. (2019). The role of GFAP and vimentin in learning and memory. Journal of Biological Chemistry 400, 1147–1156. 10.1515/hsz-2019-0199.

27. Pekny, M., Johansson, C.B., Eliasson, C., Stakeberg, J., Wallén, A., Perlmann, T., Lendahl, U., Betsholtz, C., Berthold, C.H., and Frisén, J. (1999). Abnormal reaction to central nervous system injury in mice lacking glial fibrillary acidic protein and vimentin. Journal of Cell Biology 145, 503–514. 10.1083/jcb.145.3.503.

28. Cho, M.-I., and others (2005). Enhanced synaptic recovery after injury in the absence of vimentin and GFAP. J. Neurocytol.

29. Li, L., Lundkvist, A., Andersson, D., Wilhelmsson, U., Nagai, N., Pardo, A.C., Nodin, C., Ståhlberg, A., Aprico, K., Larsson, K., et al. (2008). Protective role of reactive astrocytes in brain ischemia. Journal of Cerebral Blood Flow and Metabolism 28, 468–481. 10.1038/SJ.JCBFM.9600546/ASSET/B7BF00DE-877E-4AC8-9162-DDED93CB0245/ASSETS/IMAGES/LARGE/10.1038_SJ.JCBFM.9600546-FIG9.JPG.

30. Ding, M., Eliasson, C., Betsholtz, C., Hamberger, A., and Pekny, M. (1998). Altered taurine release following hypotonic stress in astrocytes from mice deficient for GFAP and vimentin. Molecular Brain Research 62, 77–81. 10.1016/S0169-328X(98)00240-X.

31. Lundkvist, A., Reichenbach, A., Betsholtz, C., Carmeliet, P., Wolburg, H., and Pekny, M. (2004). Under stress, the absence of intermediate filaments from Müller cells in the retina has structural and functional consequences. J. Cell Sci. 117, 3481–3488. 10.1242/JCS.01221.

32. Menet, V., Prieto, M., Privat, A., and Giménez y Ribotta, M. (2003). Axonal plasticity and functional recovery after spinal cord injury in mice deficient in both glial fibrillary acidic protein and vimentin genes. Proc. Natl. Acad. Sci. U. S. A. 100, 8999–9004. 10.1073/PNAS.1533187100/ASSET/1A2581E8-077C-4BC5-99D4-492D2E30DA16/ASSETS/GRAPHIC/PQ1533187006.JPEG.

33. Kater, M.S.J., Huffels, C.F.M., Oshima, T., Renckens, N.S., Middeldorp, J., Boddeke, E.W.G.M., Smit, A.B., Eggen, B.J.L., Hol, E.M., and Verheijen, M.H.G. (2023). Prevention of microgliosis halts early memory loss in a mouse model of Alzheimer’s disease. Brain Behav. Immun. 107, 225–241. 10.1016/j.bbi.2022.10.009.

34. Ortinski, P.I., Dong, J., Mungenast, A., Yue, C., Takano, H., Watson, D.J., Haydon, P.G., and Coulter, D.A. (2010). Selective induction of astrocytic gliosis generates deficits in neuronal inhibition. Nat. Neurosci. 13, 584–591. 10.1038/nn.2535.

35. Kulijewicz-Nawrot, M., Syková, E., Chvátal, A., Verkhratsky, A., and Rodríguez, J.J. (2013). Astrocytes and glutamate homoeostasis in Alzheimer’s disease: a decrease in glutamine synthetase, but not in glutamate transporter-1, in the prefrontal cortex. ASN Neuro 5, 273–282. 10.1042/an20130017.

36. Le Prince, G., Delaere, P., Fages, C., Lefrançois, T., Touret, M., Salanon, M., and Tardy, M. (1995). Glutamine synthetase (GS) expression is reduced in senile dementia of the Alzheimer type. Neurochem. Res. 20, 859–862. 10.1007/BF00969698/METRICS.

37. Robinson, S.R. (2000). Neuronal expression of glutamine synthetase in Alzheimer’s disease indicates a profound impairment of metabolic interactions with astrocytes. Neurochem. Int. 36, 471–482. 10.1016/S0197-0186(99)00150-3.

38. Olabarria, M., Noristani, H.N., Verkhratsky, A., and Rodríguez, J.J. (2011). Age-dependent decrease in glutamine synthetase expression in the hippocampal astroglia of the triple transgenic Alzheimer’s disease mouse model: mechanism for deficient glutamatergic transmission? 10.1186/1750-1326-6-55.

39. Pallari, H.-M., and Eriksson, J.E. (2006). Intermediate filaments as signaling platforms. Sci. STKE 2006, pe53. 10.1126/stke.3662006pe53.

40. Terzi, F., Henrion, D., Colucci-Guyon, E., Federici, P., Babinet, C., Levy, B.I., Briand, P., and Friedlander, G. (1997). Reduction of renal mass is lethal in mice lacking vimentin. Role of endothelin-nitric oxide imbalance. J. Clin. Invest. 100, 1520–1528. 10.1172/JCI119675.

41. Végh, M.J., Heldring, C.M., Kamphuis, W., Hijazi, S., Timmerman, A.J., Li, K.W., van Nierop, P., Mansvelder, H.D., Hol, E.M., Smit, A.B., et al. (2014). Reducing hippocampal extracellular matrix reverses early memory deficits in a mouse model of Alzheimer’s disease. Acta Neuropathol. Commun. 2. 10.1186/s40478-014-0076-z.

42. Hijazi, S., Heistek, T.S., Scheltens, P., Neumann, U., Shimshek, D.R., Mansvelder, H.D., Smit, A.B., and van Kesteren, R.E. (2019). Early restoration of parvalbumin interneuron activity prevents memory loss and network hyperexcitability in a mouse model of Alzheimer’s disease. Mol. Psychiatry 25, 3380–3398. 10.1038/s41380-019-0483-4.

43. Wang, T., Chen, Y., Zou, Y., Pang, Y., He, X., Chen, Y., Liu, Y., Feng, W., Zhang, Y., Li, Q., et al. (2022). Locomotor Hyperactivity in the Early-Stage Alzheimer’s Disease-like Pathology of APP/PS1 Mice: Associated with Impaired Polarization of Astrocyte Aquaporin 4. Aging Dis. 13, 1504–1522. 10.14336/AD.2022.0219.

44. Végh, M.J., Heldring, C.M., Kamphuis, W., Hijazi, S., Timmerman, A.J., Li, K.W., van Nierop, P., Mansvelder, H.D., Hol, E.M., Smit, A.B., et al. (2014). Reducing hippocampal extracellular matrix reverses early memory deficits in a mouse model of Alzheimer’s disease. Acta Neuropathol. Commun. 2, 76. 10.1186/s40478-014-0076-z.

45. Wyss-Coray, T., Loike, J.D., Brionne, T.C., Lu, E., Anankov, R., Yan, F., Silverstein, S.C., and Husemann, J. (2003). Adult mouse astrocytes degrade amyloid-β in vitro and in situ. Nat. Med. 9, 453–457. 10.1038/nm838.

46. Kraft, A.W., Hu, X., Yoon, H., Yan, P., Xiao, Q., Wang, Y., Gil, S.C., Brown, J., Wilhelmsson, U., Restivo, J.L., et al. (2013). Attenuating astrocyte activation accelerates plaque pathogenesis in APP/PS1 mice. The FASEB Journal 27, 187–198. 10.1096/fj.12-208660.

47. Liu, C.-C., Zhao, N., Fu, Y., Wang, N., Linares, C., Tsai, C.-W., and Bu, G. (2017). ApoE4 Accelerates Early Seeding of Amyloid Pathology. Neuron 96, 1024–1032.e3. 10.1016/j.neuron.2017.11.013.

48. Ceyzériat, K., Ben Haim, L., Denizot, A., Pommier, D., Matos, M., Guillemaud, O., Palomares, M.-A., Abjean, L., Petit, F., Gipchtein, P., et al. (2018). Modulation of astrocyte reactivity improves functional deficits in mouse models of Alzheimer’s disease. Acta Neuropathol. Commun. 6, 104. 10.1186/s40478-018-0606-1.

49. Diaz-Castro, B., Bernstein, A.M., Coppola, G., Sofroniew, M. V, and Khakh, B.S. (2021). Molecular and functional properties of cortical astrocytes during peripherally induced neuroinflammation. Cell Rep. 36, 109508. 10.1016/j.celrep.2021.109508.

50. Mathys, H., Davila-Velderrain, J., Peng, Z., Gao, F., Mohammadi, S., Young, J.Z., Menon, M., He, L., Abdurrob, F., Jiang, X., et al. (2019). Single-cell transcriptomic analysis of Alzheimer’s disease. Nature 570, 332–337. 10.1038/s41586-019-1195-2.

51. Sadick, J.S., O’Dea, M.R., Hasel, P., Dykstra, T., Faustin, A., and Liddelow, S.A. (2022). Astrocytes and oligodendrocytes undergo subtype-specific transcriptional changes in Alzheimer’s disease. Neuron 110, 1788–1805.e10. 10.1016/j.neuron.2022.03.008.

52. Diaz-Castro, B., Gangwani, M.R., Yu, X., Coppola, G., and Khakh, B.S. (2019). Astrocyte molecular signatures in Huntington’s disease. Sci. Transl. Med. 11, eaaw8546. 10.1126/scitranslmed.aaw8546.

53. Yu, X., Nagai, J., Marti-Solano, M., Soto, J.S., Coppola, G., Babu, M.M., and Khakh, B.S. (2020). Context-Specific Striatal Astrocyte Molecular Responses Are Phenotypically Exploitable. Neuron 108, 1146–1162.e10. 10.1016/j.neuron.2020.09.021.

54. Pekny, M., Pekna, M., Messing, A., Steinhäuser, C., Lee, J.M., Parpura, V., Hol, E.M., Sofroniew, M. V., and Verkhratsky, A. (2016). Astrocytes: a central element in neurological diseases. Acta Neuropathol. 131, 323–345. 10.1007/S00401-015-1513-1.

55. Soto, J.S., Jami-Alahmadi, Y., Chacon, J., Moye, S.L., Diaz-Castro, B., Wohlschlegel, J.A., and Khakh, B.S. (2023). Astrocyte–neuron subproteomes and obsessive–compulsive disorder mechanisms. Nature 616, 764–773. 10.1038/s41586-023-05927-7.

56. Koopmans, F., Li, K.W., Klaassen, R. V, and Smit, A.B. (2023). MS-DAP platform for downstream data analysis of label-free proteomics uncovers optimal workflows in benchmark data sets and increased sensitivity in analysis of Alzheimer’s biomarker data. J. Proteome Res. 22, 374–386. 10.1021/acs.jproteome.2c00513.

57. Dejanovic, B., Wu, T., Tsai, M.C., Graykowski, D., Gandham, V.D., Rose, C.M., Bakalarski, C.E., Ngu, H., Wang, Y., Pandey, S., et al. (2022). Complement C1q-dependent excitatory and inhibitory synapse elimination by astrocytes and microglia in Alzheimer’s disease mouse models. Nat. Aging 2, 837–850. 10.1038/s43587-022-00281-1.

58. Hong, S., Beja-Glasser, V.F., Nfonoyim, B.M., Frouin, A., Li, S., Ramakrishnan, S., Merry, K.M., Shi, Q., Rosenthal, A., Barres, B.A., et al. (2016). Complement and microglia mediate early synapse loss in Alzheimer mouse models. Science (1979). 352, 712–716. 10.1126/science.aad8373.

59. Tzioras, M., Daniels, M.J.D., Davies, C., Baxter, P., King, D., McKay, S., Varga, B., Popovic, K., Hernandez, M., Stevenson, A.J., et al. (2023). Human astrocytes and microglia show augmented ingestion of synapses in Alzheimer’s disease via MFG-E8. Cell Rep. Med. 4, 101175. 10.1016/j.xcrm.2023.101175.

60. Jeppesen, D.K., Fenix, A.M., Franklin, J.L., Higginbotham, J.N., Zhang, Q., Zimmerman, L.J., Liebler, D.C., Ping, J., Liu, Q., Evans, R., et al. (2019). Reassessment of Exosome Composition. Cell 177, 428–445.e18. 10.1016/j.cell.2019.02.029.

61. Lasič, E., Trkov Bobnar, S., Wilhelmsson, U., de Pablo, Y., Pekny, M., Zorec, R., and Stenovec, M. (2020). Nestin affects fusion pore dynamics in mouse astrocytes. Acta Physiol. (Oxf). 228. 10.1111/APHA.13399.

62. Potokar, M., Kreft, M., Li, L., Daniel Andersson, J., Pangršič, T., Chowdhury, H.H., Pekny, M., and Zorec, R. (2007). Cytoskeleton and Vesicle Mobility in Astrocytes. Traffic 8, 12–20. 10.1111/j.1600-0854.2006.00509.x.

63. Vardjan, N., Gabrijel, M., Potokar, M., Švajger, U., Kreft, M., Jeras, M., de Pablo, Y., Faiz, M., Pekny, M., and Zorec, R. (2012). IFN-γ-induced increase in the mobility of MHC class II compartments in astrocytes depends on intermediate filaments. J. Neuroinflammation 9, 144. 10.1186/1742-2094-9-144.

64. Ashburner, M., Ball, C.A., Blake, J.A., Botstein, D., Butler, H., Cherry, J.M., Davis, A.P., Dolinski, K., Dwight, S.S., Eppig, J.T., et al. (2000). Gene Ontology: tool for the unification of biology. Nature Genetics 2000 25:1 25, 25–29. 10.1038/75556.

65. Wang, Z., Qiu, H., He, J., Liu, L., Xue, W., Fox, A., Tickner, J., and Xu, J. (2020). The emerging roles of hnRNPK. J. Cell. Physiol. 235, 1995–2008. 10.1002/jcp.29186.

66. Kotah, J.M., Kater, M.S.J., Brosens, N., Lesuis, S.L., Tandari, R., Blok, T.M., Marchetto, L., Yusaf, E., Koopmans, F.T.W., Smit, A.B., et al. (2023). Early-life stress and amyloidosis in mice share pathogenic pathways involving synaptic mitochondria and lipid metabolism. Alzheimer’s & Dementia. 10.1002/alz.13569.

67. Ahmad, F., Singh, K., Das, D., Gowaikar, R., Shaw, E., Ramachandran, A., Rupanagudi, K.V., Kommaddi, R.P., Bennett, D.A., and Ravindranath, V. (2017). Reactive Oxygen Species-Mediated Loss of Synaptic Akt1 Signaling Leads to Deficient Activity-Dependent Protein Translation Early in Alzheimer’s Disease. Antioxid. Redox Signal. 27, 1269–1280. 10.1089/ars.2016.6860.

68. Sharma, K., Schmitt, S., Bergner, C.G., Tyanova, S., Kannaiyan, N., Manrique-Hoyos, N., Kongi, K., Cantuti, L., Hanisch, U.-K., Philips, M.-A., et al. (2015). Cell type– and brain region–resolved mouse brain proteome. Nat. Neurosci. 18, 1819–1831. 10.1038/nn.4160.

69. Sakers, K., Lake, A.M., Khazanchi, R., Ouwenga, R., Vasek, M.J., Dani, A., and Dougherty, J.D. (2017). Astrocytes locally translate transcripts in their peripheral processes. Proceedings of the National Academy of Sciences 114, E3830–E3838. 10.1073/pnas.1617782114.

70. Mazaré, N., Oudart, M., Moulard, J., Cheung, G., Tortuyaux, R., Mailly, P., Mazaud, D., Bemelmans, A.-P., Boulay, A.-C., Blugeon, C., et al. (2020). Local Translation in Perisynaptic Astrocytic Processes Is Specific and Changes after Fear Conditioning. Cell Rep. 32, 108076. 10.1016/j.celrep.2020.108076.

71. López-Erauskin, J., Tadokoro, T., Baughn, M.W., Myers, B., McAlonis-Downes, M., Chillon-Marinas, C., Asiaban, J.N., Artates, J., Bui, A.T., Vetto, A.P., et al. (2018). ALS/FTD-Linked Mutation in FUS Suppresses Intra-axonal Protein Synthesis and Drives Disease Without Nuclear Loss-of-Function of FUS. Neuron 100, 816–830.e7. 10.1016/j.neuron.2018.09.044.

72. Sahadevan, S., Hembach, K.M., Tantardini, E., Pérez-Berlanga, M., Hruska-Plochan, M., Megat, S., Weber, J., Schwarz, P., Dupuis, L., Robinson, M.D., et al. (2021). Synaptic FUS accumulation triggers early misregulation of synaptic RNAs in a mouse model of ALS. Nat. Commun. 12, 3027. 10.1038/s41467-021-23188-8.

73. Li, W., Dasgupta, A., Yang, K., Wang, S., Hemandhar-Kumar, N., Chepyala, S.R., Yarbro, J.M., Hu, Z., Salovska, B., Fornasiero, E.F., et al. (2025). Turnover atlas of proteome and phosphoproteome across mouse tissues and brain regions. Cell 188, 2267–2287.e21. 10.1016/j.cell.2025.02.021.

74. Schmidt, E.K., Clavarino, G., Ceppi, M., and Pierre, P. (2009). SUnSET, a nonradioactive method to monitor protein synthesis. Nat. Methods 6, 275–277. 10.1038/nmeth.1314.

75. Plaska, C.R., Jacobs, T., Bruno, D., Lee, S.H., Imbimbo, B.P., Osorio, R., Benedet, A.L., Ashton, N.J., Zetterberg, H., and Pomara, N. (2026). Sex-specific associations between astrocytic reactivity and cognitive decline in unimpaired elderly. bioRxiv, 2026.01.30.702902. 10.64898/2026.01.30.702902.

76. Sapkota, D., Kater, M., Sakers, K., Nygaard, K., Liu, Y., Koester, S., Fass, S., Lake, S., Khazanchi, R., Khankan, R., et al. (2022). Activity-dependent translation dynamically alters the proteome of the perisynaptic astrocyte process. Cell Rep. 41. 10.1016/j.celrep.2022.111474.

77. Chang, L., and Goldman, R.D. (2004). Intermediate filaments mediate cytoskeletal crosstalk. Nature Reviews Molecular Cell Biology 2004 5:8 *5*, 601–613. 10.1038/nrm1438.

78. Hol, E.M., and Pekny, M. (2015). Glial fibrillary acidic protein (GFAP) and the astrocyte intermediate filament system in diseases of the central nervous system. Curr. Opin. Cell Biol. 32, 121–130. 10.1016/j.ceb.2015.02.004.

79. Eliasson, C., Sahlgren, C., Berthold, C.H., Stakeberg, J., Celis, J.E., Betsholtz, C., Eriksson, J.E., and Pekny, M. (1999). Intermediate filament protein partnership in astrocytes. Journal of Biological Chemistry 274, 23996–24006. 10.1074/jbc.274.34.23996.

80. Potokar, M., Morita, M., Wiche, G., and Jorgačevski, J. (2020). The Diversity of Intermediate Filaments in Astrocytes. Cells 9. 10.3390/cells9071604.

81. Macauley, S.L., Pekny, M., and Sands, M.S. (2011). The role of attenuated astrocyte activation in infantile neuronal ceroid lipofuscinosis. J. Neurosci. 31, 15575–15585. 10.1523/JNEUROSCI.3579-11.2011.

82. Liddelow, S.A., Marsh, S.E., and Stevens, B. (2020). Microglia and Astrocytes in Disease: Dynamic Duo or Partners in Crime? Trends Immunol. 41, 820–835. 10.1016/J.IT.2020.07.006.

83. Salter, M.W., and Stevens, B. (2017). Microglia emerge as central players in brain disease. Nat. Med. 23, 1018–1027. 10.1038/NM.4397.

84. Sano, F., Shigetomi, E., Shinozaki, Y., Tsuzukiyama, H., Saito, K., Mikoshiba, K., Horiuchi, H., Cheung, D.L., Nabekura, J., Sugita, K., et al. (2021). Reactive astrocyte-driven epileptogenesis is induced by microglia initially activated following status epilepticus. JCI Insight 6. 10.1172/JCI.INSIGHT.135391.

85. Gotoh, M., Miyamoto, Y., and Ikeshima-Kataoka, H. (2023). Astrocytic Neuroimmunological Roles Interacting with Microglial Cells in Neurodegenerative Diseases. Int. J. Mol. Sci. 24. 10.3390/IJMS24021599.

86. Osborn, L.M., Kamphuis, W., Wadman, W.J., and Hol, E.M. (2016). Astrogliosis: an integral player in the pathogenesis of Alzheimer’s disease. Prog. Neurobiol. 144, 121–141. 10.1016/j.pneurobio.2016.01.001.

87. Fernández-Matarrubia, M., and others (2025). Early microglial and astrocyte reactivity in preclinical Alzheimer’s disease. Alzheimer’s & Dementia 21, e70502. 10.1002/alz.70502.

88. Lee, E., Jung, Y.-J., Park, Y.R., Lim, S., Choi, Y.-J., Lee, S.Y., Kim, C.H., Mun, J.Y., and Chung, W.-S. (2022). A distinct astrocyte subtype in the aging mouse brain characterized by impaired protein homeostasis. Nat. Aging 2, 726–741. 10.1038/s43587-022-00257-1.

89. Serrano-Pozo, A., Li, H., Li, Z., Muñoz-Castro, C., Jaisa-aad, M., Healey, M.A., Welikovitch, L.A., Jayakumar, R., Bryant, A.G., Noori, A., et al. (2024). Astrocyte transcriptomic changes along the spatiotemporal progression of Alzheimer’s disease. Nat. Neurosci. 27, 2384–2400. 10.1038/s41593-024-01791-4.

90. Johnson, E.C.B., Dammer, E.B., Duong, D.M., Yin, L., Thambisetty, M., Troncoso, J.C., Lah, J.J., Levey, A.I., and Seyfried, N.T. (2018). Deep proteomic network analysis of Alzheimer’s disease brain reveals alterations in RNA binding proteins and RNA splicing associated with disease. Mol. Neurodegener. 13, 52. 10.1186/s13024-018-0282-4.

91. Seyfried, N.T., Dammer, E.B., Swarup, V., Nandakumar, D., Duong, D.M., Yin, L., Deng, Q., Nguyen, T., Hales, C.M., Wingo, T., et al. (2017). A Multi-network Approach Identifies Protein-Specific Co-expression in Asymptomatic and Symptomatic Alzheimer’s Disease. Cell Syst. 4, 60–72.e4. 10.1016/j.cels.2016.11.006.

92. Johnson, E.C.B., Dammer, E.B., Duong, D.M., Ping, L., Zhou, M., Yin, L., Higginbotham, L.A., Guajardo, A., White, B., Troncoso, J.C., et al. (2020). Large-scale proteomic analysis of Alzheimer’s disease brain and cerebrospinal fluid reveals early changes in energy metabolism associated with microglia and astrocyte activation. Nat. Med. 26, 769–780. 10.1038/s41591-020-0815-6.

93. Boulay, A.-C., Saubaméa, B., Adam, N., Chasseigneaux, S., Mazaré, N., Gilbert, A., Bahin, M., Bastianelli, L., Blugeon, C., Perrin, S., et al. (2017). Translation in astrocyte distal processes sets molecular heterogeneity at the gliovascular interface. Cell Discov. 3, 17005. 10.1038/celldisc.2017.5.

94. Barton, S.K., Gregory, J.M., Chandran, S., and Turner, B.J. (2019). Could an Impairment in Local Translation of mRNAs in Glia be Contributing to Pathogenesis in ALS? Front. Mol. Neurosci. Volume 12-2019. 10.3389/fnmol.2019.00124.

95. Higashimori, H., Morel, L., Huth, J., Lindemann, L., Dulla, C., Taylor, A., Freeman, M., and Yang, Y. (2013). Astroglial FMRP-dependent translational down-regulation of mGluR5 underlies glutamate transporter GLT1 dysregulation in the fragile X mouse. Hum. Mol. Genet. 22, 2041–2054. 10.1093/hmg/ddt055.

96. Mattsson, N., and others (2016). Cerebrospinal fluid tau, neurogranin, and neurofilament light in Alzheimer’s disease. EMBO Mol. Med. 8, 1184–1196. 10.15252/emmm.201606540.

97. De Strooper, B., and Karran, E. (2016). The Cellular Phase of Alzheimer’s Disease. Cell 164, 603–615. 10.1016/j.cell.2015.12.056.

98. Langstrom, N.S., Anderson, J.P., Lindroos, H.G., Winbland, B., and Wallace, W.C. (1989). Alzheimer’s disease-associated reduction of polysomal mRNA translation. Molecular Brain Research 5, 259–269. 10.1016/0169-328X(89)90060-0.

99. Lim, D., Tapella, L., Dematteis, G., Genazzani, A.A., Corazzari, M., and Verkhratsky, A. (2023). The endoplasmic reticulum stress and unfolded protein response in Alzheimer’s disease: A calcium dyshomeostasis perspective. Ageing Res. Rev. 87, 101914. 10.1016/j.arr.2023.101914.

100. Elder, M.K., Erdjument-Bromage, H., Oliveira, M.M., Mamcarz, M., Neubert, T.A., and Klann, E. (2021). Age-dependent shift in the de novo proteome accompanies pathogenesis in an Alzheimer’s disease mouse model. Commun. Biol. 4, 823. 10.1038/s42003-021-02324-6.

101. Butterfield, D.A., and Halliwell, B. (2019). Oxidative stress, dysfunctional glucose metabolism and Alzheimer disease. Nat. Rev. Neurosci. 20, 148–160. 10.1038/s41583-019-0132-6.

102. Chang, R.C.C., Wong, A.K.Y., Ng, H.-K., and Hugon, J. (2002). Phosphorylation of eukaryotic initiation factor-2α (eIF2α) is associated with neuronal degeneration in Alzheimer’s disease. Neuroreport 13.

103. Nijholt, D.A.T., van Haastert, E.S., Rozemuller, A.J.M., Scheper, W., and Hoozemans, J.J.M. (2012). The unfolded protein response is associated with early tau pathology in the hippocampus of tauopathies. J. Pathol. 226, 693–702. 10.1002/path.3969.

104. Ma, T., Trinh, M.A., Wexler, A.J., Bourbon, C., Gatti, E., Pierre, P., Cavener, D.R., and Klann, E. (2013). Suppression of eIF2α kinases alleviates Alzheimer’s disease–related plasticity and memory deficits. Nat. Neurosci. 16, 1299–1305. 10.1038/nn.3486.

105. Ryan, T.J., Roy, D.S., Pignatelli, M., Arons, A., and Tonegawa, S. (2015). Engram cells retain memory under retrograde amnesia. Science (1979). 348, 1007–1013. 10.1126/SCIENCE.AAA5542/SUPPL_FILE/RYAN-SM.PDF.

106. Williamson, M.R., and others (2024). Learning-associated astrocyte ensembles regulate memory recall. Nature. 10.1038/s41586-024-08170-w.

107. Sun, W., Liu, Z., Jiang, X., Chen, M.B., Dong, H., Liu, J., Südhof, T.C., and Quake, S.R. (2024). Spatial transcriptomics reveal neuron–astrocyte synergy in long-term memory. Nature 627, 374–381. 10.1038/s41586-023-07011-6.

108. Badia-Soteras, A., Heistek, T.S., Kater, M.S.J., Mak, A., Negrean, A., van den Oever, M.C., Mansvelder, H.D., Khakh, B.S., Min, R., Smit, A.B., et al. (2023). Retraction of Astrocyte Leaflets From the Synapse Enhances Fear Memory. Biol. Psychiatry 94, 226–238. 10.1016/j.biopsych.2022.10.013.

109. Sharma, V., Oliveira, M.M., Sood, R., Khlaifia, A., Lou, D., Hooshmandi, M., Hung, T.-Y., Mahmood, N., Reeves, M., Ho-Tieng, D., et al. (2023). mRNA translation in astrocytes controls hippocampal long-term synaptic plasticity and memory. Proceedings of the National Academy of Sciences 120, e2308671120. 10.1073/pnas.2308671120.

110. Vanderweyde, T., Yu, H., Varnum, M., Liu-Yesucevitz, L., Citro, A., Ikezu, T., Duff, K., and Wolozin, B. (2012). Contrasting Pathology of the Stress Granule Proteins TIA-1 and G3BP in Tauopathies. Journal of Neuroscience 32, 8270–8283. 10.1523/JNEUROSCI.1592-12.2012.

111. Maziuk, B.F., Apicco, D.J., Cruz, A.L., Jiang, L., Ash, P.E.A., da Rocha, E.L., Zhang, C., Yu, W.H., Leszyk, J., Abisambra, J.F., et al. (2018). RNA binding proteins co-localize with small tau inclusions in tauopathy. Acta Neuropathol. Commun. 6, 71. 10.1186/s40478-018-0574-5.

112. Hoek, K.S., Kidd, G.J., Carson, J.H., and Smith, R. (1998). hnRNP A2 Selectively Binds the Cytoplasmic Transport Sequence of Myelin Basic Protein mRNA. Biochemistry 37, 7021–7029. 10.1021/bi9800247.

113. Traub, P., and Nelson, W.J. (1982). Interaction of the intermediate filament protein vimentin with ribosomal subunits and ribosomal RNA in vitro. Mol. Biol. Rep. 8, 239–247. 10.1007/BF00776586.

114. Kim, S., Kellner, J., Lee, C.-H., and Coulombe, P.A. (2007). Interaction between the keratin cytoskeleton and eEF1Bγ affects protein synthesis in epithelial cells. Nat. Struct. Mol. Biol. 14, 982–983. 10.1038/nsmb1301.

115. Zhang, Y., and others (2022). Host cytoskeletal vimentin serves as a structural organizer of ribonucleoprotein complexes: interactions with ribosomal and RNA-binding proteins. Proceedings of the National Academy of Sciences 119, e2113909119. 10.1073/pnas.2113909119.

116. Challa, A.A., and Stefanovic, B. (2011). A Novel Role of Vimentin Filaments: Binding and Stabilization of Collagen mRNAs. Mol. Cell. Biol. 31, 3773–3789. 10.1128/MCB.05263-11.

117. González-Reyes, R.E., Nava-Mesa, M.O., Vargas-Sánchez, K., Ariza-Salamanca, D., and Mora-Muñoz, L. (2017). Involvement of astrocytes in Alzheimer’s Disease from a neuroinflammatory and oxidative stress perspective. Front. Mol. Neurosci. 10, 427. 10.3389/fnmol.2017.00427.

118. Pérez-Sala, D., and Quinlan, R.A. (2024). The redox-responsive roles of intermediate filaments in cellular stress detection, integration and mitigation. Curr. Opin. Cell Biol. 86, 102283. 10.1016/j.ceb.2023.102283.

119. De Pablo, Y., Nilsson, M., Pekna, M., and Pekny, M. (2013). Intermediate filaments are important for astrocyte response to oxidative stress induced by oxygen-glucose deprivation and reperfusion. Histochem. Cell Biol. 140, 81–91. 10.1007/S00418-013-1110-0.

120. Kim, S., Wong, P., and Coulombe, P.A. (2006). A keratin cytoskeletal protein regulates protein synthesis and epithelial cell growth. Nature 2006 441:7091 441, 362–365. 10.1038/nature04659.

121. Mohanasundaram, P., Coelho-Rato, L.S., Modi, M.K., Urbanska, M., Lautenschläger, F., Cheng, F., and Eriksson, J.E. (2022). Cytoskeletal vimentin regulates cell size and autophagy through mTORC1 signaling. PLoS Biol. 20, e3001737-.

122. Morrow, C.S., Porter, T.J., Xu, N., Arndt, Z.P., Ako-Asare, K., Heo, H.J., Thompson, E.A.N., and Moore, D.L. (2020). Vimentin Coordinates Protein Turnover at the Aggresome during Neural Stem Cell Quiescence Exit. Cell Stem Cell 26, 558–568.e9. 10.1016/j.stem.2020.01.018.

123. Pajares, M.A., and Pérez-Sala, D. (2024). Type III intermediate filaments in redox interplay: key role of the conserved cysteine residue. Biochem. Soc. Trans. 52, 849–860. 10.1042/BST20231059.

124. Pérez-Sala, D., and Zorrilla, S. (2025). Versatility of vimentin assemblies: From filaments to biomolecular condensates and back. Eur. J. Cell Biol. 104, 151487. 10.1016/j.ejcb.2025.151487.

125. Chatterjee, S., Panda, A.C., Berwal, S.K., Sreejith, R.K., Ritvika, C., Seshadri, V., and Pal, J.K. (2013). Vimentin is a component of a complex that binds to the 5′-UTR of human heme-regulated eIF2α kinase mRNA and regulates its translation. FEBS Lett. 587, 474–480. 10.1016/j.febslet.2013.01.013.

126. Chen, K., Morizawa, Y.M., Nuriel, T., Al-Dalahmah, O., Xie, Z., and Yang, G. (2025). Selective removal of astrocytic PERK protects against glymphatic impairment and decreases toxic aggregation of β-amyloid and tau. Neuron 113, 2438–2454.e6. 10.1016/j.neuron.2025.04.027.

127. Smith, H.L., Freeman, O.J., Butcher, A.J., Holmqvist, S., Humoud, I., Schätzl, T., Hughes, D.T., Verity, N.C., Swinden, D.P., Hayes, J., et al. (2020). Astrocyte Unfolded Protein Response Induces a Specific Reactivity State that Causes Non-Cell-Autonomous Neuronal Degeneration. Neuron 105, 855–866.e5. 10.1016/j.neuron.2019.12.014.

128. Batenburg, K.L., Kasri, N.N., Heine, V.M., and Scheper, W. (2022). Intraneuronal tau aggregation induces the integrated stress response in astrocytes. J. Mol. Cell Biol. 14, mjac071. 10.1093/jmcb/mjac071.

129. Tapella, L., Dematteis, G., Moro, M., Pistolato, B., Tonelli, E., Vanella, V.V., Giustina, D., La Forgia, A., Restelli, E., Barberis, E., et al. (2022). Protein synthesis inhibition and loss of homeostatic functions in astrocytes from an Alzheimer’s disease mouse model: a role for ER-mitochondria interaction. Cell Death Dis. 13, 878. 10.1038/s41419-022-05324-4.

130. Wilhelmsson, U., Stillemark-Billton, P., Borén, J., and Pekny, M. (2019). Vimentin is required for normal accumulation of body fat. Biol. Chem. 400, 1157–1162. 10.1515/HSZ-2019-0170.

131. Ridge, K.M., Eriksson, J.E., Pekny, M., and Goldman, R.D. (2022). Roles of vimentin in health and disease. Genes Dev. 36, 391–407. 10.1101/gad.349358.122.

132. Schnitzer, J., Franke, W.W., and Schachner, M. (1981). Immunocytochemical demonstration of vimentin in astrocytes and ependymal cells of developing and adult mouse nervous system. Journal of Cell Biology 90, 435–447. 10.1083/JCB.90.2.435.

133. Mamber, C., Kamphuis, W., Haring, N.L., Peprah, N., Middeldorp, J., and Hol, E.M. (2012). GFAPδ Expression in Glia of the Developmental and Adolescent Mouse Brain. PLoS One 7, e52659. 10.1371/JOURNAL.PONE.0052659.

134. Perez-Nievas, B.G., Stein, T.D., Tai, H.C., Dols-Icardo, O., Scotton, T.C., Barroeta-Espar, I., Fernandez-Carballo, L., De Munain, E.L., Perez, J., Marquie, M., et al. (2013). Dissecting phenotypic traits linked to human resilience to Alzheimer’s pathology. Brain 136, 2510–2526. 10.1093/BRAIN/AWT171.

135. Sánchez-Juan, P., Valeriano-Lorenzo, E., Ruiz-González, A., Pastor, A.B., Rodrigo Lara, H., López-González, F., Zea-Sevilla, M.A., Valentí, M., Frades, B., Ruiz, P., et al. (2024). Serum GFAP levels correlate with astrocyte reactivity, post-mortem brain atrophy and neurofibrillary tangles. Brain 147, 1667–1679. 10.1093/brain/awae035.

136. Strouhalova, K., Přechová, M., Gandalovičová, A., Brábek, J., Gregor, M., and Rosel, D. (2020). Vimentin Intermediate Filaments as Potential Target for Cancer Treatment. Cancers 2020, Vol. 12, 12. 10.3390/CANCERS12010184.

137. Wu, K., Zeng, J., Li, L., Fan, J., Zhang, D., Xue, Y., Zhu, G., Yang, L., Wang, X., and He, D. (2010). Silibinin reverses epithelial-to-mesenchymal transition in metastatic prostate cancer cells by targeting transcription factors. Oncol. Rep. 23, 1545–1552. 10.3892/OR_00000794/HTML.

138. Kim, Y.J., Choi, W. Il, Jeon, B.N., Choi, K.C., Kim, K., Kim, T.J., Ham, J., Jang, H.J., Kang, K.S., and Ko, H. (2014). Stereospecific effects of ginsenoside 20-Rg3 inhibits TGF-β1-induced epithelial–mesenchymal transition and suppresses lung cancer migration, invasion and anoikis resistance. Toxicology 322, 23–33. 10.1016/J.TOX.2014.04.002.

139. Martínez-Peña, F., Pearson, A.D., Tang, E.L., Kuburich, N.A., Mani, S.A., Schultz, P.G., Bollong, M.J., and Lairson, L.L. (2022). Synthesis and biological evaluation of novel FiVe1 derivatives as potent and selective agents for the treatment of mesenchymal cancers. Eur. J. Med. Chem. 242, 114638. 10.1016/J.EJMECH.2022.114638.

140. Eichenbaum, H. (2004). Hippocampus: Cognitive processes and neural representations that underlie declarative memory. Neuron 44, 109–120. 10.1016/J.NEURON.2004.08.028/ASSET/A0379ACF-1838-423B-B67A-561ADB848321/MAIN.ASSETS/GR2.GIF.

141. Selkoe, D.J. (2002). Alzheimer’s Disease Is a Synaptic Failure. Science (1979). 298, 789–791. 10.1126/SCIENCE.1074069.

142. Batiuk, M.Y., Martirosyan, A., Wahis, J., de Vin, F., Marneffe, C., Kusserow, C., Koeppen, J., Viana, J.F., Oliveira, J.F., Voet, T., et al. (2020). Identification of region-specific astrocyte subtypes at single cell resolution. Nat. Commun. 11, 1220. 10.1038/s41467-019-14198-8.

143. Kálmán, M. (2025). The relative withdrawal of GFAP-An essential component of brain evolution. Front. Neuroanat. 19, 1607603. 10.3389/fnana.2025.1607603.

144. Rossi, S., and Cozzolino, M. (2021). Dysfunction of RNA/RNA-Binding Proteins in ALS Astrocytes and Microglia. Cells 10. 10.3390/cells10113005.

145. Miedema, S.S.M., Mol, M.O., Koopmans, F.T.W., Hondius, D.C., van Nierop, P., Menden, K., de Veij Mestdagh, C.F., van Rooij, J., Ganz, A.B., Paliukhovich, I., et al. (2022). Distinct cell type-specific protein signatures in GRN and MAPT genetic subtypes of frontotemporal dementia. Acta Neuropathol. Commun. 10, 100. 10.1186/s40478-022-01387-8.

146. Minkeviciene, R., Rheims, S., Dobszay, M.B., Zilberter, M., Hartikainen, J., Fülöp, L., Penke, B., Zilberter, Y., Harkany, T., Pitkänen, A., et al. (2009). Amyloid β-Induced Neuronal Hyperexcitability Triggers Progressive Epilepsy. Journal of Neuroscience 29, 3453–3462. 10.1523/JNEUROSCI.5215-08.2009.

147. Deacon, R.M.J. (2006). Assessing nest building in mice. Nat. Protoc. 1, 1117–1119. 10.1038/nprot.2006.170.

148. Gonzalez-Lozano, M.A., Koopmans, F., Sullivan, P.F., Protze, J., Krause, G., Verhage, M., Li, K.W., Liu, F., and Smit, A.B. (2025). Stitching the synapse: Cross-linking mass spectrometry into resolving synaptic protein interactions. Sci. Adv. 6, eaax5783. 10.1126/sciadv.aax5783.

149. Thanou, E., Koopmans, F., Pita-Illobre, D., Klaassen, R. V, Özer, B., Charalampopoulos, I., Smit, A.B., and Li, K.W. (2023). Suspension TRAPping filter (sTRAP) sample preparation for quantitative proteomics in the low microgram input range using a plasmid DNA micro-spin column: Analysis of the hippocampus from the 5xFAD Alzheimer’s Disease mouse model. Cells 12, 1242. 10.3390/cells12091242.

150. Wiśniewski, J.R., and Gaugaz, F.Z. (2015). Fast and sensitive total protein and Peptide assays for proteomic analysis. Anal. Biochem. 87, 4110–4116. 10.1021/ac504689z.

151. Skowronek, P., Thielert, M., Voytik, E., Tanzer, M.C., Hansen, F.M., Willems, S., Karayel, O., Brunner, A.D., Meier, F., and Mann, M. (2022). Rapid and in-depth coverage of the (phospho-)proteome with deep libraries and optimal window design for dia-PASEF. Molecular & Cellular Proteomics 21, 100279. 10.1016/j.mcpro.2022.100279.

152. Demichev, V., Messner, C.B., Vernardis, S.I., Lilley, K.S., and Ralser, M. (2020). DIA-NN: neural networks and interference correction enable deep proteome coverage in high throughput. Nat. Methods 17, 41–44. 10.1038/s41592-019-0638-x.

153. Demichev, V., Szyrwiel, L., Yu, F., Teo, G.C., Rosenberger, G., Niewienda, A., Ludwig, D., Decker, J., Kaspar-Schoenefeld, S., Lilley, K.S., et al. (2022). dia-PASEF data analysis using FragPipe and DIA-NN for deep proteomics of low sample amounts. Nat. Commun. 13, 3944. 10.1038/s41467-022-31492-0.

154. Koopmans, F. (2024). GOAT: efficient and robust identification of gene set enrichment. Communications Biology 2024 7:1 *7*, 744-. 10.1038/s42003-024-06454-5.

155. Koopmans, F., van Nierop, P., Andres-Alonso, M., Byrnes, A., Cijsouw, T., Coba, M.P., Cornelisse, L.N., Farrell, R.J., Goldschmidt, H.L., Howrigan, D.P., et al. (2019). SynGO: an evidence-based, expert-curated knowledge base for the synapse. Neuron 103, 217–234.e214. 10.1016/j.neuron.2019.05.002.

156. Raudvere, U., Kolberg, L., Kuzmin, I., Arak, T., Adler, P., Peterson, H., and Vilo, J. (2019). g:Profiler: a web server for functional enrichment analysis and conversions of gene lists (2019 update). Nucleic Acids Res. 47, W191–W198. 10.1093/nar/gkz369.

157. Ran, F.A., Cong, L., Yan, W.X., Scott, D.A., Gootenberg, J.S., Kriz, A.J., Zetsche, B., Shalem, O., Wu, X., Makarova, K.S., et al. (2015). In vivo genome editing using Staphylococcus aureus Cas9. Nature 520, 186–191. 10.1038/nature14299.

