## Supplemental Figures for "Deletion of astrocyte intermediate filaments GFAP and Vimentin enhances protein synthesis and prevents early synaptic and cognitive dysfunction in a mouse model of Alzheimer’s disease"

**SUPPLEMENTARY FIGURE 1.**

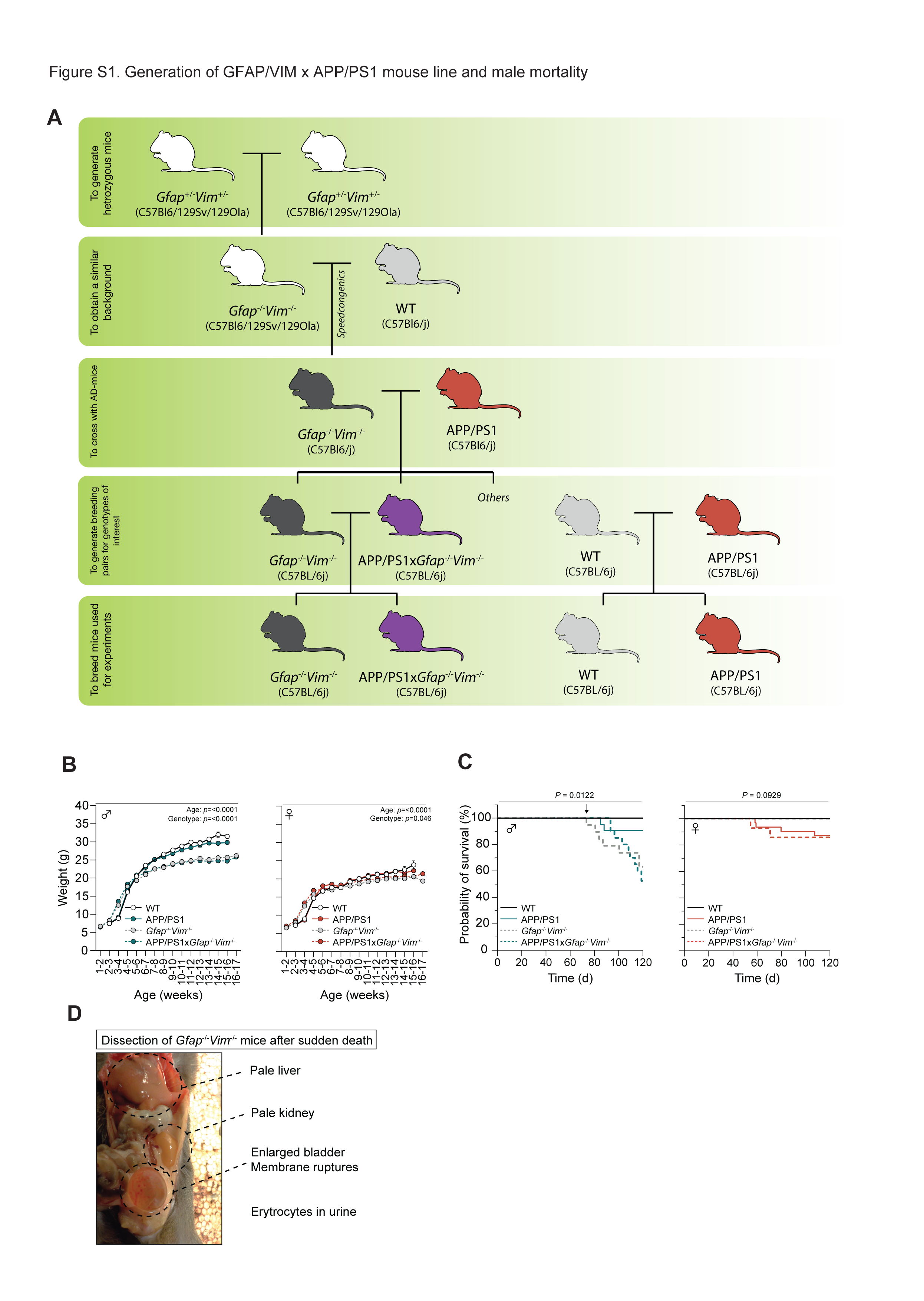

**Figure S1. Generation of APP/PS1xGfap^-/-^Vim^-/-^ and enhanced mortality in male, but not female, *Gfap*^-/-^*Vim*^-/-^ and APP/PS1x*Gfap*^-/-^*Vim*^-/-^ mice. A)** Schematic overview of the breeding scheme used to generate *Gfap*^-/-^*Vim*^-/-^ and APP/PS1x*Gfap*^-/-^*Vim*^-/-^ mice on a C57BL/6j genetic background. **B)** Average body weight measured once a week for male mice (**♂,** left) and female mice (**♀,** right). **C)** Probability of survival plot up to ~4 months of age for male (left) and female (right). The arrow in the left image is pointing towards the start of sudden non-AD related death in male *Gfap*^-/-^*Vim*^-/-^ and APP/PS1x*Gfap*^-/-^*Vim*^-/-^ mice. **D)** Representative photograph of a Gfap^-/-^Vim^-/-^ dissected abdomen after sudden death. The liver and kidneys are pale in color. The bladder is enlarged with small ruptures in the membrane. Erythrocytes are present in collected urine. Notice that in this photograph the liver is flipped towards the head to make the kidney visible. Statistical analysis performed with repeated measures ANOVA (B), Mantel-Cox (C). n = 5-33 mice per genotype. A = Age; Gen = Genotype. Data are presented as mean ± SEM. See table S3 for full statistical details.

**SUPPLEMENTARY FIGURE 2.**

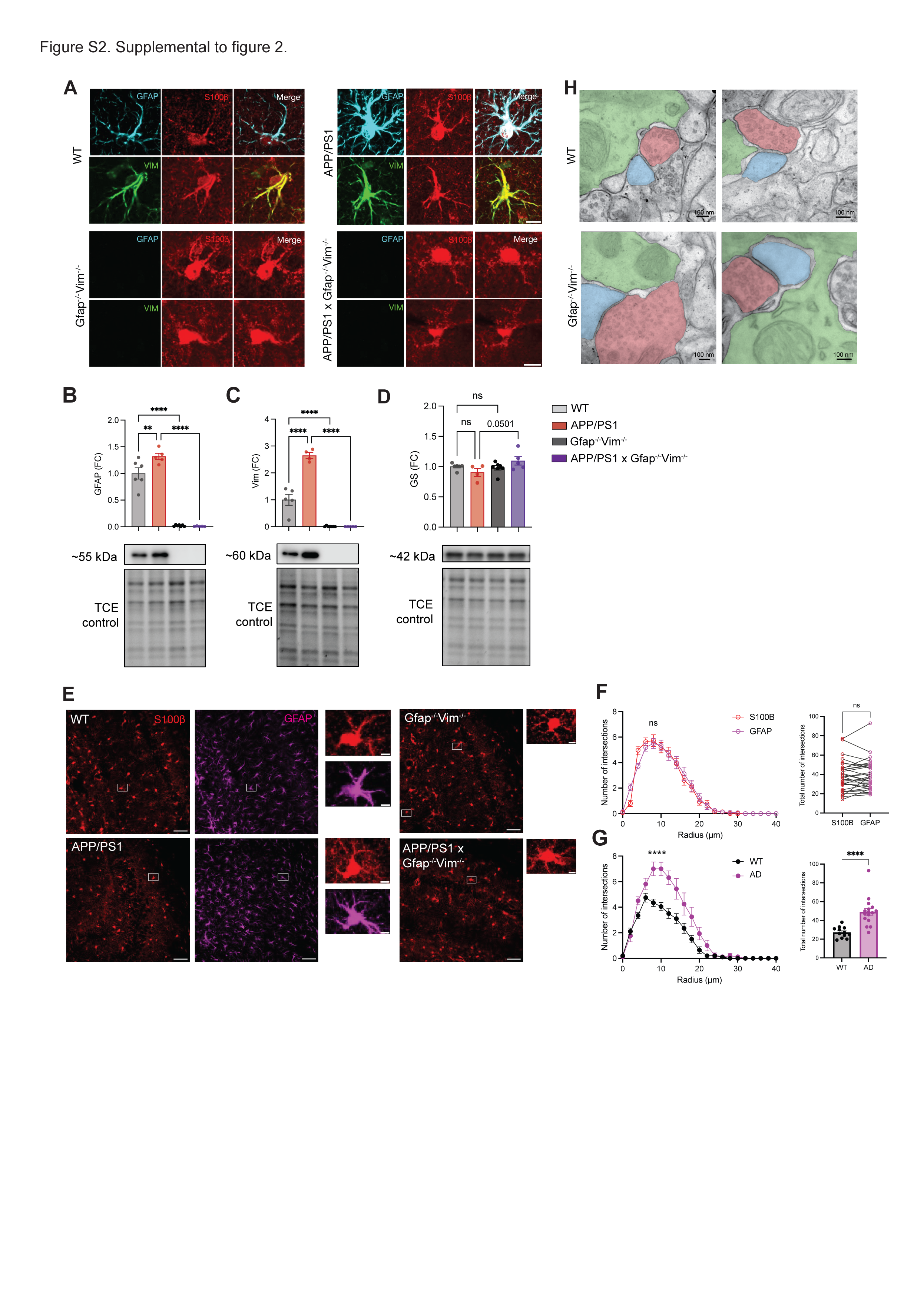

**Figure S2. Characterization of astrogliosis in APP/PS1 x Gfap^-/-^Vim^-/-^ mice. A)** Representative images of individual astrocytes shown in Fig. 2A stained for GFAP (cyan), Vimentin (green) and S100β (red). Scale bar, 10 μm. **B-D)** Quantification of protein levels of GFAP (B), VIM (C), GS (D) using immunoblotting. Data in B-D is represented as fold change relative to WT animals, and the shown expression is corrected for protein loading by 2,2,2-trichloroethanol (TCE) staining of the blot. (B-E), n = 5-6 mice per genotype. Statistical analysis performed by one-way ANOVA. **E)** Representative images of astrocytes in Fig. 2G, showing side-by-side astrocytes stained for S100β (red) and GFAP (magenta). Scale bar, 50 μm. The inset shows a zoom-in on a single astrocyte. Scale bar, 5 μm. **F)** Left, Sholl analysis profile displaying the mean number of intersections for the same astrocytes based on S100β or GFAP signal. Statistical significance was calculated by two-way repeated-measures ANOVA (Genotype: p = ns). Right, quantification of the total number of intersections from the Sholl analysis. Statistical significance calculated by paired t-test. n = 16-20 astrocytes from 3 mice per genotype. **G)** Left, Sholl analysis profile displaying the mean number of intersections for WT vs APP/PS1 astrocytes based on GFAP signal. Statistical significance was calculated by two-way repeated-measures ANOVA (Genotype: p < 0,0001). Right, quantification of the total number of intersections from the Sholl analysis. Statistical significance calculated by unpaired t-test. n = 16-20 astrocytes from 3 mice per genotype. **H)** Electron microscopy images from the hippocampal CA1 of two examples of perisynaptic astrocyte processes contacting a synapse in WT and *Gfap^-/-^Vim^-/-^* mice. Note that the ultrastructure of the tripartite synapse is preserved in *Gfap^-/-^Vim^-/-^* mice. Green, astrocyte; red, presynaptic terminal; blue, postsynapse. Scale bar, 100 nm. See table S3 for full statistical details. **p<0.01, ****p<0.0001, ns: not significant. Data are presented as mean ± SEM.

**SUPPLEMENTARY FIGURE 3.**

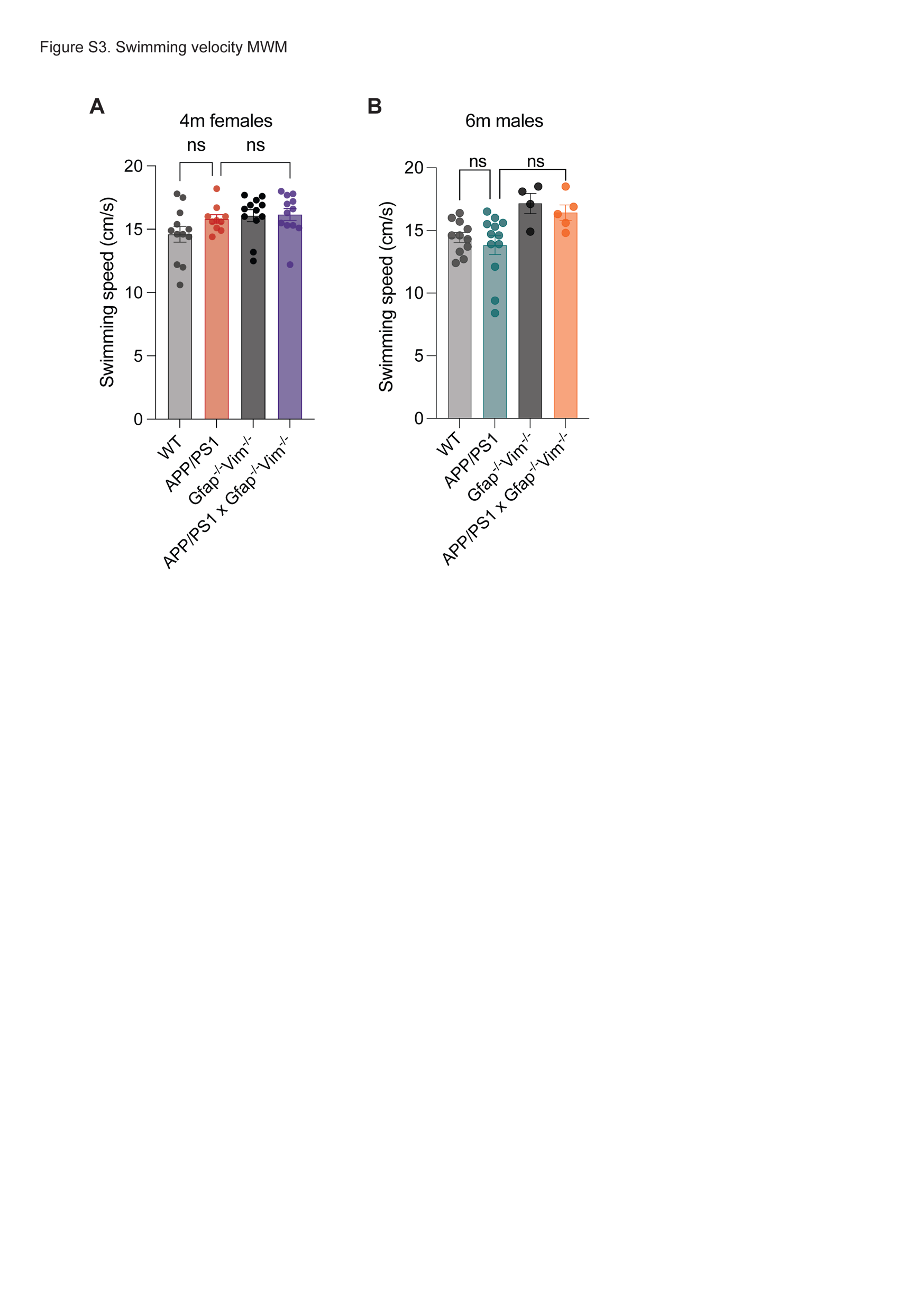

**Figure S3. Swimming speed in the Morris water maze test is not affected in APP/PS1 mice nor by deletion of GFAP and Vimentin. A)** Swimming speed for 4-month female mice. *n* = 10-12 per genotype. **B)** Swimming speed for 6-month male mice. n = 4-12 per genotype. Statistical analysis performed by Kruskal-Wallis test. See table S3 for full statistical details. #p<0.10; ns: not significant. Data are presented as mean ± SEM.

**SUPPLEMENTARY FIGURE 4.**

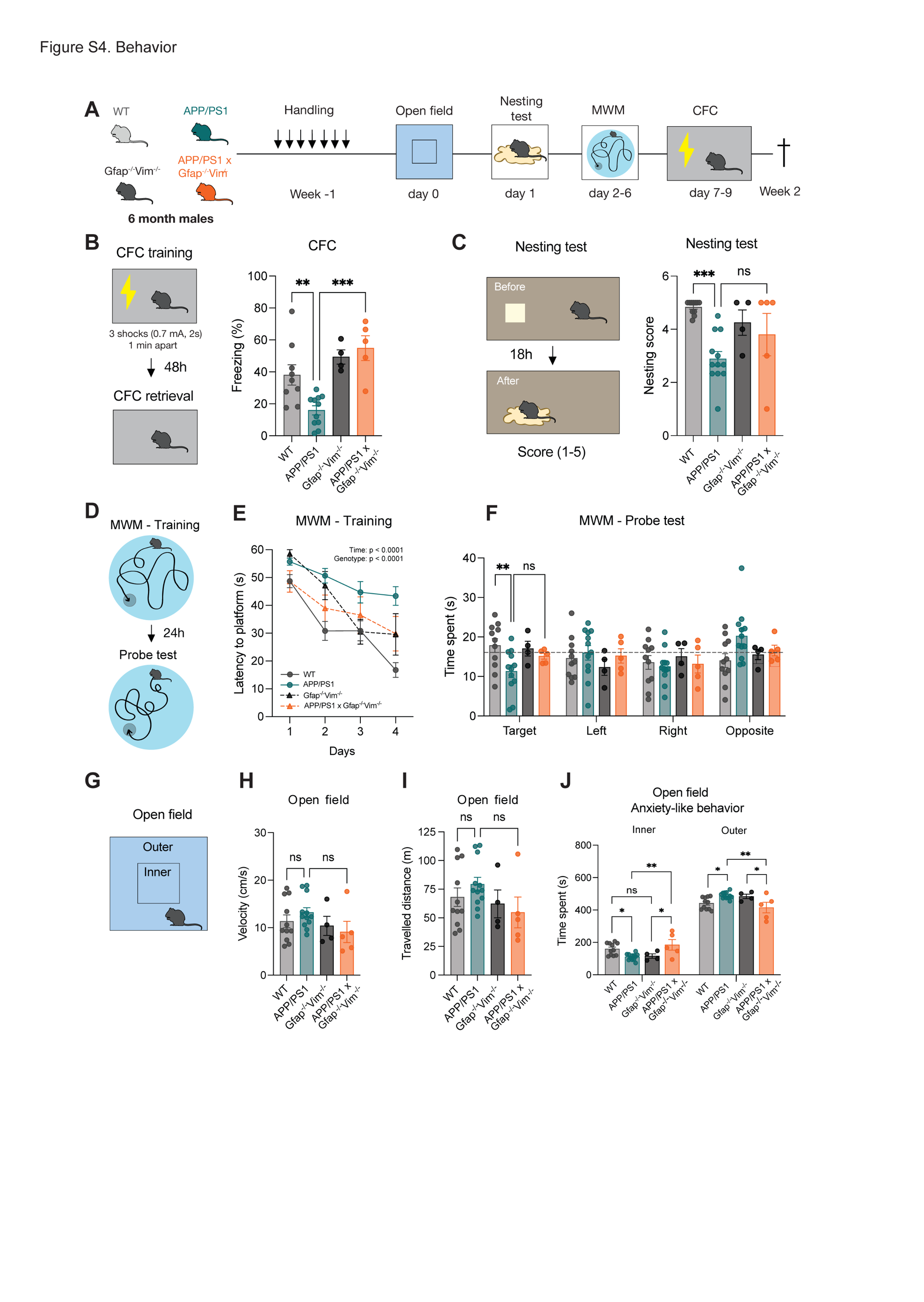

**Fig S4. Reduction of astrogliosis improves cognition in 6-month-old male APP/PS1 mice. A)** Schematic overview of the behavioral experiments. **B)** Left, experimental scheme of the contextual fear conditioning (CFC) training and retrieval. Right, freezing levels (%) measured during the CFC retrieval session 48 hours after shock. **C)** Left, experimental diagram of the nesting test. Right, nest-building scores assessed during the nesting test. **D)** Experimental scheme of the Morris water maze (MWM) training and probe test. **E)** MWM training latencies (in seconds) to reach the platform measured over four training days. **F)** Time spent (in seconds) in the MWM quadrant during the probe test, where the target quadrant is the one with the original location of the platform. **G)** Representative image of the open field area and the inner and outer areas. **H, I)** Locomotor activity determined by the velocity (H) and travelled distance (I) during the open field experiment, irrespective of the zone. **J)** Time spent (in seconds) in the inner zone and outer area of the open field experiment. Kruskal-Wallis (B, C), Repeated measures ANOVA (I), One-way ANOVA (E-G, J). n = 4-12 mice per genotype. See table S3 for full statistical details. *p<0.05, **p<0.01, ****p<0.0001, ns: not significant. Data are presented as mean ± SEM.

**SUPPLEMENTARY FIGURE 5.**

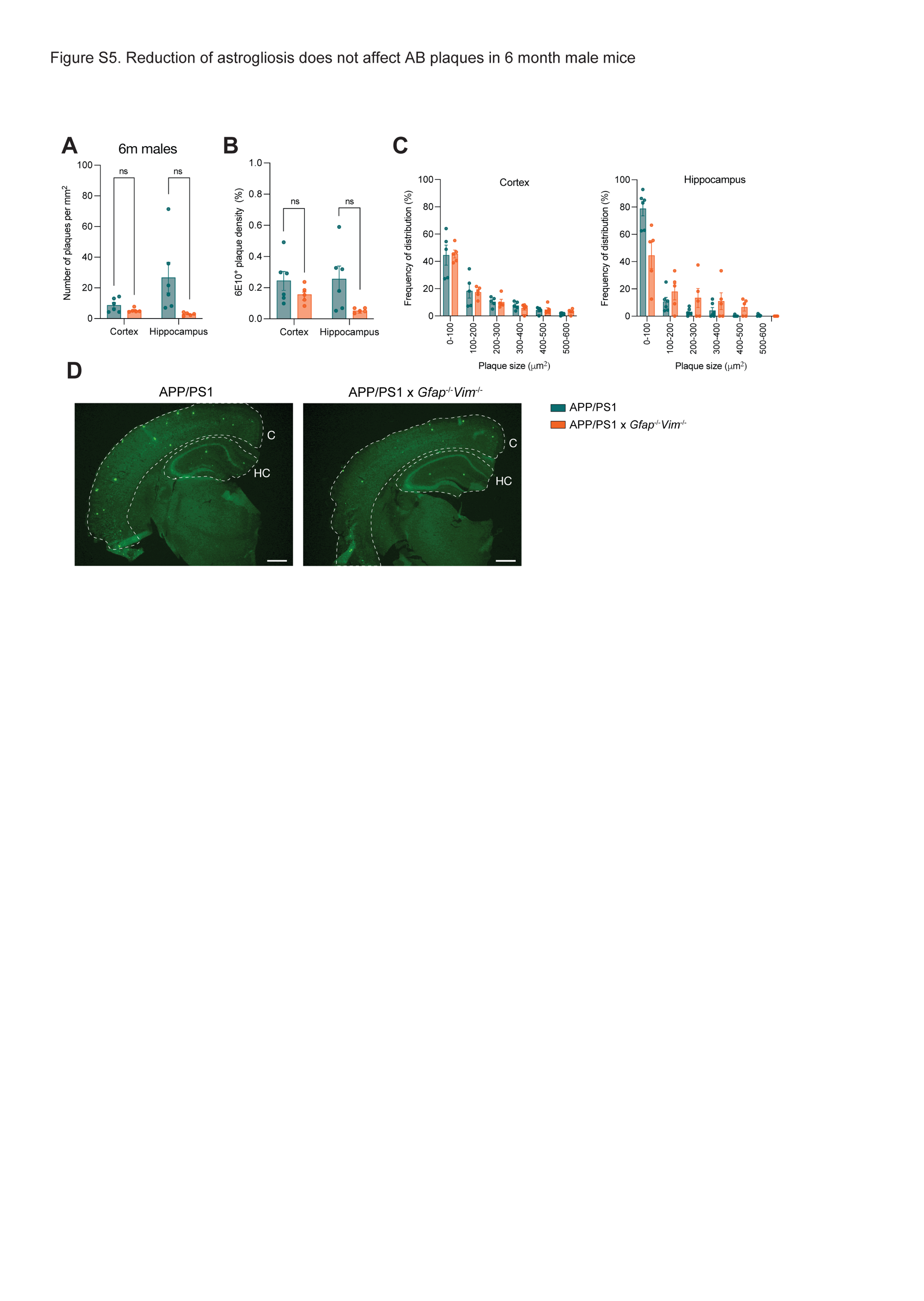

**Figure S5. Reduction of astrogliosis does not affect Aβ plaques deposition in 6-month-old APP/PS1 mice. A)** Quantification of plaque density in cortex and hippocampus, measured as the number of plaques per area of cortex or hippocampus in mm^2^ for 6-month male mice. **B)** Quantification of plaque density in cortex and hippocampus, measured as the total Aβ+ plaque area relative to total cortex or hippocampus area, respectively, for 6-month male mice. **C)** Frequency distribution analysis based on plaque size for cortex (left) and hippocampus (right). **D)** Representative images of Aβ immunohistochemistry. Top, full image of the brain left hemisphere for APP/PS1 mice (left) or APP/PS1xGfap^-/-^Vim^-/-^ mice (right). The dashed lines indicate either the cortex (C) or hippocampus (HC). Scale bar, 500 μm. Statistical analysis performed by unpaired t-test (A, B) or two-way ANOVA (C), n = 5-6 per genotype. See table S3 for full statistical details. ns, not significant. Data are presented as mean ± SEM.

**SUPPLEMENTARY FIGURE 6.**

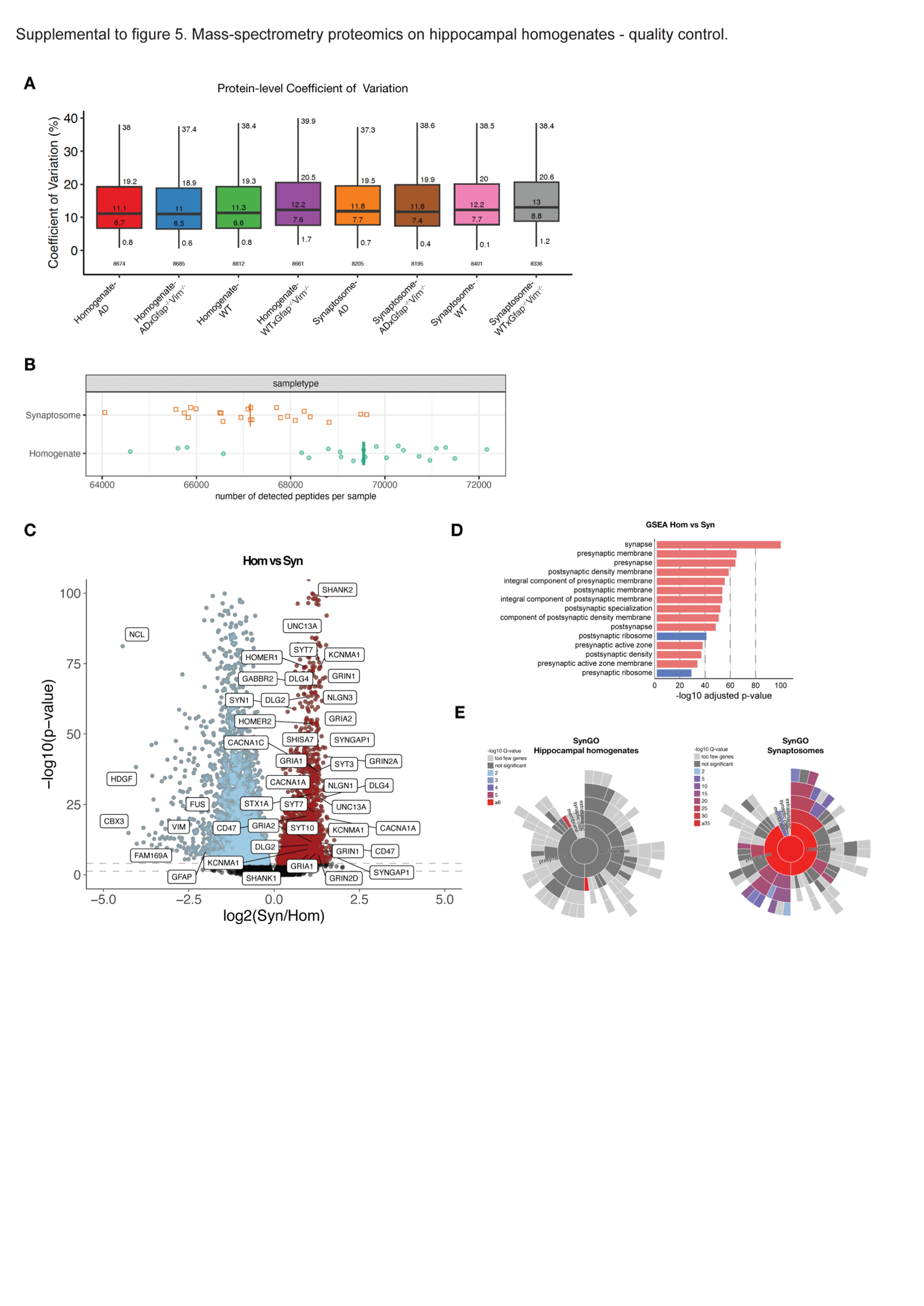

**Figure S6. Quality control of mass-spectrometry proteomics and of synaptic enrichment in synaptosomes. A)** Protein-level coefficient of variation for hippocampal homogenates and synaptosomes for the different genotypes demonstrates very high reproducibility. The number of detected proteins in at least 3 samples per group is noted below the boxplots. **B)** Number of detected peptides for the different samples in hippocampal homogenates and synaptosomes. Homogenates: n = 23 mice; synaptosomes, n = 23 mice. **C)** Volcano plot showing differential expression of proteins in hippocampal homogenates (Hom, n = 6 WT mice) vs synaptosomes (Syn, n = 6 WT mice). Note the upregulation of many synaptic proteins in synaptosomes. Lower dotted line at p<0.05; Upper dotted line at FDR<0.05. Complete DEP list in Table S1. DEPs upregulated in synaptosomes (red) and downregulated DEPs (blue). **D)** Overview of enriched pathways in downregulated (blue) or upregulated (red) DEPs in homogenates vs synaptosomes based on GSEA for GO terms. Adjusted p-values are FDR-corrected. **E)** Overrepresentation analysis of DEPs (p<0.05) in homogenates vs synaptosomes using SynGO showing synaptic enrichment for hippocampal synaptosomes but not homogenates.

**SUPPLEMENTARY FIGURE 7.**

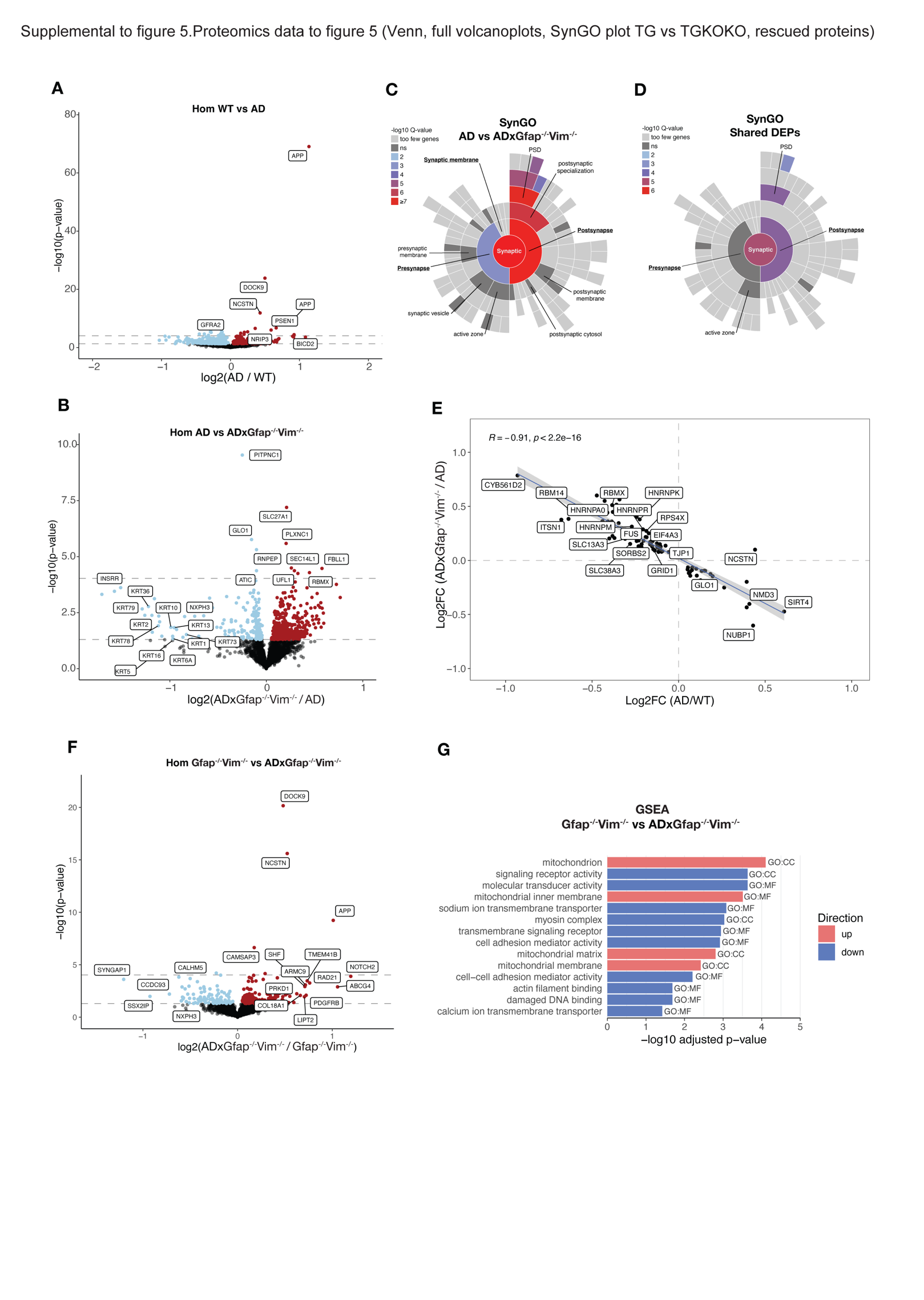

**Figure S7. Reduction of astrogliosis rescues the APP/PS1-mediated loss of synaptic proteins. A, B)** Complete volcano plots showing differential expression of proteins in WT vs APP/PS1 (A, WT: n = 6 mice; APP/PS1: n = 5 mice) or APP/PS1 vs APP/PS1xGfap^-/-^Vim^-/-^ (B, APP/PS1: n = 5 mice; APP/PS1xGfap^-/-^Vim^-/-^: n = 5 mice). Lower dotted line at p<0.05; Upper dotted line at FDR<0.05. Complete DEP list in Table S1. DEPs upregulated in APP/PS1 marked in red and downregulated DEPs marked in blue. **C)** Overrepresentation analysis of DEPs (p<0.05) in APP/PS1 vs APP/PS1xGfap^-/-^Vim^-/-^ using SynGO showing synaptic enrichment. **D)** Overrepresentation analysis of shared DEPs (p<0.05) between WT vs APP/PS1 and APP/PS1 vs APP/PS1xGfap^-/-^Vim^-/-^ using SynGO depicting synaptic enrichment. **E)** Correlation plot showing a strong inverse correlation for shared DEPs in WT vs APP/PS1 compared with APP/PS1 vs APP/PS1xGfap^-/-^Vim^-/-^. **F)** Complete volcano plot showing differential expression of proteins in Gfap^-/-^Vim^-/-^ vs APP/PS1xGfap^-/-^Vim^-/-^ (Gfap^-/-^Vim^-/^: n = 7 mice; APP/PS1xGfap^-/-^Vim^-/-^: n = 5 mice). **G)** Overview of enriched pathways in downregulated (blue) or upregulated (red) DEPs in Gfap^-/-^Vim^-/-^ vs APP/PS1xGfap^-/-^Vim^-/-^ based on GSEA for GO terms (CC, cellular component; BP, biological process; MF, molecular function). All displayed comparisons refer to hippocampal homogenates (Hom, homogenates). Adjusted p-values are FDR-corrected.

**SUPPLEMENTARY FIGURE 8.**

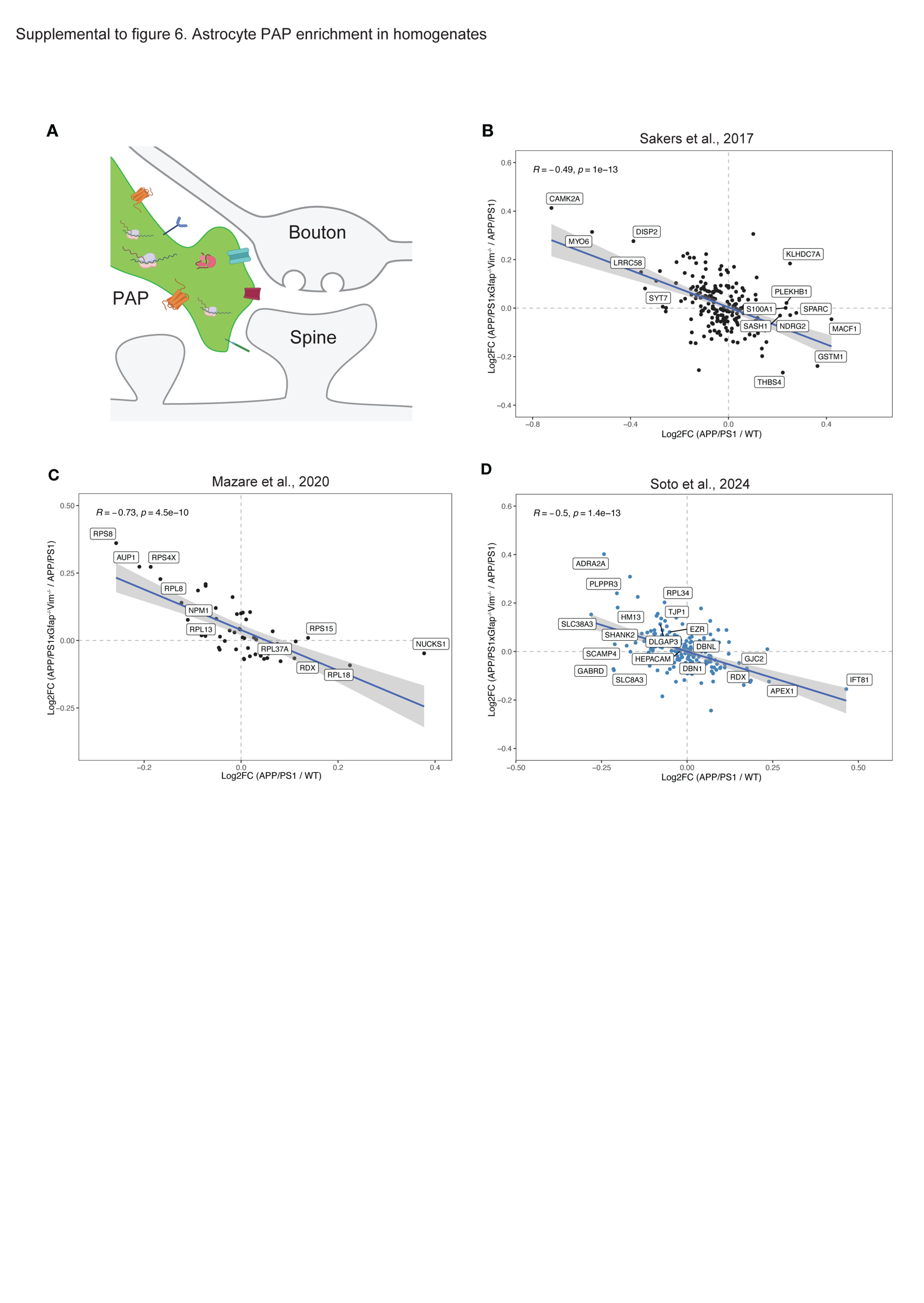

**Figure S8. Astrocytic PAP proteins are altered in APP/PS1 and restored by reduction of astrogliosis. A)** Cartoon depicting a PAP in contact with a synapse, with PAP-enriched transcripts and proteins highlighted. **B)** Correlation plot showing a significant inverse correlation in WT vs APP/PS1 compared with APP/PS1 vs APP/PS1x*Gfap^-/-^Vim^-/-^* for transcripts present in the PAPs as in Sakers et al., 2017^1^. **C)** Same as in (B), but for PAP-enriched transcripts as in Mazare et al., 2020^2^. **D)** Same as in (B) but for PAP-enriched proteins as in Soto et al., 2024^3^.

**SUPPLEMENTARY FIGURE 9.**

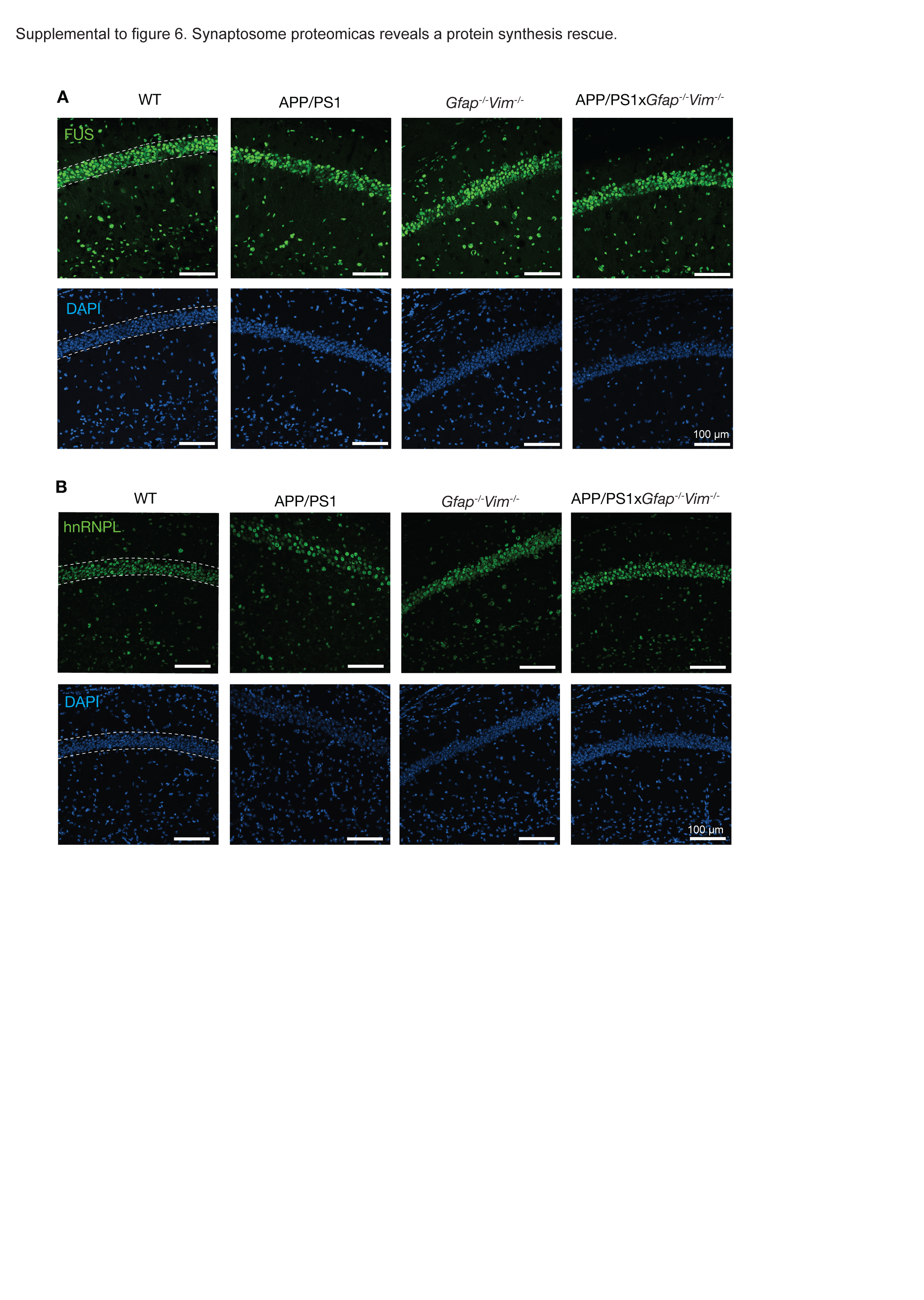

**Figure S9. Expression of RBPs FUS and hnRNPL is not changed in the hippocampal CA1 pyramidal layer in APP/PS1 mice, nor after deletion of GFAP and Vimentin.** **(A)** Representative images of FUS expression in CA1, with DAPI for nuclear staining and detection of the pyramidal layer (highlighted with a dashed line, for reference), for mice with depicted genotypes. **(B)** Same as in (A) but for hnRNPL. Scale bar, 100 μm.

**SUPPLEMENTARY FIGURE 10.**

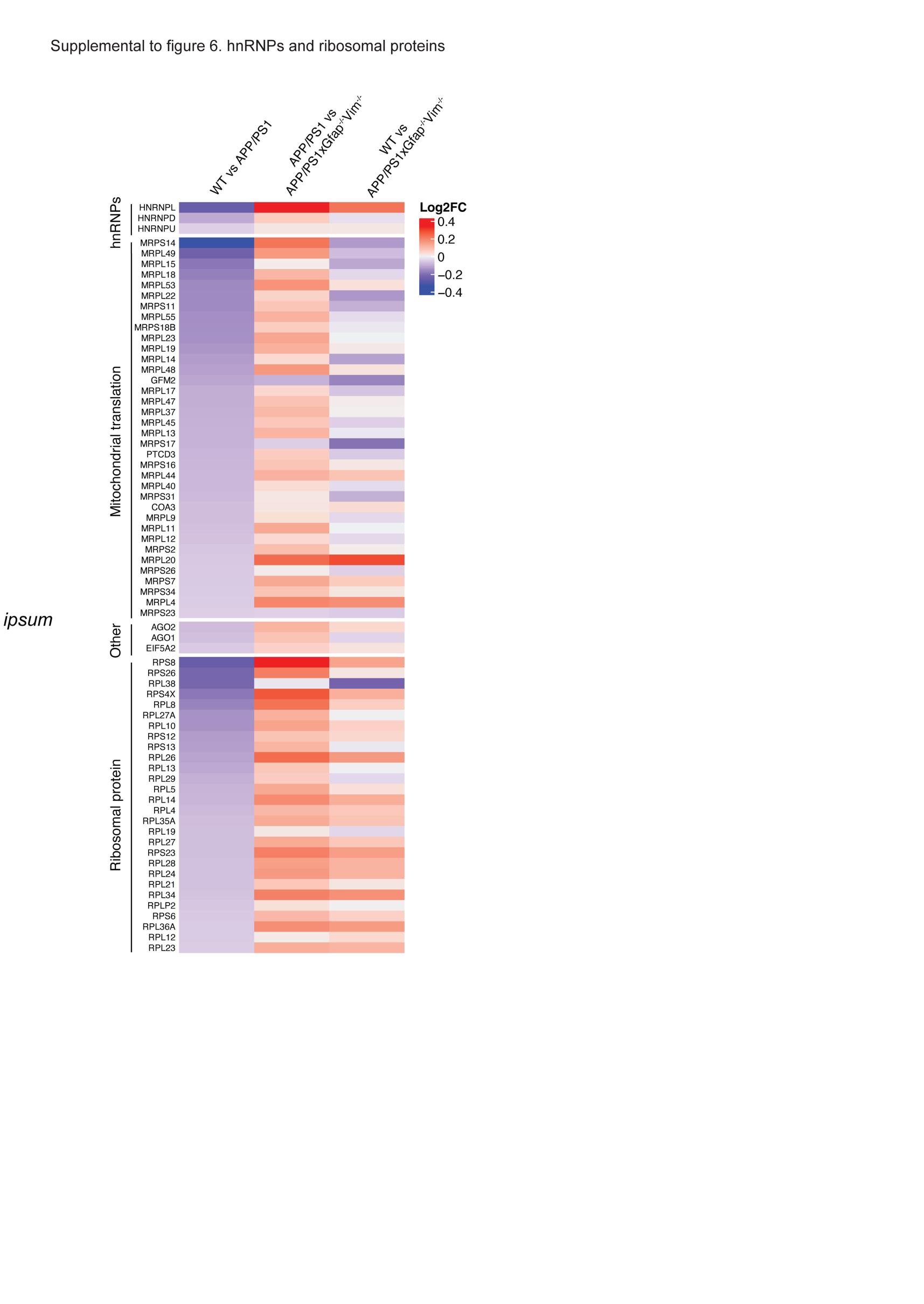

**Figure S10. Expression of proteins involved in translation.** Heatmap showing the expression changes (in Log2) for the top 75 downregulated proteins in hippocampal homogenates from WT vs APP/PS1 4-month female mice from the “translation” term in GO:0006412, annotated according to their function. Also shown are the expression changes for this set of proteins in APP/PS1 vs APP/PS1x*Gfap^-/-^Vim^-/-^* and WT vs APP/PS1x*Gfap^-/-^Vim^-/-^* 4-month female mice.

**SUPPLEMENTARY FIGURE 11.**

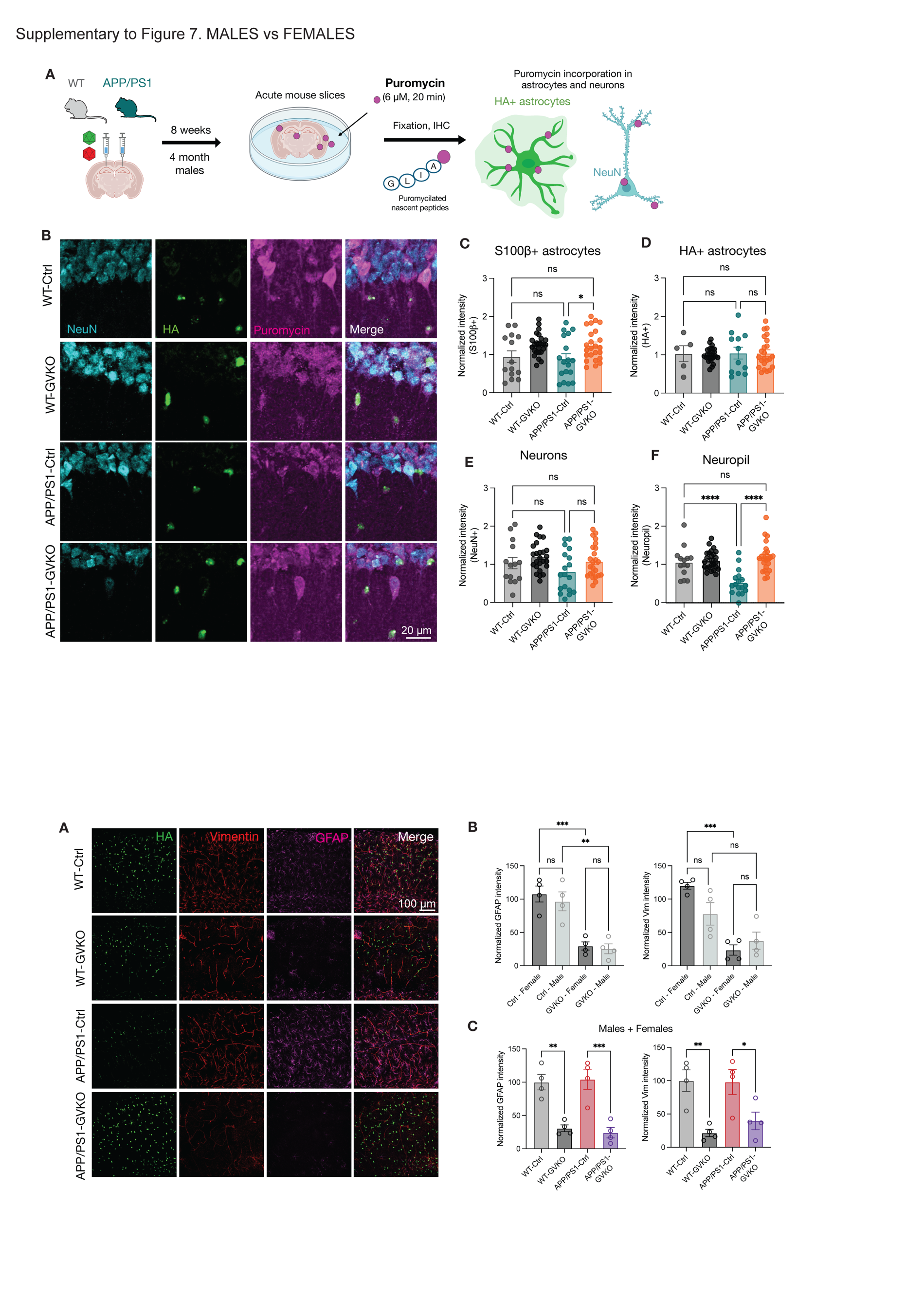

**Figure S11. CRISPR-Cas9 deletion of GFAP and Vimentin in astrocytes from male and female mice. A)** Representative images for Ctrl and GFAP/Vim gRNA WT and APP/PS1 mice stained for HA, GFAP and Vimentin. Scale bar, 100 μm. **B)** Quantification of GFAP (left) and Vimentin (right) protein levels per astrocyte (HA+S100β+) in Ctrl and GVKO male and female mice. **C)** Quantification of GFAP (left) or Vimentin (right) protein levels per astrocyte (HA+S100β+). 2092 astrocytes from 4 mice/group. Statistical significance calculated by one-way ANOVA. See table S3 for full statistical details. *p<0.05, **p<0.01, ***p<0.001, ns: not significant. Data are presented as mean ± SEM.

**SUPPLEMENTARY FIGURE 12.**

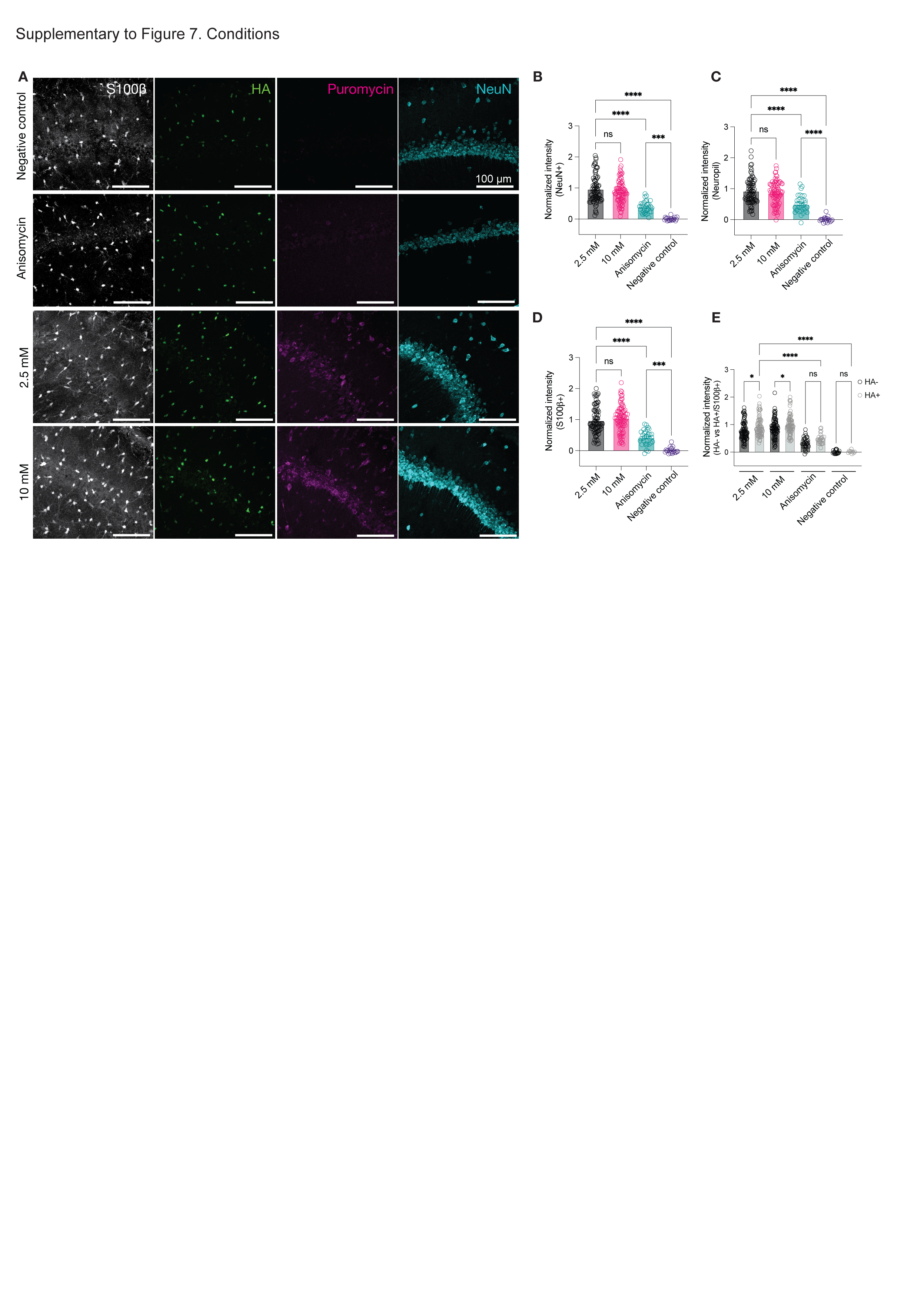

**Figure S12. Protein translation-dependent puromycin labelling of newly synthetized peptides in neurons and astrocytes.** Slices were treated with 6 μM puromycin for 20 min in aCSF with 2.5 or 10 mM K^+^ to increase neuronal activity, and with the translational inhibitor anisomycin (200 μM) for 10 min prior to puromycin treatment. No puromycin was added to the negative control. **A)** Representative images of the different conditions showing puromycin labelling in neurons (NeuN), astrocytes (S100β) and transduced astrocytes (HA/S100β). Scale bar, 100 μm. **B-E)** Quantification of the puromycin fluorescence intensity in the different in neurons (B), neuropil as a proxy for synaptic translation (C), astrocytes (D) and comparison of non-transduced (HA-) vs transduced (HA+) astrocytes (E). The negative control background was subtracted from all conditions. See table S3 for full statistical details. *p<0.05, ***p<0.001, ****p<0.0001, ns: not significant. Data are presented as mean ± SEM.

**SUPPLEMENTARY FIGURE 13**

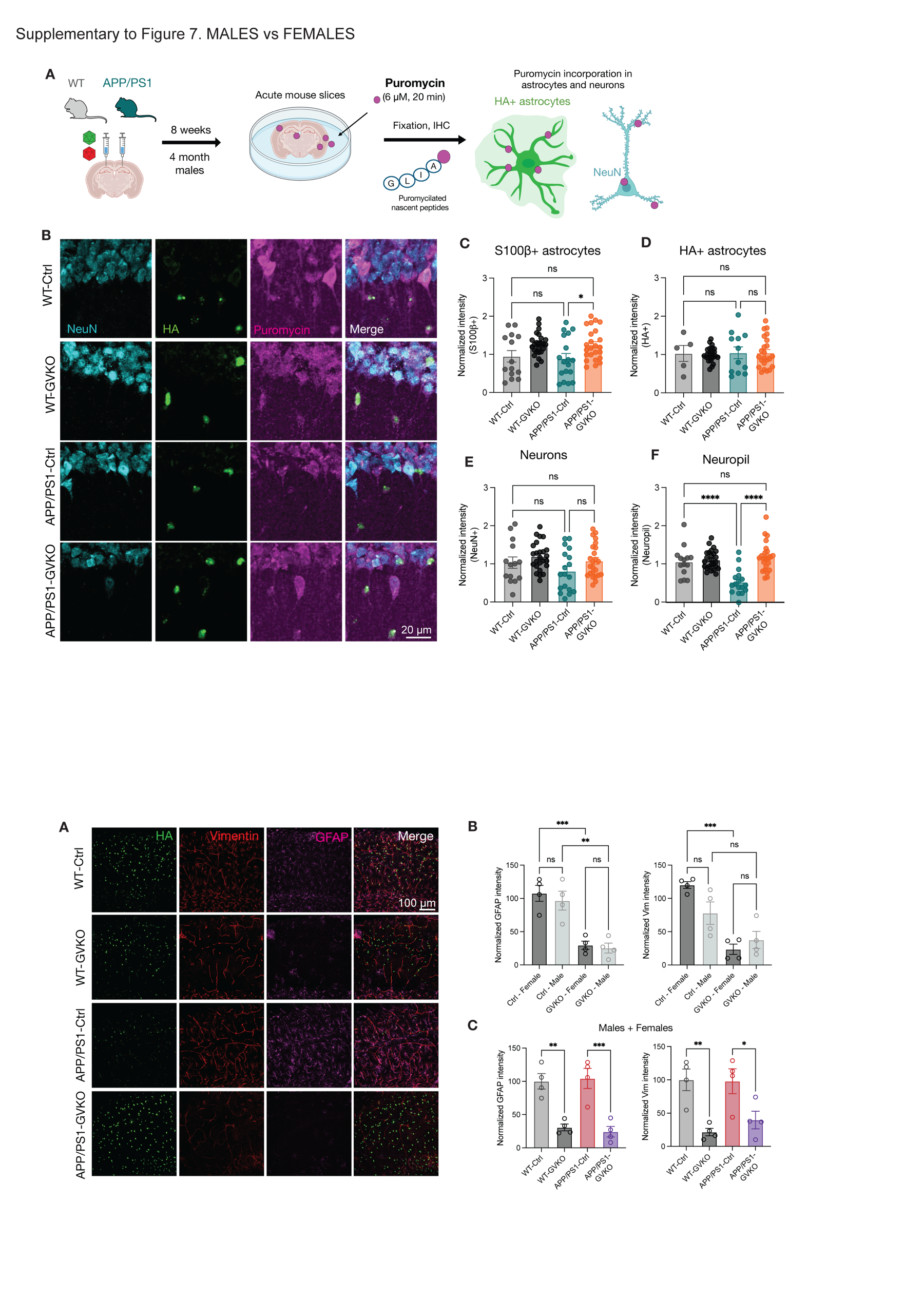

**Figure S13. Evaluation of the effects on protein synthesis of deleting astrocytic GFAP and Vimentin in male WT and APP/PS1 mice. A)** Schematic of SUnSET assay for *in vivo* puromycin labelling of newly synthetized peptides in hippocampal neurons and astrocytes using astrocyte-specific CRISPR-Cas9 in 4-month male WT and APP/PS1 mice. **B)** Representative images showing puromycin incorporation (magenta) in neurons (NeuN, cyan) and astrocytes (S100β, green). Scale bar, 20 μm. **C-F)** Normalized puromycin incorporation in all astrocytes (C), transduced astrocytes (D), neurons (E) and neuropil (F). n = 14-25 slices from 4-5 mice/group. Statistical significance calculated by one-way ANOVA. See Table S3 for full statistical details *p<0.05, ****p<0.0001, ns: not significant. Data are presented as mean ± SEM.

**Methods S1. MS-DAP file.**

**Table S1. Differential abundance analysis homogenates/synaptosomes**

**Table S2. Full results from GOAT.**

**Table S3. Statistical details.**

**Supplementary references**

1.	Sakers, K., Lake, A.M., Khazanchi, R., Ouwenga, R., Vasek, M.J., Dani, A., and Dougherty, J.D. (2017). Astrocytes locally translate transcripts in their peripheral processes. Proceedings of the National Academy of Sciences *114*, E3830–E3838. https://doi.org/10.1073/pnas.1617782114.

2.	Mazaré, N., Oudart, M., Moulard, J., Cheung, G., Tortuyaux, R., Mailly, P., Mazaud, D., Bemelmans, A.-P., Boulay, A.-C., Blugeon, C., et al. (2020). Local Translation in Perisynaptic Astrocytic Processes Is Specific and Changes after Fear Conditioning. Cell Rep *32*, 108076. https://doi.org/https://doi.org/10.1016/j.celrep.2020.108076.

3.	Soto, J.S., Jami-Alahmadi, Y., Chacon, J., Moye, S.L., Diaz-Castro, B., Wohlschlegel, J.A., and Khakh, B.S. (2023). Astrocyte–neuron subproteomes and obsessive–compulsive disorder mechanisms. Nature *616*, 764–773. https://doi.org/10.1038/s41586-023-05927-7.
